# Microphysiological Flow Batteries For Dynamic EDC Screening Of mESC-derived Thyroid Organoids

**DOI:** 10.64898/2026.05.07.722520

**Authors:** Anna M. Kip, Daniel J. Carvalho, Marta Nazzari, Mírian Romitti, James Waddington, Carlotta Branca, Conor Bryan, Barry Jutten, Whitney van de Vin, Prakash Patel, Rikke Poulsen, Martin Hansen, Simon Thomas, Stephen R. Pennington, Sabine Costagliola, Florian Caiment, Stefan Giselbrecht, Lorenzo Moroni

## Abstract

Endocrine disrupting chemicals (EDCs) are ubiquitous environmental contaminants capable of dysregulating the production of thyroid hormones. Traditional thyroid toxicological assays rely on 2D cell cultures and animal models, both of which fail to accurately recapitulate human thyroid physiology and provide limited mechanistic insight into EDC toxicity. To overcome these limitations, we report a novel *thyroid-on-chip* platform integrating mouse embryonic stem cell–derived thyroid organoids with advanced organ-on-chip (OoC) technology and downstream multi-omics analysis. The platform leverages a reversibly-sealed microphysiological flow battery (MFB) to allow scale up of dynamic organoid culture and controlled chemical exposure while reducing operational complexity. Upon EDC exposure, transcriptomic and proteomic analysis revealed new molecular signatures of thyroid disruption across four different EDC classes, even at very low EDC concentrations (1nM), validating the capacity of this system to mechanistically dissect EDC-induced responses. This represents an integrated platform consists of an advanced physiologically relevant assay framework for next-generation endocrine toxicity testing, bridging the gap between *in vitro* screening and *in vivo* thyroid physiology.

## 1. Introduction

Humans and wildlife are exposed daily to a multitude of synthetic chemicals, some of which can adversely interfere with the endocrine system^1^. These endocrine disrupting chemicals (EDCs) can dysregulate thyroid hormone production, which is a major biochemical regulator of crucial physiological processes, including growth, development, and metabolism. As a result, exposure of the thyroid gland to EDCs has been associated with the increased prevalence of several chronical diseases, such as reproductive disorders, obesity, and hormone-sensitive cancers^2,3^. Since the early 1990’s, governments and regulatory agencies around the world have made significant efforts to raise awareness, identify EDCs and to control their continuous release into the environment^4^. However, current assays of EDC thyroid toxicity are still based on animal models and *in vitro* cell-based cultures, which often fail to accurately identify EDCs^5,6^.

Thyroid cell-based assays are widely used and normally comprise mammalian cell lines cultured on microtiter plates for interrogation of the chemical effects, for example on iodide uptake by sodium iodide symporter (Nis)^7^ and on the dysregulation of thyroperoxidase (Tpo)^8^. Despite offering high-throughput capabilities, these methodologies do not capture the 3D structure and functionality of the thyroid, hence fail to reproduce human responses to EDC exposure^4^. In addition, existing assays lack capability to decipher fully the intricate molecular events triggered by EDC exposure due to their incompatibility with mechanistic analytical tools. Consequently, the mechanisms and mode of actions associated with EDC toxicity remain poorly understood^1^. Therefore, the development of new platforms that properly emulate the functionality of the thyroid gland while being amenable to mechanistic analysis by, for example omics technologies^9,10^, is urgently needed.

In recent years, stem-cell derived organoids have emerged as promising 3D multicellular models that can replicate the complex anatomy and functionality of various tissues *in vitro*, including the thyroid^11^. Owing to their improved physiological relevance, thyroid organoid models differentiated from human and mouse pluripotent stem cells have been shown to outperform 2D culture assays in reproducing thyroid physiological and pathological conditions^12–14^. However, the high fidelity of such 3D models comes at the expense of its analytical power and control over culture parameters^15,16^. To overcome these limitations, organoids have recently been integrated into microfluidic devices, such as organ-on-chips (OoC). These dynamic culture systems have demonstrated to enhance maturation and organization of organoids, while enabling parallelization and automation^15,17,18^. Therefore, OoC devices have the capacity to transform organoid culture into more informative assays for drug toxicity studies^19^. Although various OoC devices have been successfully combined with organoids for assessing drug efficacy and toxicity in tumour^20^, cardiac^21–23^, and lung^24^, few have been reported in thyroid^25,26^. In addition, the integration of multi-omics analysis and the execution of large chemical screenings on-chip has proven challenging due to the lack of scalability and the inefficient cell culture protocols of traditional chip designs^23,27^.

To mitigate these limitations, we have developed a customized thyroid-on-chip platform for advancing EDC screening by combining mouse embryonic stem cell (mESC)-derived thyroid organoids, OoC technology, and multi-omics analysis (**Figure 1a**). More specifically, by using reversibly-sealed OoC devices, we engineered a microphysiological flow battery (MFB) to significantly decrease operation complexity and (dis-)assembly times, which allowed a multiplexed on-chip screening of thyroid organoids exposed to panels of EDCs under controlled fluid dynamic conditions. EDC exposures were followed by thyroid hormone quantification and detailed differential expression and functional enrichment pathway analysis of the biological effects at the gene and protein levels. Finally, integrated omics studies unraveled new signatures of EDC-mediated toxicity and demonstrated the ability of our platform to mechanistically understand EDC exposure.

**Figure 1.**
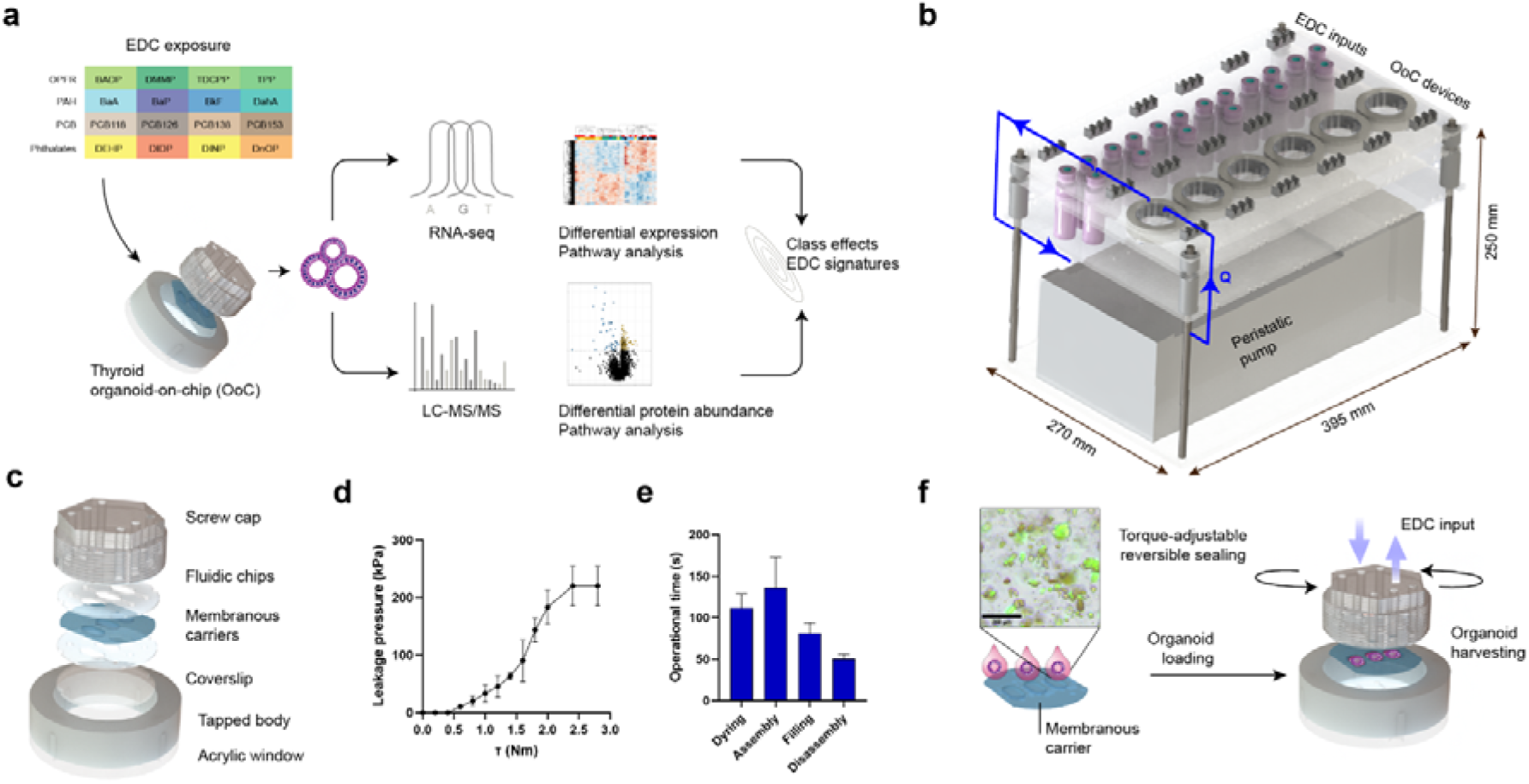
Microphysiological flow battery (MFB) for EDC screening of perfused mESC-derived thyroid organoids. **(a)** Schematics of the experimental pipeline of the EDC screening on thyroid OoC devices. Thyroid organoids were exposed to a panel of *sixteen* EDCs belonging to the four different classes*: OPFR, PAH, PCB, phthalates*. These EDCs were perfused for 24h in the thyroid OoC devices of the MFBs and their effects were assessed by transcriptomics and proteomics. OPFR, organophosphate flame-retardants; PAH, polycyclic aromatic hydrocarbons; PCB, polychlorinated biphenyls. **(b)** 3D CAD model of the MFB. The platform comprises 54x cell culture medium reservoirs loaded with multiple EDCs; six OoC devices, and a multichannel peristaltic pump. The blue circuit represents the fluidic interconnection via tubing between all components. The arrow indicates the direction of flow. **(c)** 3D CAD explosion view of the OoC device and its components. The OoC device comprises a polycarbonate housing and a coverslip (grey), a PMMA window (transparent), two PDMS fluidic chips (white), and a 50 µm thick membranous polycarbonate carrier (blue). **(d)** Quantification of the maximum internal air pressure (kPa) that the OoC device can withstand for different applied torque values (τ, Nm). **(e)** Times of the different operations involving assembling and disassembling of OoC devices. First, membranous carriers are removed from culture wells and dried, followed by integration on the OoC device (i.e. assembly). Subsequently, channels are filled with cell culture medium. After EDC exposure, cells are harvested by opening the OoC device (i.e. disassembly). **(f)** Schematic illustration of the culture operations using the OoC device. The Lock-and-play (LnP) sealing enables easy (un-)loading of thyroid organoids in/out the OoC device. After EDC exposure, organoids were collected and the EDC effects were analysed by transcriptomics and proteomics. Merged brightfield image with fluorescence signal (GFP) of mESC-derived thyroid follicles pre-matured for three days in membranous carriers. Scale bar, 250 µm.

## 2. Results

### 2.1. Design of the microphysiological flow batteries

We developed modular MFBs that enable screening of multiple EDCs on physiological relevant thyroid organoids under dynamic cell culture conditions (**Figure 1b**). Each battery consisted of a four-layered, acrylic manifold that hosted up to six OoC devices, 54 x 20 mL glass reservoirs, and a peristaltic pump. The reservoirs and OoC devices were positioned in such a way that all fluidic interfaces were aligned at the top level, while the pump was located below them (**Figure S1a**). This defined and reproducible configuration allowed to minimize tubing length required to connect each component in the circuit loop down to 107 cm, which represented a 60% drop in material when compared to our previous microfluidic set-ups^12^. Moreover, the system facilitated tubing handling and provided a maximum compactness of 270 x 395 x 250 mm (W x L x H), corresponding to a total volume of ∼ 25 L (**Figure 1b**). Up to three microphysiological flow batteries were placed inside a CO_2_ incubator with an active humidity control system (**Figure S1b and c**). For the tubing, a pharmaceutical grade material (PharMed^®^ BPT) was used, which enabled a reduction of compound ab-/adsorption in-/to the tubing walls, making it a preferred choice for chemical screening (**Figure S2a** and **b**).

The OoC device comprised a tapped body, where multiple fluidic compartments and a membranous carrier were vertically stacked and tightened against a polymethyl methacrylate (PMMA) bottom window by fastening a screw cap (**Figure 1c**). To ensure proper alignment between components, the OoC devices were assembled on a custom-made tool featuring two vertical alignment pins (**Figure S3a** and **S3b**). To facilitate the on-chip culture and harvesting of organoids for downstream analysis, all components of the OoC device were reversible sealed by a novel lock-and-play (LnP) clamping system^28^. In brief, the top part of the screw cap was hexagonally shaped to enable controlled tightening with a hex bit socket and an adjustable torque screwdriver. In this way, a pre-determined clamping load could be applied and maintained, also enabling external adjustment of the sealing strength at any time of the experiment and with minimal intervention to the internal fluidic culture. In calibration experiments, resulting sealing strengths were quantitively assessed against increasing torque (**Figure 1d**). The engineered LnP sealing showed no leakages and no cross-contamination between neighboring channels (**Figure S4a** and **S4b**). Additionally, the developed OoC device reduced dis-/assembly times, which is essential for increasing throughput in screening experiments (**Figure 1e**). The addition of a coverslip to the OoC device was crucial to prevent substantial displacement and deformation of the fluidic chips by decoupling the rotational motion at the interface between the acrylic bottom and the coverslip (**Figure S5a**).

Matrigel-embedded mESC-derived thyroid follicles were loaded into three thermoformed rectangular cell compartments of a membranous polycarbonate carrier (**Figure 1f**). The geometry of the cell compartments and channels was 1 mm in height. The size of the cell compartments were designed with footprints of 25 and 50 mm^2^ to ensure sufficient availability of cell material for transcriptomic and proteomic downstream analysis, respectively (**Figure S5b**). The cell chamber was set to 0.7 mm in depth. Thyroid organoid distribution and growth were monitored throughout seeding, pre-culture (outside MFB) and culture using the endogenous expression of bTg promoter-driven green fluorescent protein (GFP, green)^12^ (**Figure 1f**).

### 2.3. Screening of EDCs on thyroid OoC model

We aimed to investigate the effects of EDC exposure on perfused mESC-derived thyroid organoids. Perfused thyroid OoC devices were exposed to 16 different EDCs from four EDC classes at two concentrations (1 nM and 10 µM): polycyclic aromatic hydrocarbons (PAH: BaA, BaP, BkF, DahA), polychlorinated biphenyls (PCB118, PCB126, PCB138, PCB153), phthalates (DEHP, DIDP, DINP, DnOP), and organophosphate flame retardants (OPFR: BADP, DMMP, TDCPP, TPP). Exposures were performed in four consecutive screenings (one EDC class per run) at a flow rate of 12 µL/min (**Figure 2a**). In each screening, exposure to the reference compound Methimazol (MMI, 10 µM) and an untreated control (DMSO used as vehicle control) were included. After 24h exposure, organoids were collected for transcriptomics (1 nM and 10 µM) and proteomics analysis (10 µM).

**Figure 2.**
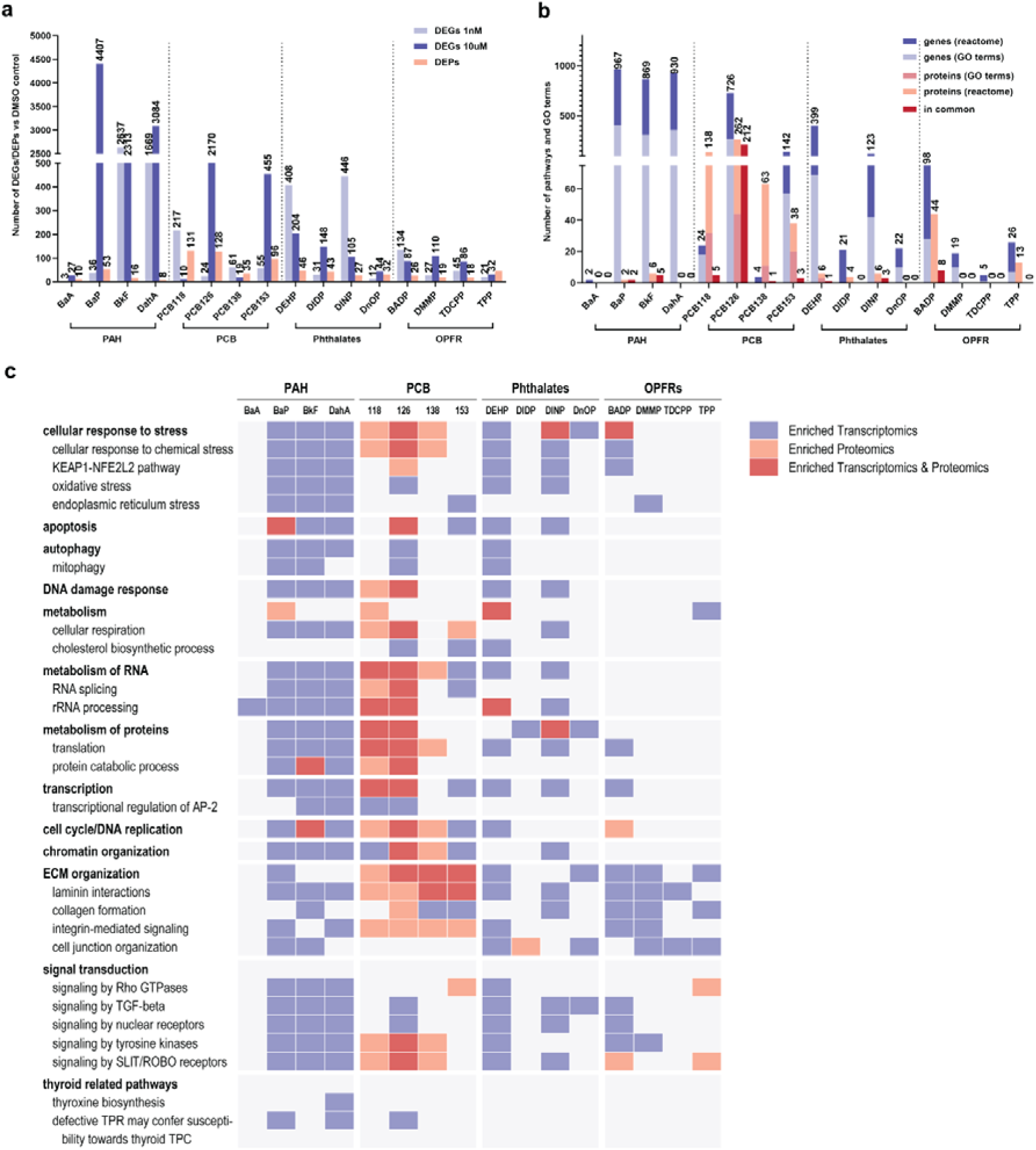
Differential expression analysis for protein and gene expression upon 24h EDC exposure. **(a)** The number of differentially expressed genes (DEGs) and proteins (DEPs) upon a 24h-exposure to sixteen different EDCs. For transcriptomic analysis, thyroid OoC devices were exposed to two different EDC concentrations of 1 nM and 10 µM. *P* < 0.01 was considered statistically significant. **(b)** Numbers of significantly enriched GO biological processes and Reactome pathways. Enrichment analysis was performed per EDC using lists of DEPs, and a combined list (both concentrations) of DEGs, as presented in (b). The number of GO terms/pathways which were enriched both at gene and protein level are presented as ‘in common’. A GO or pathway was considered statistically significant when Adj P < 0.05 and at least three DEG/DEP in the GO/pathway. **(c)** Significantly enriched biological processes and pathways at gene level (blue), protein level (orange) and both (red) for all sixteen EDCs. Several central terms representative of overarching biological processes were determined. Multiple GO terms and Reactome pathways are subsumed under a single central term. In our analysis, if any of the subordinate processes exhibited a statistically significant enrichment, it was positively labelled in the tabular representation. OPFR, organophosphate flame-retardants; PAH, polycyclic aromatic hydrocarbons; PCB, polychlorinated biphenyls.

The MFBs were capable of screening 54 samples per run (216 in total) with a success rate of up to 90%. A few devices failed primarily because of the appearance of air bubbles in the cell chambers. This issue was mitigated by using a positive pressure set-up^29^. RNA sequencing resulted in a median raw read count of 64.9 million per sample. LC-MS/MS analysis resulted in the identification of an average of 3503 proteins. For both transcriptomic and proteomic datasets, principal component analysis (PCA) of only vehicle controls from each run revealed that the samples were clustered based on the screening run, indicating that the main source of variation among samples across experiments is related to batch-to-batch variations (**Figure S6**). Differential expression analysis was performed for each EDC exposure compared to the vehicle control and numbers of differentially expressed genes (DEGs) and proteins (DEPs) (*P* < 0.01) are shown in **Figure 2a**. There was limited overlap between DEGs and DEPs (**Figure S7**). The number of DEPs and DEGs upon MMI exposure varied substantially across the screenings, suggesting again biological variation across batches (**Figure S8**). Heat maps of differentially expressed genes revealed clustering of samples within classes, for instance BkF, DahA and 10 µM BaP in the class of PAH, and

DEHP and DINP for the phthalates (**Figure S9**). Overall, chemicals of the OPFR class had the least impact on the number of DEGs, while exposure to BkF, BaP, DahA, and PCB126 resulted in the highest number of DEGs (**Figure 2a**). As expected, the highest dosage led for most EDCs to the greatest number of DEGs. Some exceptions were BADP, PCB118, PCB138, DEHP, and DINP, where the lowest concentration resulted in higher numbers. At the protein level, PCBs showed the strongest effects leading to up to 130 DEPs. We next performed functional enrichment analysis of gene ontology (GO) terms and Reactome pathways on the transcriptomics and proteomics data. BaP, DahA, BkF, PCB126, and DEHP caused gene-level activation of many biological pathways, while PCBs and BADP, followed by BkF and DahA triggered the most pathways at the protein level (**Figure 2b**). Among all EDCs, PCB126 revealed the highest correlation between both omics by exhibiting 216 biological processes dysregulated at gene and protein levels.

The main activated pathways are illustrated for each EDC in **Figure 2c**. BaP, BkF, and DahA induced gene-level activation of numerous processes related to cellular response to various stresses - including chemical stress, oxidative stress and endoplasmic reticulum (ER) stress - apoptosis, DNA damage, and metabolism of RNA and proteins. In addition, various signal transduction pathways were enriched, including signaling by TGF-β, SLIT/ROBO and nuclear receptors. In contrast, BaA exposure led to dysregulation of only rRNA processing pathways. Overall, PCBs caused significant pathway changes both at gene and protein level, with a substantial overlap, particularly for PCB126. Significant enrichment was observed for processes related to cellular respiration, DNA replication, and ECM organization, including laminin interactions, collagen formation and integrin-mediated signaling (**Table S1**). PCB118, 126, and 138 regulated signaling by tyrosine and by SLIT/ROBO receptors, recently identified as mediators in cancer progression^30,31^, as well as cellular responses to chemical stress. In particular, PCB153 showed gene-level responses to ER stress and apoptosis.

Interestingly, PCB126 resulted in activating thyroid carcinogenic processes. Among the phthalates, DEHP and DINP caused regulation of multiple pathways involved in chemical and oxidative stresses and the KEAP1-NFE2L2 pathway, apoptosis, ECM organization, and signal transduction pathways, while DIDP and DnOP exposure affected metabolism of proteins and cell junction organization. In the class of OPFRs, BADP induced a strong cellular response to stress both at gene and protein levels, as well as regulation of several signaling pathways. Regulation of processes associated with ECM organization was observed for all OPFRs (**Table S1**).

In addition to examining enriched pathways, we evaluated the activation of the aryl hydrocarbon receptor (AhR) pathway. Some PAHs and PCBs are known agonists of AhR, and consequently inducers of the cytochromes *Cyp1a1* and *Cyp1b1*, two members of the cytochrome P450 (CYP450) family. BkF was found to be the strongest inducer of these Cyp genes at 10 µM, followed by DahA, BaP, and BaA. In addition, 10 µM PCB126 induced *Cyp1a1* and *Cyp1b1* expression (**Figure S10**).

### 2.4. EDC class effects

To unravel omics signatures of the different EDC classes, we performed pathway enrichment analysis of genes and proteins differentially expressed by at least three out of the four EDCs per class. While no overlapping DEPs were observed in the PAH class, we found a large overlap in DEGs within this class, with a focus on cellular responses to stress, such as ER, chemical, and oxidative stress involving the KEAP1-NFE2l2, ROBO receptors, and AhR pathways (**Figure 3**). Similar stress responses have been reported following acute PAHs exposures in animal models^32^ and in occupational workers^33,34^. Oxidative stress markers *Nqo1* and *Cyp1a1* consistently ranked among the top15 upregulated DEGs for most of the PAHs, along with *Plin2*, suggesting a potential response to ER stress^35^ (**Table S2**). Other interesting genes in common included the *Ctsl* gene, which was strongly upregulated following exposure to BaP, DahA, and BaA, and *Nid2*, a gene encoding for a cell adhesion glycoprotein and among the top15 downregulated genes across all PAH exposures.

**Figure 3.**
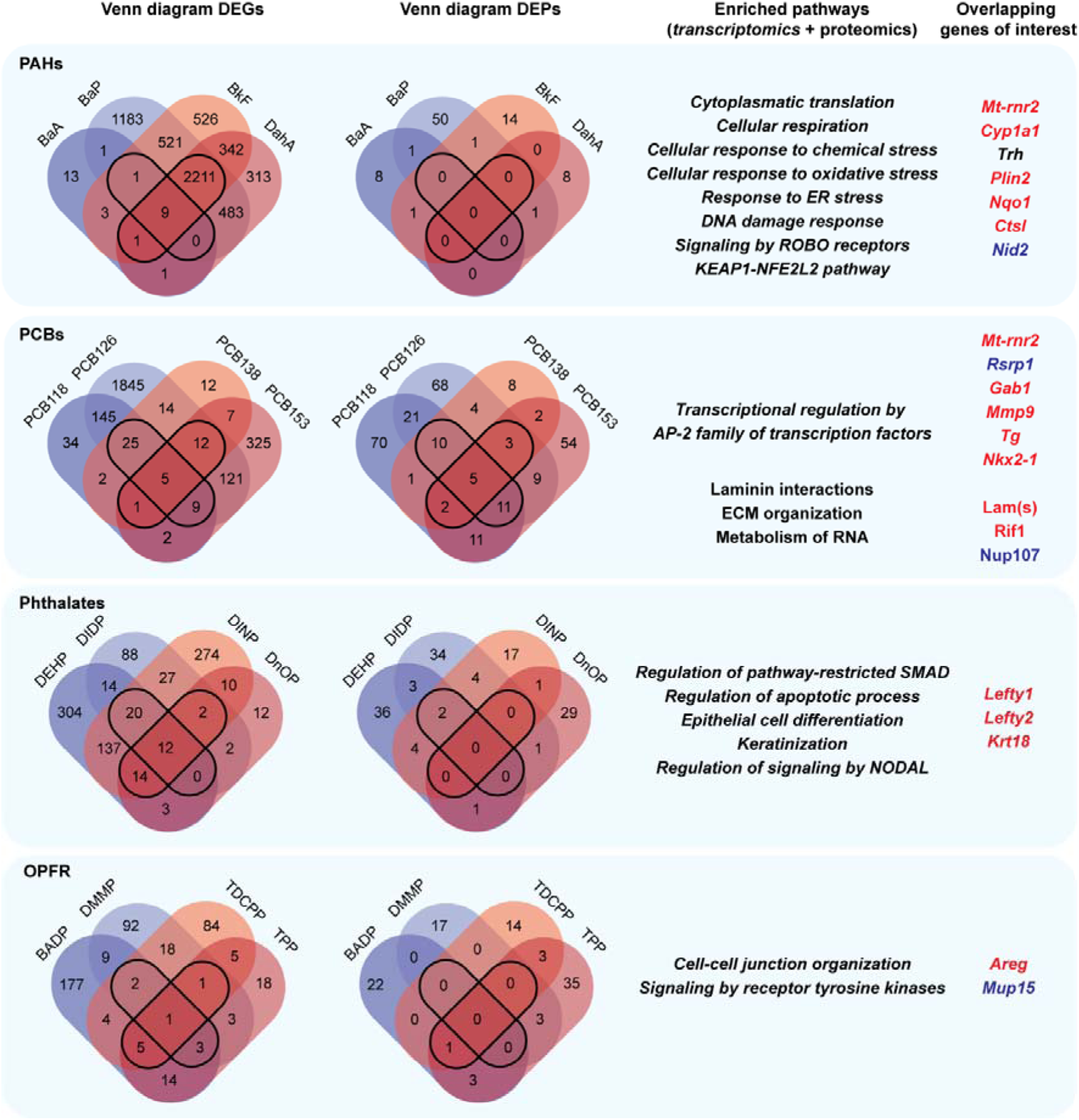
Identification of transcriptomic and proteomic signatures for each EDC class. Venn diagrams highlight the overlapping number of DEGs and DEPs across at least three out of four EDCs within the same class. Subsequently, functional enrichment analysis was performed for the list of overlapping DEGs and DEPs. Representative significantly enriched processes are stated for each class. The significantly enriched pathways were identified using Reactome and gene ontology (GO) databases. Pathways obtained from transciptomic databases are italicized. Genes of interest were identified based on their relevance in class-specific enriched pathways and their expression levels. Genes (italic) and proteins of interest belonging to the top15 upregulated or downregulated are marked in red and blue, respectively. Statistical significance threshold for pathway analysis was set at *P* < 0.05.

At the transcriptional level, the PCB class was characterized by transcriptional regulation by the AP-2 family associated with PCB-induced oxidative stress^36,37^. Notably, the mitochondrial gene *Rnr2* emerged consistently on the top5 upregulated DEGs across most of the PAHs and PCBs compounds (**Table S2, S3**), indicating its potential as a universal signature of EDC-induced cellular stress. Furthermore, all PCBs caused a strong upregulation of key thyroid genes for the synthesis, metabolism, and secretion of thyroid hormones, suggesting a class effect on the dysregulation of T_4_ synthesis, in contrast to other classes (**Figure 4, Figure S11**). Importantly, differential expression of these genes was also detected for the 1 nM concentration, indicating thyroid dysregulation even at very low concentrations of EDCs. For PCB-153, PCB-126, BkF, and DahA, the low concentration was potent enough to dysregulate multiple thyroid genes, including *Tpo*, *Tg*, *Dio1*, and *Trh* (**Figure 4, Table S4, and S5**). Functional enrichment analysis of overlapping DEPs revealed regulation of pathways related to ECM organization, laminin interactions, and metabolism of RNA (**Figure 3**). Several laminin subunit proteins (including LAMB1, LAMB2, LAMC1) are consistently upregulated in response to all PCBs. Interestingly, the PCB class caused differential expression of the Rif1 (up) and Nup107 (down) proteins, which play key roles in the DNA damage response and in the nuclear pore complex (NPC) assembly, respectively^38,39^.

**Figure 4.**
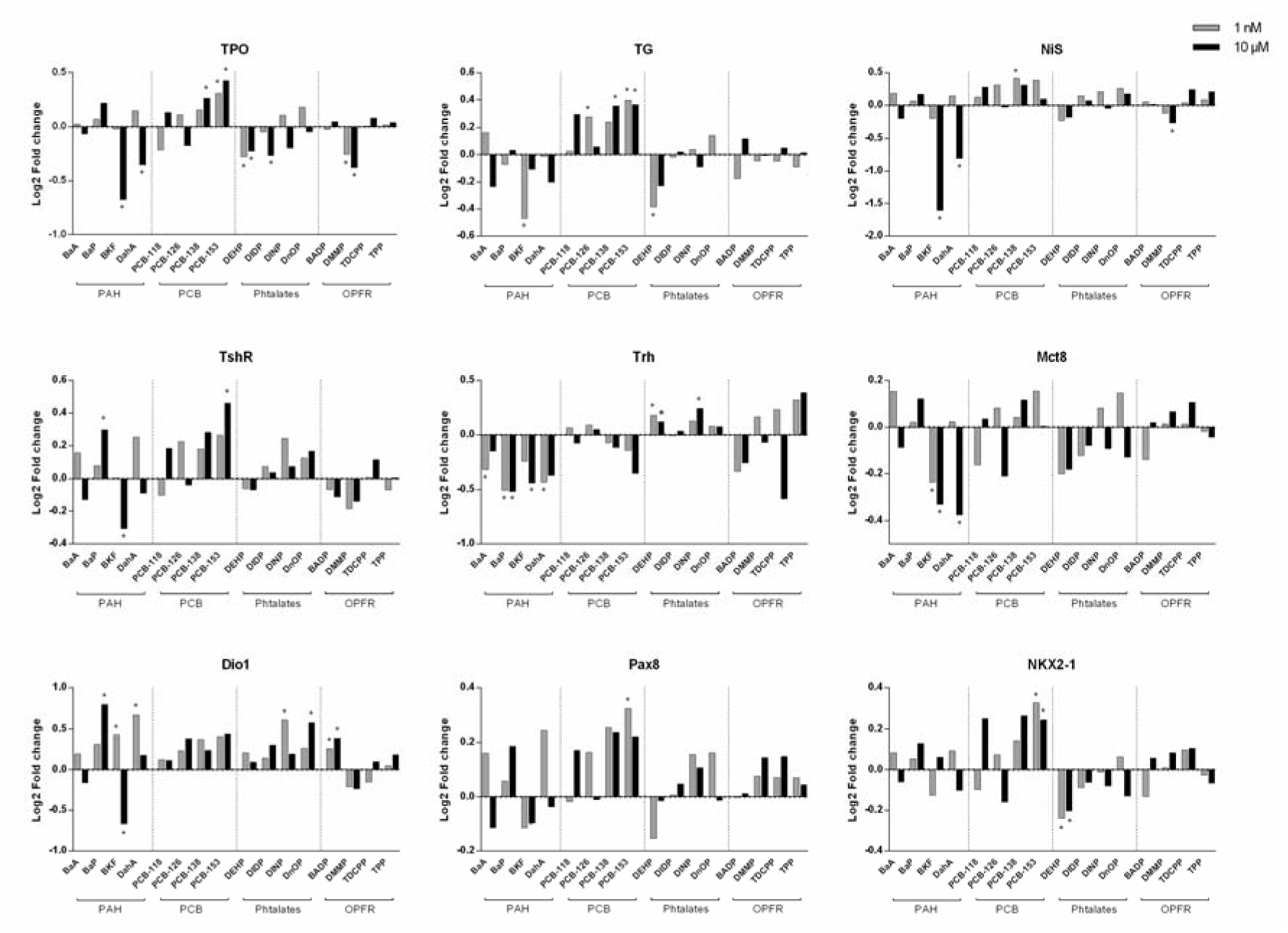
Regulation of key thyroid genes upon EDC exposure. Gene expression analysis (RNAseq) of thyroid OoC exposed to multiple EDCs for 24h, presented as Log2 fold change relative to vehicle controls (DMSO) \**P* < 0.01. See Figure S10 for additional genes. Tpo, thyroperoxidase; Tg, thyroglobulin; Nis, sodium iodide symporter; TshR, thyroid stimulating hormone receptor; Trh, thyroid releasing hormone; Mct8, monocarboxylate transporter 8; Dio1, iodothyronine deiodinase.

At the gene level, phthalates were characterized by enriched GO term pathways related to keratinization and regulation of signaling by NODAL, with specific genes (*Lefty1* and *Lefty2*) appearing on the top15 upregulated DEGs for all phthalates (**Table S6**). These genes are inhibitors of the TGF-beta/SMAD signaling pathway, which is known to affect thyroid function^40^. The phthalate class showed minimal overlap of DEPs, with only Fkbp2 and Tbca being differentially expressed following exposure to phthalates.

Finally, GO terms related to cell-cell junction and ECM organization processes were enriched for the overlapping DEGs in the OPFR class, along with signal transduction pathways, including the PI3K-Akt pathway. Notably, *Areg*, which has been reported to be involved in several stress responses^41,42^, appeared in the top15 upregulated genes for most OPFRs (**Table S7**). No significant protein signature was found for the OPFR class with Stx12 being the only shared DEP among three OPFRs. The top15 DEPs are listed in **Table S8**.

### 2.4. Prolonged exposure to EDCs

The human population is continuously exposed to low-dose environmental EDCs that can dysregulate thyroid functioning^43^. To emulate chronic exposure *in vitro* and to investigate duration effects of EDC exposure, we exposed thyroid OoC devices to BaP, DEHP, and PCB153 (two concentrations) for a duration of 10 days. MFBs were assembled and cells inside the OoC devices were perfused at a flow rate of 50 µL/min and cultured in serum-free differentiation medium to allow quantification of secreted T_4_ by mass spectrometry. The serum-free medium conditions were tested first and it was shown that thyroid morphology and functionality were not affected (**Figure S12**).

After prolonged EDC exposure, thyroid organoids were imaged and collected for posterior transcriptomic and proteomic analysis. By day 10, among the typically small, round thyroid organoids, we noticed an overgrowth of large, off-target (GFP-negative) cellular structures, which may have an impact on gene and protein expression (**Figure 5a**). Omics analysis of the total cell population revealed that, among the three prolonged exposures, 10 µM BaP and 1 nM DEHP resulted in the highest number of DEGs and DEPs, respectively (**Figure 5b**). Noticeably, the lowest concentration had a minimal effect on the transcriptome when compared to the 10 µM dosage. No overlap was found between DEGs and DEPs following prolonged exposure to BaP, PCB153 and DEHP (**Figure S13**).

**Figure 5.**
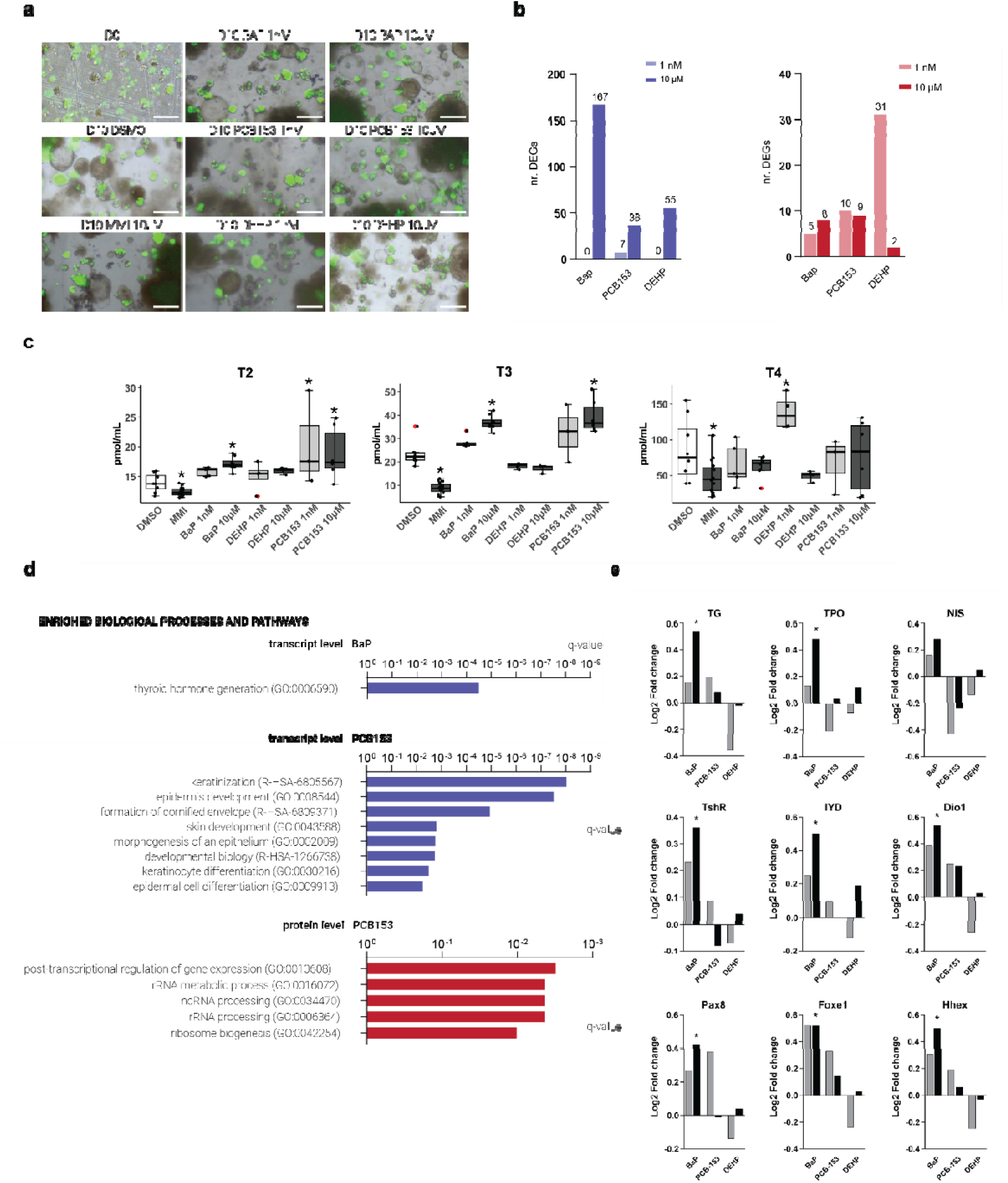
Effects of prolonged EDC exposure on thyroid OoC devices. **(a)** Brightfield images of thyroid organoids before (day 0) and after EDC exposure (day 10) at two concentrations (1 nM and 10 µM). Pictures taken with a 4x magnification. Scale bar, 250 µm. **(b)** Number of DEGs and DEPs upon 10 days-exposure to BaP, PCB153, and DEHP (1 nM and 10 µM). P < 0.01 was considered statistically significant. **(c)** Thyroid hormone concentrations after prolonged EDC exposure on thyroid OoC devices. Targeted quantification of the thyroid hormones 3,3’-T2, T3 and T4 in medium was performed upon 10 days-exposure to BaP, PCB153, and DEHP (1 nM and 10 µM, light and dark grey, respectively). Exposure to the reference compound Methimazol (MMI, 10 µM) and the vehicle control (DMSO) were included for each batch but showed no significant difference (students T-test, P<0.05) so they were pooled for further analysis. The box plots show the median, the lower and upper hinges, which correspond to the first and third quartiles and circles show individual replicates. The whiskers extend from the hinge to the largest/smallest value no further than 1.5x the inter-quartile range. * indicate significant difference from the vehicle control (Kruskal-Wallis non-parametric test followed by Dunn’s post hoc test, P<0.05). **(d)** Significantly enriched GOBP terms and Reactome pathways at transcript (blue) and protein (red) levels. Enrichment analysis was performed per EDC using combined lists (both concentrations) of DEPs, and DEGs. A GO term or pathway was considered statistically significant when q < 0.05 and at least three genes or proteins within the GO term/pathway were differentially expressed. No significantly enriched processes were found after prolonged DEHP exposure (transcript and protein level). **(e)** Fold change in expression of thyroid genes upon exposure to BaP, PCB153, and DEHP for 10 days (grey, 1 nM concentration; black 10 µM). RNA seq results are presented as Log2 fold change relative 0.5% DMSO (vehicle controls). * P < 0.01.

Thyroid hormone (TH) analysis by LC-MS/MS showed quantifiable levels of T2, T3 and T4 (**Figure 5c**). As expected, exposure to the reference compound MMI resulted in significantly lower levels of all three THs compared to the vehicle control, suggesting inhibition of TH production by MMI. In contrast, exposure to the tested EDCs led to higher levels of the hormones compared to the vehicle control. In particular, the 10 µM dosage of BaP and PCB153 significantly increased both T2 and T3, and 1nM of PCB153 also significantly increased T2. DEHP exposure (1 nM) resulted in a significantly higher T4 level compared to the control (P<0.05).

Pathway enrichment analysis showed that prolonged BaP exposure triggered a more targeted response in thyroid organoids as compared to 24h exposures (**Figure 5d**). Prolonged BaP exposure caused gene-level activation of the GO term ‘thyroid hormone generation’ with significant upregulation of key thyroid genes, such as *Tshr*, *Tpo*, *Tg*, *Pax8*, *Iyd*, *Foxe1*, *Duoxa1*, and *Dio1* (**Figure 5e**). Among these, *Tg*, *Foxe1*, and *Iyd* were listed on the top10 upregulated genes (**Table S9**). Prolonged exposure to PCB153 resulted in gene level regulation of epidermal cell pathways with dysregulation of genes coding cytokeratins (**Figure 5d**) and of the Stratifin (Sfn) protein. Congruently, exposure to PCBs has previously been shown to disrupt keratins expression in a breast cancer cell line^44^ and in mice^45^. On protein level, PCB153 exposure resulted in significant enrichment of processes related to RNA metabolism, with differential expression of the nucleolar proteins NOL6, NOLC1, NOL8 (**Table S10**). The latter two were also DEPs upon BaP exposure. Interestingly, both concentrations of PCB153 and BaP exposure resulted in downregulation of parafibromin protein (CDC73), which functions as a tumor suppressor protein in parathyroid carcinomas^46^. Finally, sigma-2 receptor protein (TMEM97), another cancer-related protein, was differentially expressed by BaP (both concentrations), 10 uM PCB153 and MMI, and 1 nM DEHP. This protein plays a role in controlling cholesterol levels^47^, which has been shown to be increased following exposure to BaP^48^ and PCB153^49^ in mice.

## 3. Discussion

Traditional thyroid assays based on 2D cell lines fail to recreate the luminal compartment where T_4_ is synthetized and they lack the hemodynamic microenvironment, which is detrimental for preserving the functionality of a glandular tissue such as the thyroid^12,17^.

While most of these assays are compatible with common high-throughput technologies^23^, they lack important morphological and functional features to comprehensively understand the mechanisms involved in thyroid exposure to EDCs. Moreover, currently available assays normally evaluate thyroid disruption upon exposure to EDCs at concentrations ranging tens to maximum hundreds of µM, which are not representative concentrations of the daily human exposure^50,51^. Here, we combined a microphysiological thyroid model with omics technologies to develop an advanced platform for the identification and assessment of EDC-driven thyroid disruption. Following features of this customized platform allowed us to overcome some of the aforementioned limitations.

mESC-derived thyroid organoids were cultured in geometrically-defined and perfused compartments in our microfluidic platform. Previously, we demonstrated that mESC-derived thyroid organoids preserve a 3D follicular-like structure *in vitro* and restore plasma T_4_/T_3_ levels in athyreotic mice after transplantation^11^. Furthermore, when exposed to flow, perfused thyroid organoids enhanced T_4_ synthesis, suggesting a positive effect of perfusion on thyroid functionality^12^. Considering these findings, we believe that the present thyroid OoC model more closely replicates the native thyroid microenvironment and functionality when compared to conventional 2D thyroid cultures.

To facilitate the omics analysis and culture of thyroid organoids in scalable microfluidic devices, we employed a novel, reversible LnP clamping concept. Recently, LnP clamps have been shown to facilitate the culture of 3D printed constructs and stem cell-derived organoids on-chip while improving the efficiency of cell harvesting for the performance of multiple analytic downstream assays^27^. As reported previously, a clamp sealing can yield up to a 2-fold increase in extracted RNA content when compared to permanently bonded chips^52^ and therefore constitutes a good strategy for implementation of omics analysis when high cell numbers are needed^27,53^. Our torque-adjustable LnP clamp enabled reversible (un-)loading of thyroid organoids inside the OoC device in a single step and within only a few minutes. The short assembly times were essential to successfully scale up the experiment to 54x cell chambers per screening. Moreover, the LnP clamp enabled a continuous and manual refining of the sealing strength to compensate for any gradual loss of sealing strength throughout the culture period. This approach also added an extra safety level, ensuring that potential leakages of harmful components were contained within microfluidic device, even in the event of sealing failure. For these reasons, adjustable LnP clamps hold promise for the development of next-generation microfluidic devices designed to support complex dynamic biological systems.

EDC-mediated thyroid disruption was evaluated based on changes in gene and protein expression. Differential expression analysis and subsequent pathway enrichment analysis successfully elucidated novel thyroid responses to multiple EDCs. Overall, the 24h-exposures showed that PAHs and PCBs triggered strong responses at gene and protein levels, respectively, whereas phthalates and OPFR chemicals caused smaller effects. PAHs and PCBs are known endocrine disruptors and agonists of the AhR. Here, we observed induction of *Cyp* genes, especially for BkF, DahA, BaP, BaA, and PCB126. These findings are congruent with previous studies, in which BkF was found to be the strongest AhR activator of the PAHs, followed by DahA^54–56^, while PCB126 was the most potent chemical among the studied PCB compounds^57^. Our results are in line with a publicly available adverse outcome pathway (AOP), in which activation of the AhR pathway is a molecular initiating event upon exposure to certain PCBs and dioxins^58^.

While each EDC exerted a variety of biological processes and pathways, distinct signatures were identified for each EDC class. In general, PCBs induced dysregulation in processes linked to ECM organization, whereas PAHs triggered cellular stress responses with activation of the KEAP1-NFE2l2 pathway, as previously reported in *in vivo* models^59,60^. Notably, phthalates consistently upregulated *Lefty1* and *Lefty2* genes, an effect that has remained undocumented until now. Further studies are needed to comprehensively decipher and validate these mechanisms in EDC-mediated toxicity.

The proper functionality of the thyroid gland is closely linked to the production of T_4_. Previous studies have reported EDC-induced alterations in gene expression of key molecules involved in iodide metabolism, including *Tpo*, *Nis*, and *Tg*. For instance, PCB126 was found to decrease expression of Nis^61,62^ while PCB118 decreased Tg expression^63^ in human thyroid cell lines. Other studies have reported dysregulation of *Tpo* and increased expression of Nis upon exposure to PAHs^64^ and DEHP^65,66^, respectively. Similarly, we demonstrated that 24h-exposure to some EDCs, especially PCBs caused dysregulation of the expression of key thyroid genes. For instance, BaP, BkF, PCB126, and PCB153 affected expression of multiple genes responsible for regulation of T_4_ production, such as *Tg*, *Nis*, *Tpo*, and *Mct8*. Importantly, some of these effects were also observed for the 1 nM dosing. Most data on adverse health effects of EDCs have been derived from studies using high concentrations resulting from industrial accidents and occupational exposures. In contrast, typical human exposure occurs at low concentrations from daily contact with small residues found in plastics, cosmetics, and food cans among others. Therefore, studying the effects of EDCs at very low concentration is relevant and desired. Whether EDCs can exert effects at low concentrations has yet to reach a consensus among the scientific community^2,50^. Here, we provided important data suggesting that EDCs, such as BaP and PCBs have thyroid-disrupting effects at very low concentrations of 1 nM.

When exposed to 10 µM EDCs for the extended period of 10 days, omics analysis revealed a more targeted response from thyroid organoids as compared to short-term exposures, while minimal effects were detected at the 1 nM concentrations. Notably, prolonged 10 µM BaP exposure induced significant upregulation of multiple thyroid genes crucial in T_4_ hormone production. These increased genes expression levels were accompanied by elevated levels of T2 and T3 in the culture medium after 10 days exposure. Previous studies have reported BaP-induced gene dysregulation, which was followed by changes of plasma thyroid hormone concentration in rodent thyroid glands^67^. While PCB153 and DEHP did not affect gene expression levels of these thyroid genes, exposure to these compounds showed a similar increase on TH levels. MMI showed the opposite effect, namely a decrease in TH levels, which is a known effect of MMI^68^.

As mentioned, the response to prolonged EDC exposure on the transcriptome and proteome level seemed attenuated when compared to the shorter exposure time. It remains unclear whether this stems from a cellular adaptation to EDCs over the course of time or from experimental factors. Although the 10 day-exposure to the low concentration of 1 nM provided a more realistic representation of daily human exposure to environmental pollutants than acute dosing, it does not fully emulate the chronic exposure to EDCs experienced in real-life. Throughout lifetime, humans are often continuously exposed to a complex mix of numerous chemicals, which can have a synergistic impact on thyroid functionality^69,70^. In future studies, integrating longer periods of culture and mixtures of chemicals into screening assays may offer additional insights. Another confounding factor could be that the mESC-derived thyroid organoid culture is a heterogeneous cell culture model, which exhibits a growing proportion of non-thyroid/off-target cells over time, complicating thyroid assessment by bulk sequencing and proteome analysis. In principle, conducting omics analysis on sorted thyroid cells could unravel such thyroid-specific effects for prolonged exposures. However, the practical implementation of this approach is challenging due to the limited experimental throughput.

Another challenge inherent to the model is the batch-to-batch variability. PCA analysis of the complete transcriptome and proteome showed clustering of samples from the same differentiation batch, indicating that this was a substantial source of variation. Stem cell-derived organoids are known to develop often with lack of consistency in size, composition, and functionality. In fact, this variability is a major issue limiting organoid technology^16^.

Previous studies have employed various bioengineering approaches for an improved spatiotemporal control over the microenvironment and to minimize organoid variability^71,72^. Similarly, we executed various steps to limit the interorganoid variability of our model. Prior to EDC exposure, thyroid organoids were enriched, counted, and seeded at high cell density into membranous carriers with well-defined dimensions. The defined and controlled arrangement of the microfluidic elements and tubing in our MFB allowed us to set up all flow circuits in a reproducible manner. Additionally, the flow rate was assessed with in-line liquid flow sensors in at least three circuits to discriminate any changes in flow rate over time.

Despite these efforts, some degree of organoid batch-to-batch variation still influenced technical reproducibility. This inherent variability, however, also reflects the biological richness and maturity of ESC-derived organoids, reinforcing their value as highly physiologically relevant models. As organoid technology continues to advance, scaling up platforms like ours will be critical for boosting throughput and sample sizes, enabling statistically robust conclusions. In addition, the integration of automated organoid handling and on-chip biosensing capabilities will further enhance consistency, data fidelity and reproducibility.

Collectively, the developed platform made a substantial stride towards the development of advanced thyroid cell-based assays, while bringing attention to major challenges in the field. The generated omics data hold the potential to help understanding the complex chain of events underlying EDC exposure, which is a crucial requirement for novel advanced *in vitro* assays. Additionally, our platform enables the investigation of thyroid disruption at a functional level. Importantly, this work can contribute to the formation of relevant AOPs and computational models of EDC modes of action, which ultimately will improve our understanding of thyroid disruption.

## 4. Conclusions and outlook

The developed platform allows quick and easy loading of thyroid organoids, which is key for the scalability of experiments with complex 3D *in vitro* models and for efficient sample/cell harvesting enabling omics analysis. By combining physiologically relevant thyroid organoids, microfluidics, and omics technologies, we successfully screened a panel of EDCs and elucidated thyroid responses to EDC exposure. Importantly, the developed platform demonstrated its sensitivity in detecting biological changes upon EDC exposure at a low concentration (1 nM). In the future, we plan to incorporate human-derived thyroid organoids in the described platform and screen the effects of EDC cocktails for longer exposure periods. This will be important to unravel mechanisms underlying chronic effects of EDCs on human thyroid surrogates. Furthermore, integration with automated instrumentation would be key to improve the current platform concerning reproducibility and interorganoid variability. Nevertheless, we believe that our innovative modular platform provides a new avenue for the development of next-generation *in vitro* tools to assess the effect of potential EDCs on tissues and organs.

## 5. Materials and Methods

### 5.1. Organoid culture

Thyroid follicles were differentiated from a recombinant mESC line (A2LoxNkx2-1-Pax8) as previously described^11,12,73^. In brief, the murine ESCs were routinely cultured on mouse embryonic fibroblasts (MEF) feeders and in DMEM supplemented with 15% ES Cell qualified FBS (Sigma Aldrich, St. Louis, USA), Leukemia Inhibitory Factor (LIF, IK0701, 1000 UmL^−1^) (ORF Genetics, Kopavogur, Iceland); non-essential amino acids (0.1×10^−3^ M, final), sodium pyruvate (1×10^−3^ M), penicillin and streptomycin (50 UmL^−^1, final), and 2-mercaptoethanol (0.1×10^−3^ M). mESCs were forced to aggregate into embryoid bodies (EBs) (1000 cells/droplet) by hanging drop culture in differentiation medium containing DMEM supplemented with 15% FBS, vitamin C (50 μgmL^−1^), non-essential amino acids (0.1×10^−3^ M), sodium pyruvate (1×10^−3^ M), penicillin and streptomycin (50 UmL^−1^), and 2-mercaptoethanol (0.1×10^−3^ M). After 4 days, EBs were collected, embedded in Matrigel Growth Factor Reduced (354230, Corning, New York, USA), and 50 μL drops were plated into 12-well plates. EBs were committed to the thyroid lineage by using differentiation medium supplemented with Doxycycline (1 μgmL^−1^) for 3 days, followed by 14 days in differentiation medium supplemented with 8-Br-cAMP (10×10^−6^ M, B 007, Biolog, Hayward, USA). After differentiation, organoids were extracted from Matrigel drops through digestion with an enzymatic cocktail of collagenase type IV (17104019, Gibco, Waltham, USA) (100 UmL^−1^) and dispase II (04942078001, Roche, Basel, Switzerland) (4 UmL^−1^) in HBSS (14175095, Gibco) at 37°C for a maximum of 1h30min. Subsequently, the cell suspension was enriched for thyroid follicles by filtering it through a 100 μm filter and reverse filtering using a 30 μm cell strainer (pluriSelect Life Science GmbH, Leipzig, Germany). Membranous carriers were cleaned with 70% EtOH, washed with ddH_2_O, placed in 12-well plates, and pre-coated with 5 μL of Matrigel for 5 min at 37°C to prevent organoid attachment to the bottom surface. Next, 3500 follicles in 30 μL Matrigel and 7000 follicles in 60 μL Matrigel were seeded in each carrier for transcriptomics and proteomics analysis, respectively. Prior to EDC exposure, follicles were pre-matured for 3 days in differentiation medium supplemented with 8-Br-cAMP (10×10^−6^ M) and TGF-,BRI inhibitor SB431542 (10×10^−6^ M, 1614, Tocris, Bristol, UK).

### 5.2. Engineering of the OoC device and flow batteries

CAD models of the OoC devices and the flow batteries were created using Solidworks (2015, Dassault Systems, Vélizy-Villacoublay, France). The housing components of the OoC device were made from PC and were fabricated by a company (Maastricht Instruments, B.V., The Netherlands). The device was completed by fastening the PC tapped body against an in-housed CNC milled PMMA window via a set of four M3 screws. Before usage, the acrylic window and the threads were lubricated with Vaseline. Fluidic chips were made by PDMS casting, while membranous carriers were fabricated by microthermoforming 50 µm-thick PC foils, as previously described^12^. In brief, PDMS (10:1) (Sylgard 184, Midland) was poured into pre-fabricated negative molds and degassed to remove air from the solution. The mould was closed with a flat lid and the system was compressed with a table C clamp. After curing at 80°C for 2h, PDMS chips were peeled off the moulds and cleaned. For the generation of membranous carriers, 50 μm thick PC films (it4ip, S.A., Louvain-la-Neuve) were compressed between a CNC milled brass mould and a counter-plate in a hydraulic hot press. The temperature was risen to 150°C and differential nitrogen pressure was applied to stretch the softened film into the negative mould. Afterwards, the plates and film were cooled down and the 3D formed film was demoulded. The flow batteries comprised four PMMA sheets (379575, VINK), which were laser cut and stacked by four corner metallic columns. The height of the upper acrylic layers was adjustable by tuning cylinder clamps installed on each column.

### 5.3. EDC exposure and microfluidic screening

Thyroid OoCs were exposed to 16 EDCs, listed in **Figure 1a**, for 24H, and subsequently to a selection of EDCs for 10 days. The 24H exposures were performed in differentiation medium supplemented with 8-Br-cAMP (10×10^−6^ M) and SB431542 (10×10^−6^ M), while for long-term exposure, we changed to serum-free differentiation medium from the start of exposure. Serum-free differentiation medium was supplemented with B-27 supplement minus vitamin A (1:50, Gibco), sodium iodide (NaI) (0.1 ×10^−6^ M), 8-Br-cAMP and SB431542. At day 5 of exposure, extra concentrations of 8-Br-cAMP and SB431542 were added to the medium. Thyroid follicles were exposed to 1 nM and 10 µM of the EDCs. EDC stock solutions were prepared in DMSO, requiring the addition of an equal amount of DMSO (0,5%) to culture media, including the control DMSO condition (vehicle control). Each OoC device contains biological triplicates of the same experimental condition (n=3); two OoC devices were allocated for the vehicle control condition (n=6). Because of the increased risk of sample loss during prolonged exposure, extra replicates were included (n=5 for transcriptomics analysis, n=4 for proteomics analysis).

One day prior to the start of exposure, 54 flow circuits were pre-assembled and inserted into three IPC-N Ismatec peristaltic pumps (78000-47, Metrohm, The Netherlands). All fluidic connections were established via barbed to male Luer slip adapters (CIL-P-854, IDEX) and Ø0.8 mm PharMed^®^ BPT tubing (224-0555, Saint-Gobain). On day 0, glass medium reservoirs, containing 15 ml of medium containing EDCs, and OoC devices were plugged in the flow batteries. In brief, after three days of static pre-maturation in the membranous carriers, thyroid organoids were stacked vertically together with two PDMS fluidic chips and tightened inside each OoC device (**Figure 1c**). Torques of 1.8 Nm and 2.3 Nm were applied to the OoC devices for the 24h and 10 days exposure experiments, respectively. Up to 18 OoC devices were plugged to the flow circuits pre-filled with cell culture medium. A flow rate of 12 µL/min was set for the 24h exposure runs, whereas a flow rate of 50 µL/min was used for the 10 days exposure experiments. Finally, the flow batteries were incubated inside an ICO240 incubator (Memmert GmbH + Co. KG, Schwabach, Germany) with active humidity control (set to 70% rh). After EDC exposure, the flow rate of three randomly-selected flow circuits was evaluated by a liquid flow rate sensor (SLF3S-0600F, Sensirion AG). Sensing data was acquired for two minutes and processed using GraphPadPrism 9 software. The OoC devices were opened and thyroid organoids were retrieved for RNA-seq and LC-MS/MS analysis as described in the respective sections Conditioned medium was collected and stored at - 80°C until use.

### 5.4. RNA sequencing

#### 5.4.1 RNA-Seq sample preparation and sequencing

Thyroid follicles, exposed to EDCs for 24h and 10 days, were collected in cold PBS, centrifuged,and lysed with QIAzol Lysis Reagent. The lysate was then used for total RNA extraction. Total RNA was extracted with the miRNeasy Micro Kit (Qiagen GmbH, Hilden, Germany). For the 10-day exposure samples, Poly(A) libraries were prepared from 1 µg of total RNA on an automated system (Zephyr G3® NGS) with the NEXTFLEX^®^ Rapid Directional RNA-Seq Kit 2.0 (NOVA-5198-02, PerkinElmer), NEXTFLEX^®^ Poly(A) Beads 2.0 (NOVA-512992, PerkinElmer) and NEXTFLEX^®^ Unique Dual Index Barcodes (NOVA-512923, PerkinElmer). 10 PCR cycles were performed. Conversely, for the 24h exposure samples, libraries were prepared from 20 ng of total RNA using the NEXTFLEX® Combo-Seq™ mRNA/miRNA Kit (NOVA-5139-53, PerkinElmer, Waltham, MA, USA), employing the NEXTFLEX® tRNA/YRNA Blocker during preparation to deplete tRNA and Y RNA fragments, and utilizing 14 PCR cycles. Both sets of libraries were sequenced on an Illumina NovaSeq 6000 platform; the 10-day samples were sequenced in paired-end mode on an S1 flowcell (200 cycles, v1.5), while the 24h samples were sequenced in single-end mode on an S4 flowcell (35 cycles, v1.5).

#### 5.4.2. RNA-Seq data processing

Data from the 10-day from poly(A) libraries was trimmed with fastp (v0.23.2)^74^ and quantified with Salmon (v1.9.0)^75^ using the mouse genome GRCm39 (version M27 of the primary assembly, Ensembl 104)^76^. The 24h exposure Combo-Seq libraries were processed according to our previously published CODA pipeline^77^ on the exact same genome version.

#### 5.4.3. Differential expression and gene ontology analysis

Differential expression analysis was performed with R using the DESeq2 package^78^. To select relevant differentially expressed genes (DEGs), a series of stringent filtering steps was applied during the differential gene expression analysis as outlined by the Omics Data Analysis Framework for regulatory application (R-ODAF) developed by our group^79^. In brief, a gene is considered expressed in any EDC vs vehicle control comparison if its count per million (CPM) value is ≥1 in at least 75% of the replicates of either group. A gene was considered differentially expressed when the False Discovery Rate (FDR) was less than 0.01. In addition, DEGs undergo a second filtering step to identify any spurious spikes. Differential expression analysis was performed by comparing each concentration (1 nM or 10 µM) (n = 3) to the vehicle control. For every EDC, a combined list of DEGs from 1 nM and 10 µM concentrations was used to perform gene ontology (GO)^80^ and pathway enrichment analysis on Reactome^81^ using enrichR^82^. We set an adjusted P value threshold of 0.05, and additionally we retained only the enriched pathways that contained at least 3 DEGs. To elucidate EDC class effects, we generated a list of DEGs overlapping in at least 3 EDCs across each class and performed GO and Reactome pathway enrichment analysis. Overlapping genes and proteins were identified.

### 5.5. Proteomics analysis

#### 5.5.1. Protein sample preparation

Cells were lysed in urea lysis buffer and lysates were digested using trypsin on the S-Trap (Protifi) 96-well plate platform. Samples were fractionally distributed according to treatment condition across a 96 well plate to limit technical biases from the digestion process. Additional quality controls (NthyOri lysate and a reference serum sample from a healthy control) to measure digestion reproducibility were included. In brief, lysates were thawed from -80°C and proteins solubilized and denatured using a 10% SDS, 100 mM TEAB (pH 7.55) solution, added at a 1:1 ratio. Proteins were reduced and alkylated using final concentrations of 10 mM DTT and 40 mM Iodoacetamide, respectively. S-Trap binding buffer (9:1 ratio of methanol:TEAB) was added to each sample and loaded onto a 96-well S-Trap plate. After multiple wash steps, sequencing grade modified trypsin (Promega, Madison, USA) was used to digest the proteins overnight (16 h) at 37°C. After digestion, peptides were recovered and dried down using a vacuum centrifuge.

Subsequently, samples were prepared for discovery LC-MS/MS (Bruker timsTOF Pro coupled to an EvoSep One LC system). In brief, all mESC cell digests and quality controls were resuspended in 2% acetonitrile/0.1% formic acid and peptides were quantified using A215 on a DeNovix DS-11 spectrophotometer. 500ng of peptide were loaded onto EvoSep EvoTips using the manufacturers’ protocol. Samples were introduced onto a Bruker timsTOF in pre-determined worklist run orders. Worklists were flanked by commercially available HeLa cell line digests to monitor instrument performance over time. Samples were injected in order of their plate position to maintain fractional distribution across the digestion plate layout.

#### 5.5.2. LC-MS/MS data acquisition

LC-MS/MS data acquisition of the thyroid organoids (and quality control samples) was performed in a single batch. The 30 samples per day (SPD) EvoSep method (44-minute gradient) was used for these experiments. The timsTOF Pro mass spectrometer was operated in positive ion polarity with Trapped Ion Mobility Spectrometry (TIMS) and Parallel Accumulation Serial Fragmentation (PASEF) modes enabled. The accumulation and ramp times for the TIMS were both set to 100 ms with an ion mobility (1/k0) range from 0.62 to 1.46 Vs/cm. Spectra were recorded in the mass range from 100 to 1,700 m/z. The precursor (MS) Intensity Threshold was set to 2,500 and the precursor Target Intensity set to 20,000. Each PASEF cycle consisted of one MS ramp for precursor detection followed by 10 PASEF MS/MS ramps, with a total cycle time of 1.16 s. Data acquired from the LC-MS/MS runs were searched using Max Quant v1.6.17.0, in a single batch using the ‘match between runs’ to enhance feature detection. HeLa and Nthy Ori quality controls were run using the same parameters, with the human UniProt Fasta file substituted with the mouse equivalent. Further data processing was performed using Perseus v1.6.15.0 software, where protein data was filtered according to internal workflows. First, proteins identified as ‘potential contaminants’ by Max Quant were removed, followed by those that appeared in the reverse protein database. Remaining proteins were exported and analysed further.

#### 5.5.3. Differential expression analysis

Univariate statistical analysis was performed using a Students t-test on each EDC vs the vehicle control within the screening run. A p-value of 0.01 was used as a cut off to determine significance. Proteins were only considered valid if they appeared in at least three samples in one of the test groups (i.e., vehicle control or EDC exposed). For every EDC, the list of DEPs was used to perform GO and Reactome pathway enrichment analysis using enrichR. We set an adjusted P value threshold of 0.05, and additionally we retained only the enriched pathways that contained at least 3 DEPs.

### 5.6. Thyroid hormone quantification by LC-MS/MS

#### 5.6.1 Sample preparation

Samples were analyzed based on the method described elsewhere^83^. In brief, 1 mL medium was spiked with isotopic-labelled (^13^C)-thyroid hormone standards (internal standards (IS), cT2, cT3 and cT4), vortexed and enriched using solid-phase micro-extraction (SOLAµ HRP 10 mg/1 mL 96 well plate, ThermoFisher Scientific, Germany). After elution and evaporation to dryness the samples were reconstituted in 100 µL 5% methanol containing an instrument control standard (ICS, crT3). Procedural blanks containing water instead of medium were included from the beginning.

#### 5.6.2 Data acquisition

The targeted analysis of THs was performed on an Agilent 6495c triple-quadrupole system with a hyphenated Agilent 1290 Infinity II ultra-high performance liquid chromatography (UHPLC) system (binary pump, degasser, and autosampler; Agilent Technologies) as previously described^83^. Targeted analytes were thyroxine (T4), 3,3’,5-triiodothyronine (T3), 3,3’,5’-triiodothyronine (rT3), 3,5-diiodothyronine (3,5-T2), 3,3’-diiodothyronine (T2), 3-iodothyronine (T1), thyronine (T0), 3-iodothyronamine (T1Am), 3-iodothyroacetic acid (T1Ac), 3,5-Diiodothyroacetic acid (Diac), triiodothyroacetic acid (Triac) and tetraiodothyroacetic acid (Tetrac). LoDs as fmol on column for all THs were 4.0 (T4), 0.79 (T3), 0.76 (rT3), 0.56 (T2), 0.60 (3,3’-T2), 0.74 (T1), 0.23 (T0), 0.05 (T1Am), 9.4 (T1Ac), 15 (Diac), 10 (Triac), 9.6 (Tetrac). Neat standard ten-point equimolar calibration curves (0.04 – 20.0 pmol/mL TH, n=2) were prepared in 5 % methanol and all vials contained a fixed amount IS and ICS (100 pmol/mL).

#### 5.6.3 Quality assurance

A pooled QC sample was used to monitor instrument fluctuations during the run. Variability for individual samples were monitored by comparing the signal from the instrument control standard (crT3) and all signals were normalized to their corresponding C13-labelled internal standard to correct for recovery. To allow for the intended high-through-put application, the sample preparation was kept to a minimum. This resulted in high background signals for some samples, which decreased the reliability of the result. Hence results with a signal-to-noise ratio below 3 were excluded. The number of replicates remaining for analysis is given in Table S11 along with the measured concentration averages and standard deviations.

#### 5.6.4 Data analysis

Data analysis was conducted in MassHunter version 10.1 (Agilent Technologies). Quantifiable hormones were T4, T3 and T2. T1Am and T0 were also detected, but they were also present in the pure medium (data not shown), hence their concentrations were not relatable to the organoid biology and effects of EDCs so they were not analysed further. Statistical analysis and visualization were done using R ver. 4.4.1^84^. The univariate statistical analysis was performed by Kruskal-Wallis non-parametric test followed by pairwise comparison by Dunn’s test^85^. To allow for statistical comparisons, TH levels below the limit of detection (LoD) were replaced by ½ LoD. Data between the LoD and limit of quantification (LoQ) remained as they were. The data was visualized using ggplot2^86^.

## Acknowledgements

This study was financially supported by the European Union’s Horizon 2020 Research and Innovation Programme under grant agreement no. 825745.

