## Supplementary Information for "Microphysiological Flow Batteries For Dynamic EDC Screening Of mESC-derived Thyroid Organoids"

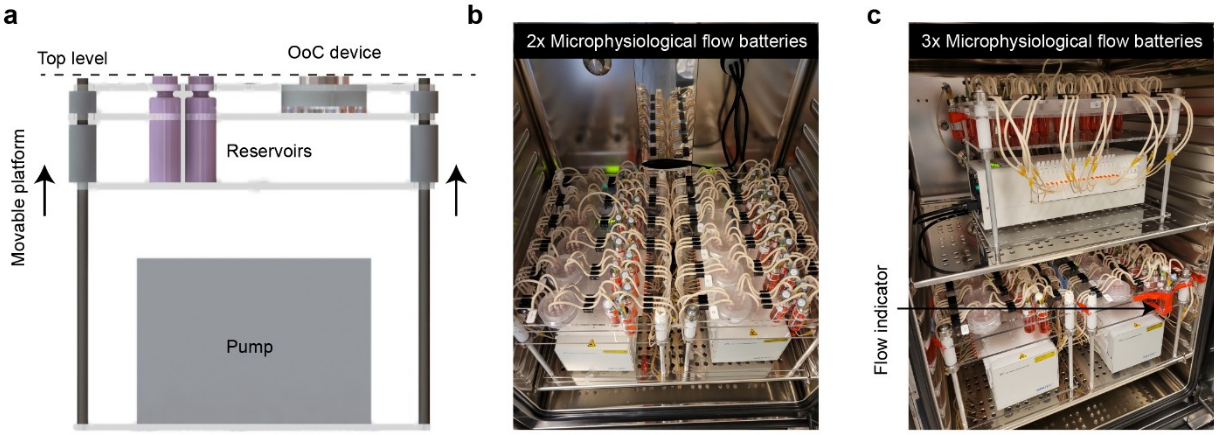


**Figure S1 – Microphysiological flow batteries. (a)** Side view of the CAD model of a microphysiological flow battery (MFB). The upper levels can be lifted up or even removed to allow easy plugging of the 2-stop pump tubings in the pump. Cell culture medium reservoirs and OoC devices are levelled at the top (dashed line) to shorten the amount of tubing required to connect them. Photographs of two **(b)** and three **(c)** fully assembled MFBs placed inside a $CO_2$ incubator with active humidity control. The relative humidity was set to 70% to prevent any damage to the electronic component of the pumps. An extra flow circuit was added to each battery acting as a flow rate indicator **(c)**.

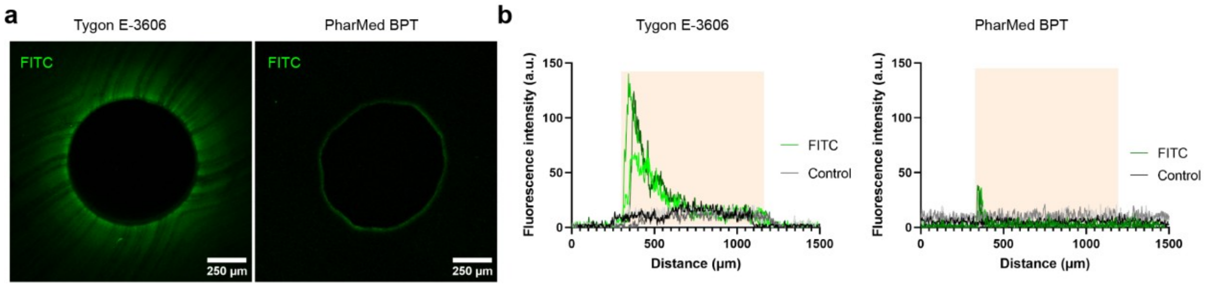


**Figure S2 – Characterization and selection of the tubing material.** The ab-/adsorption properties of the PharMed® BPT, a pharmaceutical grade tubing, were compared to a commonly used Tygon tubing. To evaluate the extent of compound loss in the tubing, we perfused them with a FITC solution for 24 hours and subsequently analysed the presence of any fluorescence signal by confocal microscopy. **(a)** Confocal images of cross-sections of Tygon E-3606 vs PharMed BPT tubings after being perfused with FITC for 24h. **(b)** Profiling of the fluorescence intensity along the tubing cross-section. The distance of 0 µm represents the centre of the luminal space while the yellow band represents the tubing body. As a control, tubings were perfused with $ddH_2O$.

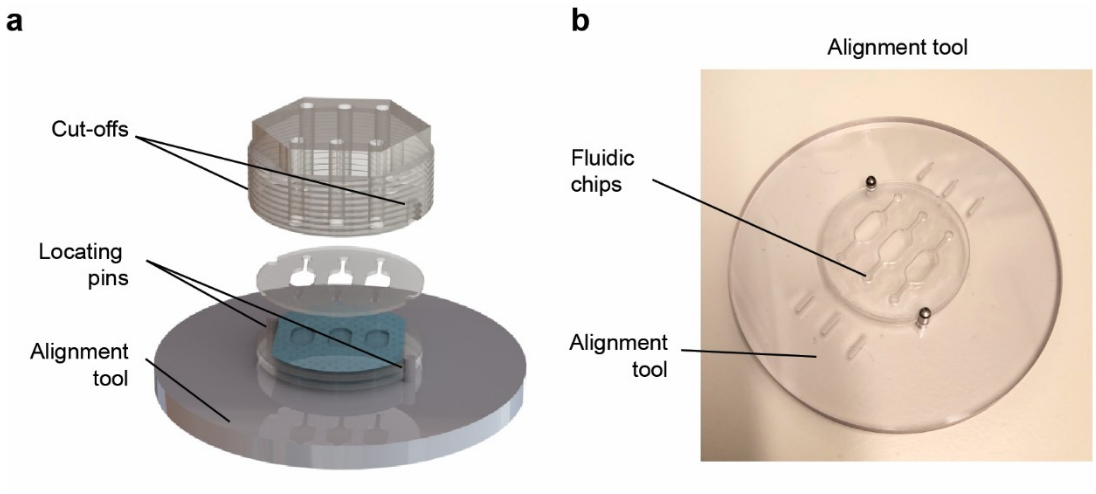


**Figure S3 – Design of the alignment tool. (a)** CAD model of the OoC being assembled while using the alignment tool. The coverslip, fluidic chips, and screw cap are aligned via two metallic locating pins of the alignment tool. Subsequently, the system is transferred and screwed with the tapped body. **(b)** Photograph of the alignment tool with two fluidic chips.

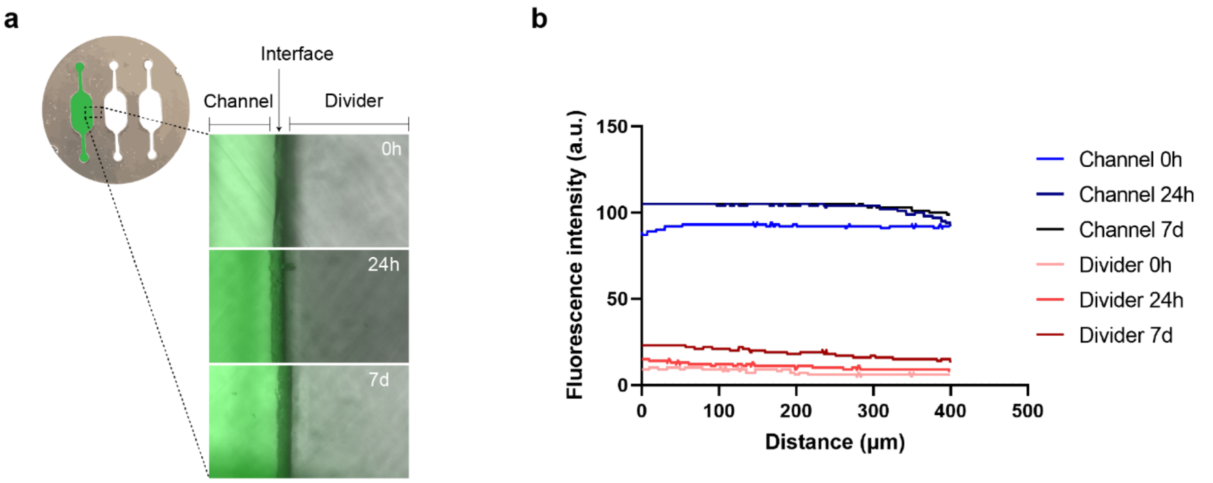


**Figure S4 – Leakage and cross-contamination test. (a)** Photographs of possible FITC cross-contamination between neighbouring channels in the developed OoC device. One fluidic channel was perfused with FITC at 50 µL/min while the neighbouring channels were kept with $ddH_2O$. Leakage through channels was monitored over a period of seven days. **(b)** Quantification of the fluorescence intensity (arbitrary units, a.u.) in channels and divider regions at different time points.

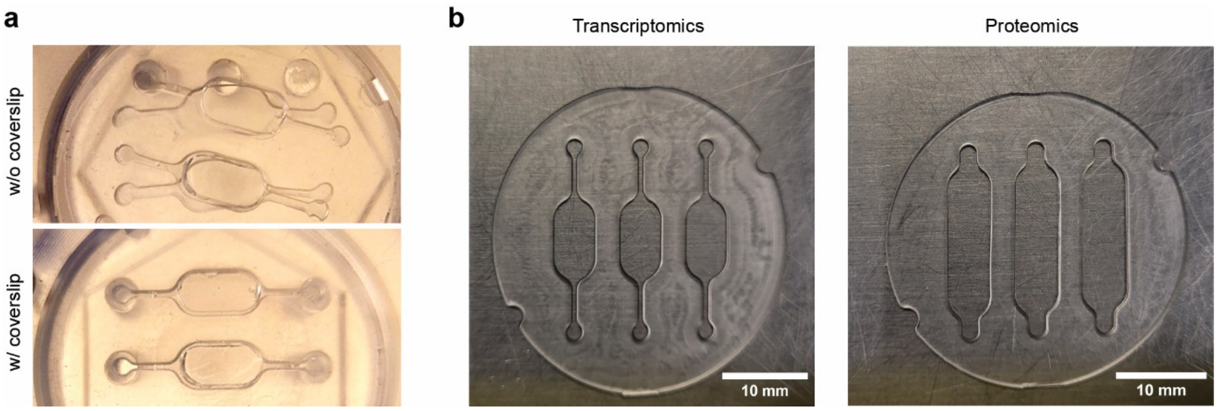


**Figure S5 – PDMS fluidic chips. (a)** Photographs of OoC devices with vs without a bottom coverslip after tightening. **(b)** PDMS fluidics chips for transcriptomics and proteomics. Scale bar, 10 mm. Gaskets were made by PDMS casting and contain two side cut-offs to allow alignment of multiple units during OoC assembly.

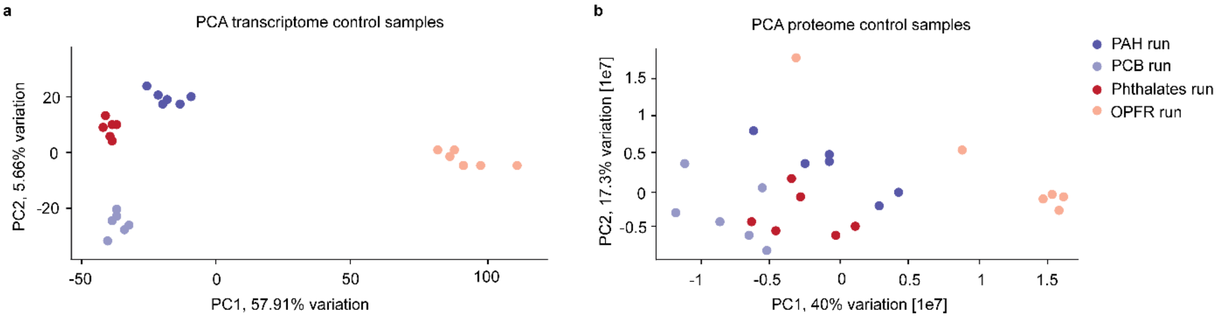


**Figure S6 – PCA analysis of DMSO controls across all four experimental runs. (a)** PCA analysis of the transcriptome of DMSO control samples. **(b)** PCA analysis of the proteome of DMSO control samples.

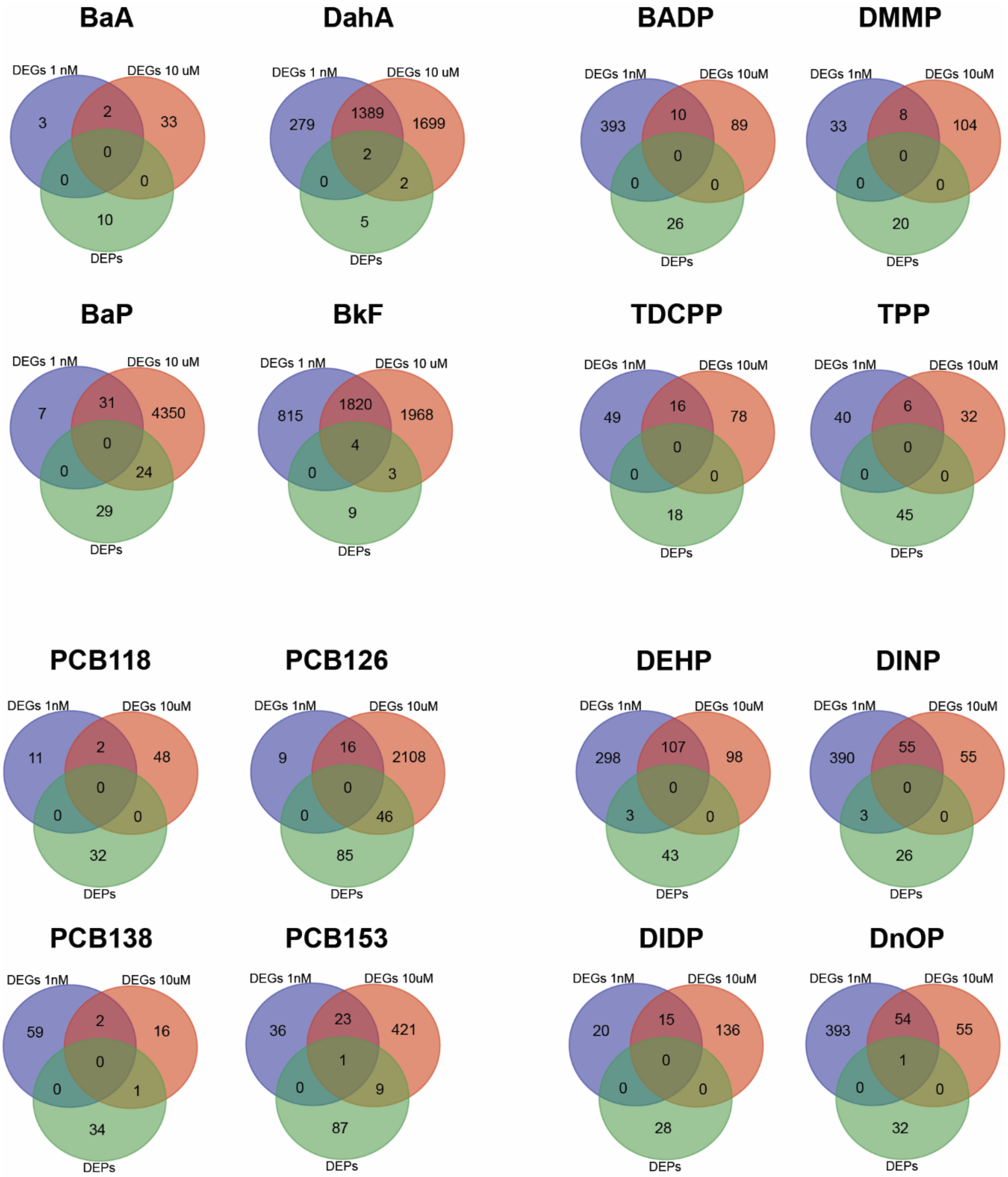


**Figure S7 – Venn diagrams depicting the overlap in DEGs 1 nM, DEGs 10 μM, and DEPs for all 16 EDCs in response to 24h exposure.**

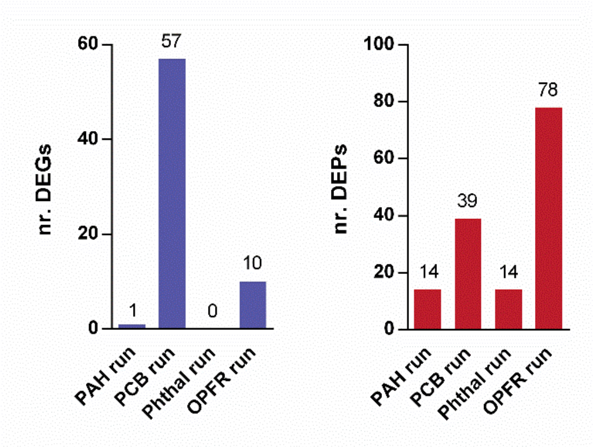


**Figure S8 – Number of DEGs and DEPs after 24H exposure to MMI.** MMI exposure (10 μM) was included in all four screening runs, and differential expression analysis of MMI vs control was performed per screening run.

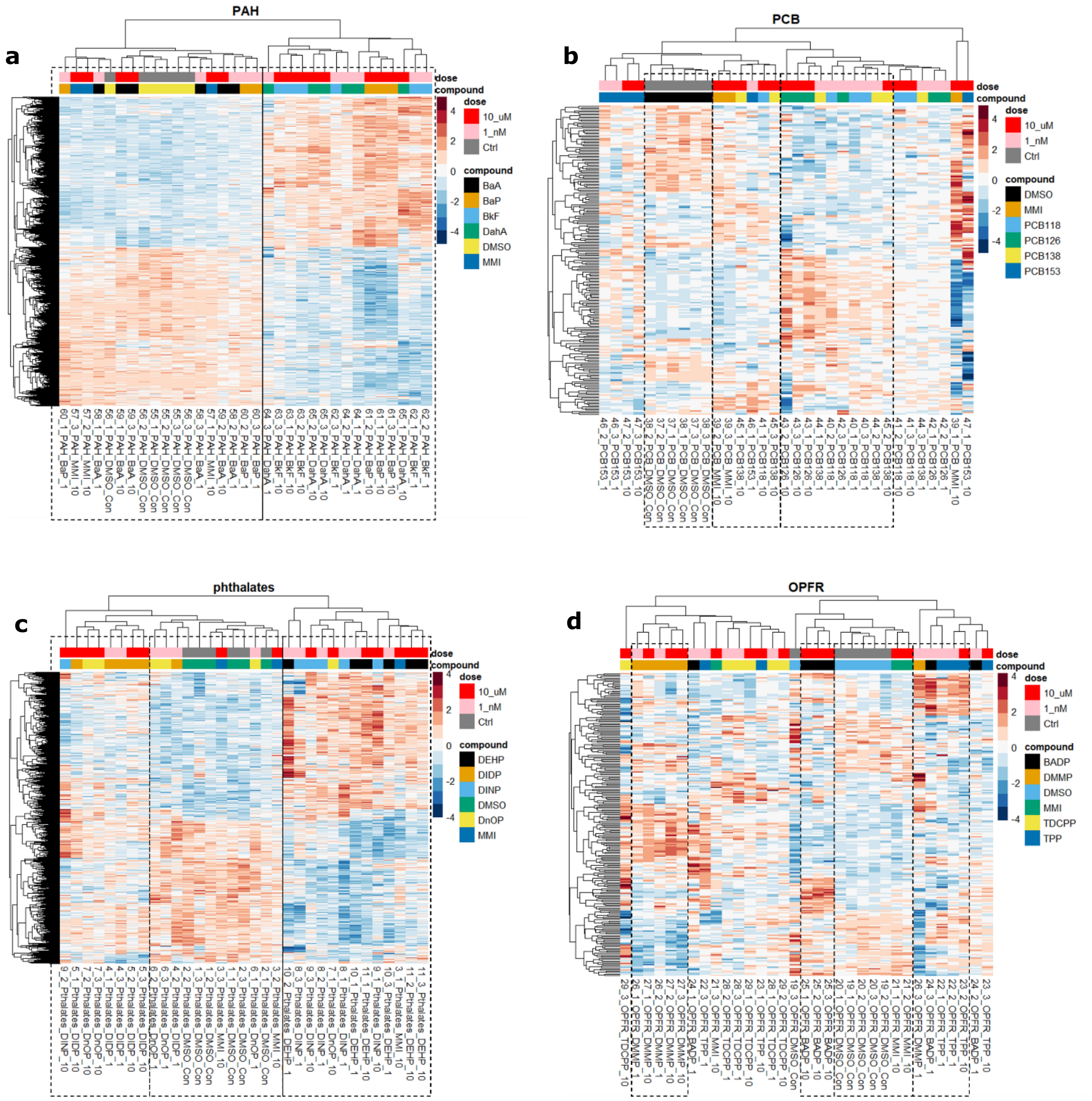


**Figure S9 – Heatmap of selection of differentially expressed genes within the different screening runs**. **(a)** PAH, **(b)** PCB, **(c)** Phthalates, **(d)** OPFR run. The selection was performed by identifying the genes which are differentially expressed in at least one EDC in the class vs DMSO control comparison within the analysed run. Some interesting subsets of samples based on the pattern of gene expression have been highlighted by the dashed boxes.

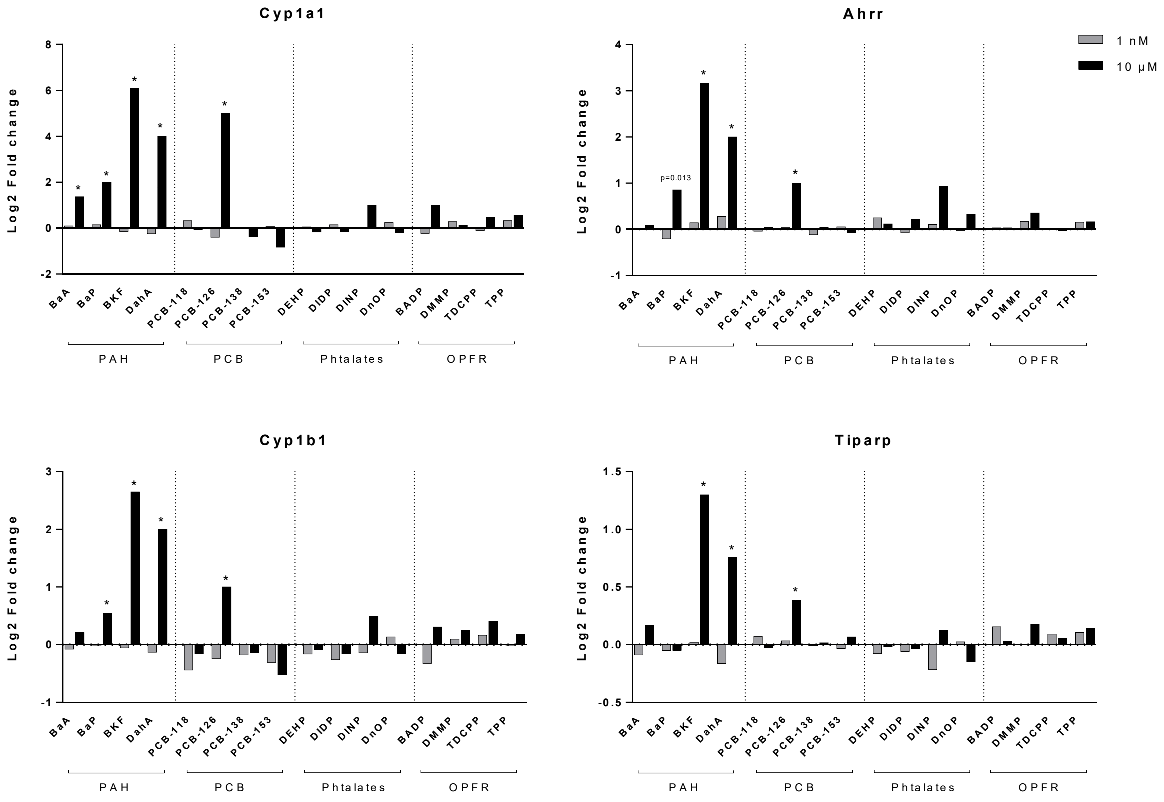


**Figure S10 – Gene expression of genes in involved in the aryl hydrocarbon receptor (AhR) pathway.** RNA-seq gene expression data of thyroid organoids exposed to different EDCs in the MFBs for 24H, shown as log2 fold change relative to unexposed control. * P < 0.01. Ahrr, aryl hydrocarbon receptor repressor. Cyp1a1, Cyp1b1 (cytochrome P450 enzymes) and TIPARP (TDCC inducible poly(ADP-ribose)polymerase) are AhR targets.

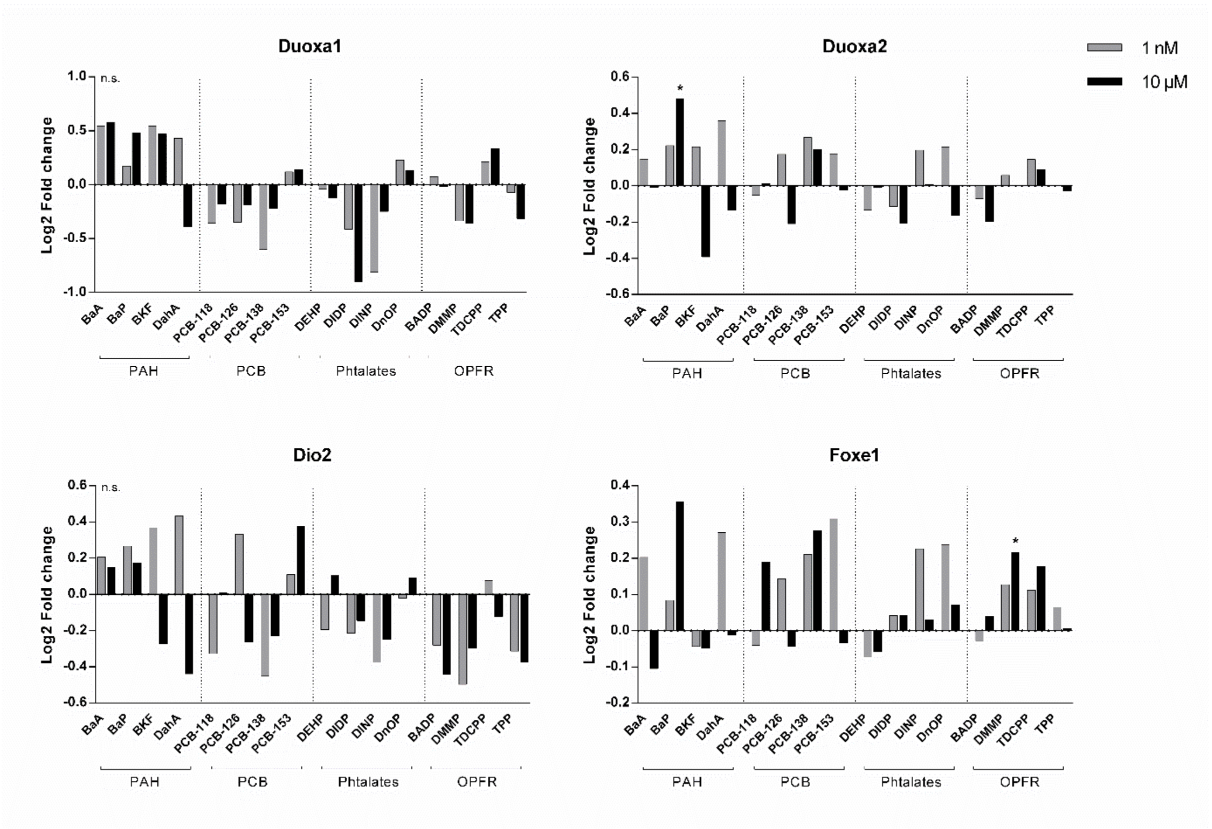


**Figure S11 – Regulation of thyroid genes upon EDC exposure.** RNA seq gene expression results of thyroid OoC exposed to multiple EDCs for 24h, presented as Log2 fold change relative to DMSO controls. **P* < 0.01. Duoxa1, dual oxidase maturation factor 1; Duoxa2, dual oxidase maturation factor 2; Dio2, iodothyronine deiodinase 2; Foxe1, forkhead Box E1.

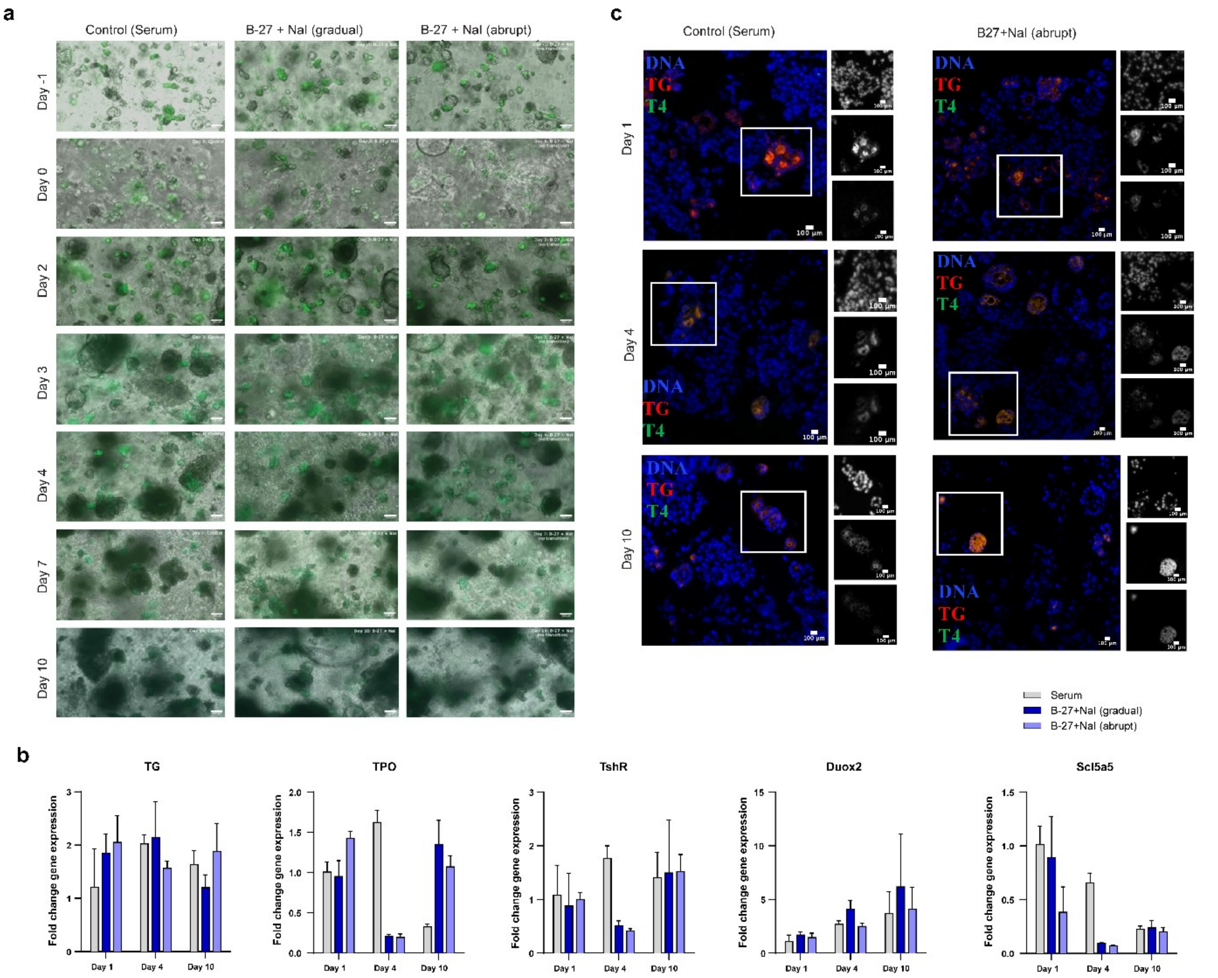


**Figure S12 – Thyroid functionality is maintained in serum-free culture medium.** Culture in serum-free medium (supplemented with B-27 and NaI) was monitored following a gradual transition from control medium (with serum) at day -3 to serum-free medium at day 0 (gradual) or following an abrupt change to serum-free medium at day 0. **(a)** Brightfield pictures overlapped with GFP signal from TG thyroid follicles monitored over the course of 11 days. Scale bar, 100 µm. **(b)** qPCR analysis of key thyroid genes. No dramatic changes in gene expression are observed when comparing the different media. Data are normalized by PAX8 expression and presented as fold change relative to control medium at day 1. Data from one experiment (n=3). Error bar indicates SD. **(c)** Immunofluorescence images of thyroid follicles cultured in control medium (serum) or serum-free medium (B-27 + NaI) for up to 10 days. Follicles were stained against TG (red) and T4 (green) and counterstained with DAPI. Individual channels are shown on the right side of each fluorescence image (from top: DAPI, TG, T4). Scale bar, 100 µm.

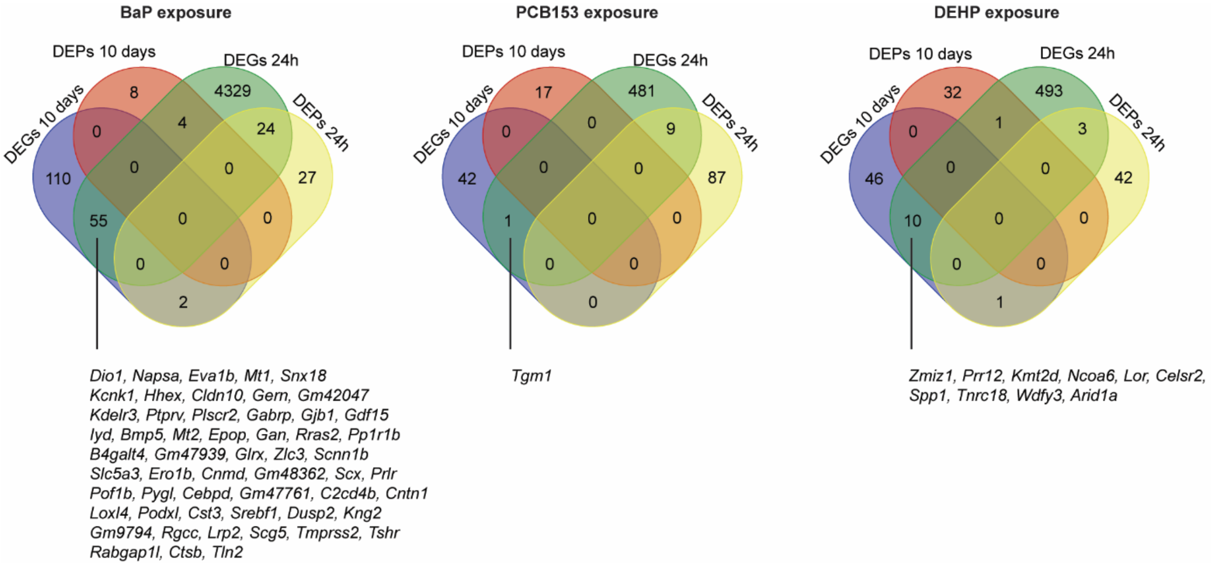


**Figure S13 – Venn diagrams showing the overlap of DEGs and DEPs following exposure to EDCs for 24 hours and for 10 days.** The list of DEGs was a combined list of both doses (1 nM and 10 µM). There was no overlap in DEPs when comparing short and long exposure. The genes that are differentially expressed both after 24 and 10 days exposure are mentioned in the figure.

**Table S1. DEGs and DEPs in processes related to ECM and cell junction organization.**

| | BaA | BaP | | | | BkF | | | | DahA | |
|---|---|---|---|---|---|---|---|---|---|---|---|
| D | | ACTN1 | CRB3 | ITGB3 | PXDN | AFDN | COL4A1 | ITGA6 | PRKCI | AGRN | LUM |
| E | | AFDN | CRK | ITGB4 | RAP1A | ARVCF | COL4A2 | ITGB1 | PXDN | CD44 | NID2 |
| G | | ARHGEF6 | CSK | LAMA1 | RASGRP2 | CADM3 | COL4A3 | ITGB4 | RHOC | CD47 | VCAM1 |
| s | | BCAR1 | CTNNB1 | LAMA5 | RSU1 | CD151 | COL4A4 | LAMA5 | SDK2 | COL18A1 | |
| | | CADM1 | CTSB | LAMB1 | SDK1 | CD9 | COL5A1 | LAMB3 | SMAD7 | COL4A3 | |
| | | CD151 | CTSL | LAMB3 | SDK2 | CDH1 | COL6A3 | LAMC1 | TJP1 | COL4A4 | |
| | | CDH2 | DMTN | LAMC1 | SOS1 | CDH11 | COL6A6 | LAMC2 | TLN1 | COL9A3 | |
| | | CDH24 | DST | LAMC2 | SRC | CDH17 | CRB3 | LOXL2 | TLN2 | FN1 | |
| | | CDH5 | EMP2 | LIMS1 | SYK | CDH2 | CTNNB1 | MMP3 | TRPV4 | FZR1 | |
| | | CLDN10 | F11R | LMNB2 | TLN1 | CDH24 | CTSB | MPDZ | VCL | HSPG2 | |
| | | CLDN4 | FBLIM1 | LOXL2 | VASP | CDH3 | CTSL | NECTIN1 | WDR1 | ITGA2 | |
| | | CLDN6 | FLNA | MMP3 | | CDH5 | DST | NECTIN4 | | ITGA5 | |
| | | COL15A1 | FLNC | NECTIN1 | | CDH6 | F11R | NID2 | | ITGA7 | |
| | | COL18A1 | FN1 | NID1 | | CLDN10 | FSCN1 | PALS1 | | ITGA8 | |
| | | COL2A1 | FZR1 | NID2 | | CLDN12 | FZR1 | PARD3 | | ITGA9 | |
| | | COL4A1 | GRB2 | PALS1 | | CLDN18 | GJB1 | PATJ | | ITGB1 | |
| | | COL4A2 | HSPG2 | PARD3 | | CLDN4 | GJB6 | PCOLCE | | ITGB1 | |
| | | COL4A3 | ITGA2 | PARD6B | | CLDN6 | GJC1 | PIP5K1C | | ITGB7 | |
| | | COL4A4 | ITGA6 | PCOLCE | | CLDN7 | HSPG2 | PKP1 | | LAMA5 | |
| | | COL5A1 | ITGA7 | PDPK1 | | COL15A1 | ITGA1 | PKP3 | | LAMC1 | |
| | | COL6A3 | ITGB1 | PLEC | | COL18A1 | ITGA2 | PKP4 | | LAMC2 | |
| | | COL6A6 | ITGB1 | PRKCI | | COL2A1 | ITGA3 | PLEC | | LMNB2 | |

| | PCB118 | PCB126 | | | | | PCB138 | PCB153 | |
|---|---|---|---|---|---|---|---|---|---|
| D | | ADAM10 | COL2A1 | ITGAV | NCAM1 | SDC3 | COL2A1 | AGRN | LAMA5 |
| E | | ADAMTS9 | COL4A2 | ITGB4 | NID2 | SH3PXD2A | COL6A3 | BGN | LTBP1 |
| G | | AGRN | COL7A1 | ITGB6 | P3H2 | TGFB2 | LAMA5 | BSG | MMP2 |
| s | | BSG | DAG1 | JAM2 | P4HA1 | | LOXL2 | CD44 | NID2 |
| | | CAPN6 | DDR1 | LAMA5 | P4HA2 | | MMP9 | COL18A1 | SDC4 |
| | | CAPNS1 | DST | LOXL2 | PLEC | | P4HA1 | COL1A2 | |
| | | CASK | FBLN1 | LRP4 | PLOD1 | | PLOD2 | COL25A1 | |
| | | CD47 | FBN1 | LTBP3 | PLOD2 | | VCAN | COL3A1 | |
| | | COL11A1 | FBN2 | MFAP2 | PTPRS | | | COL6A1 | |
| | | COL12A1 | FN1 | MMP10 | PXDN | | | COL6A3 | |
| | | COL14A1 | FURIN | MMP15 | SCUBE1 | | | CTSB | |
| | | COL17A1 | HSPG2 | MMP3 | SDC1 | | | HSPG2 | |
| D | LAMA1 | COL18A1 | LAMA1 | NID1 | | | COL18A1 | COL18A1 | LAMB1 |
| E | LAMB1 | CAPN2 | LAMA5 | NID2 | | | LAMA1 | COL4A1 | LAMB2 |
| P | LAMB2 | COL4A1 | LAMB1 | PPIB | | | LAMB1 | FLOT1 | LAMC1 |
| s | LAMC1 | COL4A2 | LAMB2 | | | | LAMC1 | HSPG2 | NID1 |
| | NID1 | HSPG2 | LAMC1 | | | | NID1 | LAMA1 | NID2 |
| | | | | | | | | LAMA5 | |

| | DEHP | | | | DIDP | DINP | | | DnOP | |
|---|---|---|---|---|---|---|---|---|---|---|
| D | AFDN | DAG1 | LAMA5 | NOTCH1 | | CD151 | CTSB | NID2 | CLDN4 | ITGA3 |
| E | AGRN | FBLIM1 | LAMB1 | OPTC | | COL17A1 | CTSL | P4HA2 | COL17A1 | JUP |
| G | CD151 | FN1 | LAMB2 | PLEC | | COL18A1 | ITGA3 | PPIB | COL17A1 | LTBP1 |
| s | COL17A1 | HSPG2 | LAMC1 | SDC4 | | COL2A1 | ITGB1 | | DAG1 | P4HA2 |
| | COL4A1 | ITGB1 | NECTIN1 | | | | | | | |
| | COL4A2 | LAMA1 | NID2 | | | | | | | |
| D | | | | | CLDN3 | | | | | |
| E | | | | | FLNA | | | | | |
| P | | | | | ITGA6 | | | | | |

| | BADP | | | | DMMP | | TDCPP | TPP | |
|---|---|---|---|---|---|---|---|---|---|
| D | AGRN | CTSD | LAMA1 | NID2 | CAPN6 | LAMC2 | CADM3 | BGN | COL1A2 |
| E | CD44 | DAG1 | LAMA5 | NOTCH1 | CLDN4 | PLOD1 | CDH11 | CDH11 | COL5A2 |
| G | COL17A1 | FN1 | LAMB1 | PLEC | CLDN6 | PXDN | CLDN4 | CLDN4 | DCN |
| s | COL1A1 | GREM1 | LAMB2 | SDC4 | COL4A3 | SCUBE3 | ITGA3 | CLDN6 | ITGA3 |
| | COL1A2 | HSPG2 | LAMC1 | TGFBI | COL4A4 | SPARC | ITGB4 | COL11A2 | LOX |
| | COL4A1 | ICAM1 | LOX | THBS1 | CTSL | THBS1 | JUP | COL1A1 | |
| | COL4A2 | ITGA3 | LOXL4 | | ITGA3 | | LAMC2 | COL1A1 | |
| | COL5A2 | ITGB1 | MMP9 | | LAMB3 | | | | |

**Table S2 – Top 15 upregulated and downregulated genes after 24h exposure to 1 nM and 10 µM of EDCs in the class of PAH (BaA, BaP, BkF, DahA).**

| BaA 1nM \| top15 upregulated | | |
|---|---|---|
| gene | log2 fold change | adj pvalue |
| mt-Rnr2 | 0.224 | 3.27E-04 |

| BaA 1nM \| top15 downregulated | | |
|---|---|---|
| gene | log2 fold change | adj pvalue |
| Gm15109 | -20.766 | 5.43E-05 |
| Gm42556 | -16.353 | 5.43E-05 |
| Gm50139 | -1.803 | 7.66E-03 |
| Trh | -0.320 | 5.10E-03 |

| BaA 10uM \| top15 upregulated | | |
|---|---|---|
| gene | log2 fold change | adj pvalue |
| Lce1a1 | 4.374 | 3.52E-05 |
| Lce1a2 | 3.994 | 7.85E-04 |
| Lce1f | 3.990 | 1.25E-04 |
| Dsc1 | 3.488 | 1.83E-06 |
| Kprp | 2.941 | 6.60E-06 |
| Klk7 | 2.315 | 5.30E-07 |
| 2610528A11Rik | 2.268 | 1.53E-05 |
| Dsg1a | 2.206 | 6.34E-05 |
| Them5 | 1.869 | 7.85E-04 |
| Cyp1a1 | 1.370 | 6.38E-07 |
| Agr2 | 1.229 | 2.64E-04 |
| Calml3 | 0.980 | 6.17E-05 |
| Aqp4 | 0.853 | 5.42E-05 |
| mt-Rnr2 | 0.370 | 4.94E-16 |
| Socs3 | 0.362 | 1.68E-04 |

| BaA 10uM \| top15 downregulated | | |
|---|---|---|
| gene | log2 fold change | adj pvalue |
| Gm49359 | -23.797 | 2.07E-07 |
| Gm15109 | -19.800 | 5.42E-05 |
| Gm49388 | -19.429 | 8.38E-05 |
| Aldh1a3 | -0.399 | 5.53E-03 |
| Prickle1 | -0.370 | 5.03E-03 |
| Rsrp1 | -0.267 | 3.66E-07 |
| Sesn3 | -0.206 | 3.20E-03 |
| Fat1 | -0.160 | 1.33E-03 |

| BaP 1nM \| top15 upregulated | | |
|---|---|---|
| gene | log2 fold change | adj pvalue |
| Pagr1a | 2.892 | 9.16E-17 |
| mt-Rnr2 | 0.457 | 1.69E-23 |
| Arhgef6 | 0.403 | 1.85E-03 |
| Gas6 | 0.385 | 8.30E-05 |
| Gm28438 | 0.312 | 1.97E-06 |
| Plin2 | 0.292 | 6.81E-03 |
| Cd24a | 0.253 | 5.23E-08 |
| Ctsl | 0.245 | 7.67E-05 |
| Anxa1 | 0.238 | 2.07E-03 |
| Ctsb | 0.207 | 8.38E-06 |
| Tsc22d1 | 0.165 | 3.80E-03 |
| mt-Nd1 | 0.157 | 5.03E-05 |
| mt-Nd2 | 0.140 | 5.87E-03 |
| mt-Cytb | 0.132 | 8.30E-05 |
| Mt2 | 0.126 | 6.81E-03 |

| BaP 1nM \| top15 downregulated | | |
|---|---|---|
| gene | log2 fold change | adj pvalue |
| Gm42713 | -29.173 | 5.23E-13 |
| Gm15109 | -25.682 | 2.33E-09 |
| Usp26 | -0.534 | 1.33E-04 |
| Prtg | -0.398 | 1.45E-04 |
| Sohlh2 | -0.383 | 4.46E-03 |
| Gm42047 | -0.352 | 2.00E-03 |
| Rsrp1 | -0.251 | 6.02E-09 |
| Sptbn2 | -0.243 | 4.46E-03 |
| Slc2a3 | -0.238 | 2.31E-06 |
| Zdbf2 | -0.237 | 2.00E-03 |
| Lima1 | -0.222 | 7.67E-05 |
| Lars2 | -0.208 | 5.21E-05 |
| Mycn | -0.206 | 1.55E-04 |
| Gldc | -0.178 | 2.49E-03 |
| Nid2 | -0.177 | 8.38E-06 |

| BaP 10uM \| top15 upregulated | | |
|---|---|---|
| gene | log2 fold change | adj pvalue |
| Adh7 | 1.525 | 1.76E-48 |
| Nqo1 | 1.330 | 1.37E-57 |
| Slc7a11 | 0.950 | 2.62E-45 |
| Plin2 | 0.866 | 8.77E-36 |
| Ccng1 | 0.777 | 5.85E-123 |
| Esd | 0.759 | 7.16E-40 |
| Trp53inp1 | 0.634 | 3.02E-36 |
| Sdcbp | 0.607 | 2.29E-39 |
| Ctsl | 0.591 | 5.02E-41 |
| mt-Rnr2 | 0.547 | 8.46E-36 |
| Ftl1-ps1 | 0.499 | 2.88E-50 |
| mt-Cytb | 0.442 | 6.22E-79 |
| mt-Co1 | 0.434 | 3.54E-54 |
| mt-Nd2 | 0.428 | 1.61E-35 |
| mt-Nd1 | 0.416 | 1.37E-46 |

| BaP 10uM \| top15 downregulated | | |
|---|---|---|
| gene | log2 fold change | adj pvalue |
| Nid2 | -0.783 | 2.99E-114 |
| Trim71 | -0.763 | 2.01E-84 |
| Arid1a | -0.734 | 2.29E-136 |
| Sall4 | -0.705 | 1.30E-85 |
| Larp1 | -0.680 | 6.39E-77 |
| Srcap | -0.669 | 4.08E-90 |
| Ankrd11 | -0.666 | 2.86E-104 |
| Prrc2a | -0.664 | 7.21E-110 |
| Brd4 | -0.633 | 1.72E-77 |
| Igf2bp1 | -0.632 | 3.32E-78 |
| Hcfc1 | -0.622 | 5.51E-89 |
| Prrc2c | -0.563 | 2.34E-78 |
| Srrm2 | -0.545 | 1.82E-91 |
| Trim28 | -0.529 | 2.34E-78 |
| Huwe1 | -0.519 | 4.88E-84 |

| BkF 1nM \| top15 upregulated | | |
|---|---|---|
| gene | log2 fold change | adj pvalue |
| Plin2 | 0.605 | 2.39E-25 |
| Gm28438 | 0.509 | 1.45E-36 |
| mt-Rnr2 | 0.544 | 4.71E-32 |
| Anxa2 | 0.486 | 8.37E-22 |
| mt-Rnr1 | 0.425 | 1.01E-23 |
| mt-Co1 | 0.362 | 5.56E-25 |
| mt-Cytb | 0.361 | 1.73E-30 |
| mt-Nd1 | 0.348 | 2.31E-24 |
| Rpl10-ps3 | 0.317 | 8.37E-22 |
| Rpl18a | 0.312 | 8.68E-22 |
| Rpl7 | 0.295 | 1.88E-23 |
| Rpl3-ps1 | 0.288 | 2.42E-22 |
| Tpt1 | 0.286 | 2.73E-27 |
| Rpl13a | 0.276 | 3.18E-21 |
| Eef1a1 | 0.255 | 8.68E-22 |

| BkF 1nM \| top15 downregulated | | |
|---|---|---|
| gene | log2 fold change | adj pvalue |
| Notch3 | -0.702 | 1.05E-26 |
| Hspg2 | -0.592 | 1.65E-30 |
| Kmt2d | -0.550 | 4.48E-42 |
| Lama5 | -0.549 | 1.29E-27 |
| Fat1 | -0.525 | 6.80E-41 |
| Nrk | -0.503 | 1.22E-27 |
| Ago2 | -0.496 | 5.42E-28 |
| Dync1h1 | -0.486 | 5.68E-28 |
| Erdr1 | -0.474 | 2.06E-28 |
| Polr2a | -0.469 | 1.50E-26 |
| Kmt2a | -0.467 | 2.39E-25 |
| Srcap | -0.450 | 3.02E-28 |
| Brd4 | -0.429 | 7.93E-25 |
| Prrc2b | -0.422 | 3.77E-28 |
| Ubr4 | -0.405 | 1.19E-26 |

| BkF 10uM \| top15 upregulated | | |
|---|---|---|
| gene | log2 fold change | adj pvalue |
| Cyp1a1 | 6.084 | 1.51E-164 |
| Ahrr | 3.162 | 2.22E-77 |
| Tgfbi | 2.704 | 0 |
| Cyp1b1 | 2.647 | 3.33E-113 |
| Nqo1 | 2.105 | 9.61E-66 |
| Adh7 | 2.049 | 1.27E-111 |
| Atp6v0a4 | 1.527 | 9.94E-82 |
| Tiparp | 1.298 | 1.77E-119 |
| Selenbp1 | 1.283 | 2.56E-90 |
| Slc7a11 | 1.039 | 2.44E-62 |
| Acaa2 | 0.965 | 4.46E-70 |
| Esd | 0.869 | 2.28E-63 |
| Emp1 | 0.839 | 1.90E-57 |
| Chd1l | 0.770 | 3.94E-73 |
| Nfe2l2 | 0.693 | 2.15E-69 |

| BkF 10uM \| top15 downregulated | | |
|---|---|---|
| gene | log2 fold change | adj pvalue |
| Slc5a5 | -1.608 | 7.23E-41 |
| Adm2 | -1.570 | 1.32E-43 |
| Tpo | -0.674 | 1.48E-58 |
| Timp3 | -0.627 | 7.85E-29 |
| Adgrl1 | -0.507 | 7.43E-28 |
| Nid2 | -0.479 | 3.49E-50 |
| Zfp462 | -0.429 | 1.87E-29 |
| Kmt2d | -0.423 | 1.41E-27 |
| Arid1a | -0.415 | 6.22E-29 |
| Spen | -0.408 | 1.19E-27 |
| Ago2 | -0.387 | 4.69E-28 |
| Srcap | -0.378 | 4.95E-35 |
| Polr2a | -0.377 | 1.42E-27 |
| Bptf | -0.337 | 9.52E-28 |
| Prrc2c | -0.321 | 5.28E-34 |

| DahA 1nM \| top15 upregulated | | |
|---|---|---|
| gene | log2 fold change | adj pvalue |
| Fos | 0.935 | 1.55E-11 |
| Gpx2 | 0.742 | 4.62E-11 |
| mt-Rnr2 | 0.619 | 1.86E-22 |
| Ctsl | 0.534 | 2.54E-15 |
| Lgals1 | 0.505 | 3.44E-12 |
| mt-Cytb | 0.450 | 5.92E-24 |
| Ftl2-ps | 0.401 | 5.73E-11 |
| Mt2 | 0.395 | 3.60E-11 |
| Ftl1-ps1 | 0.393 | 3.37E-14 |
| Gpx1 | 0.376 | 2.29E-11 |
| mt-Nd1 | 0.362 | 3.69E-19 |
| mt-Rnr1 | 0.355 | 3.06E-10 |
| Ccng1 | 0.353 | 4.21E-13 |
| mt-Nd2 | 0.337 | 1.10E-11 |
| mt-Co1 | 0.302 | 3.71E-12 |

| DahA 1nM \| top15 downregulated | | |
|---|---|---|
| gene | log2 fold change | adj pvalue |
| Aire | -1.241 | 5.24E-18 |
| Slc16a3 | -0.920 | 2.20E-45 |
| Pfkp | -0.653 | 1.06E-18 |
| Pfkl | -0.626 | 1.91E-27 |
| Notch3 | -0.621 | 7.49E-18 |
| Slc2a1 | -0.616 | 6.62E-32 |
| Ddit4 | -0.606 | 4.20E-20 |
| Nid2 | -0.589 | 1.92E-23 |
| Gm49759 | -0.582 | 5.85E-18 |
| Hk2 | -0.572 | 2.74E-18 |
| Slc2a3 | -0.564 | 9.35E-27 |
| Igfbp2 | -0.532 | 1.93E-17 |
| Dock6 | -0.514 | 7.67E-19 |
| Kdm5b | -0.430 | 1.07E-17 |
| Pgam1 | -0.387 | 2.19E-19 |

| DahA 10uM \| top15 upregulated | | |
|---|---|---|
| gene | log2 fold change | adj pvalue |
| Cyp1a1 | 3.838 | 4.00E-45 |
| Tgfbi | 1.743 | 1.69E-255 |
| Cyp1b1 | 1.679 | 7.48E-39 |
| Nqo1 | 1.634 | 1.43E-86 |
| Adh7 | 1.390 | 6.89E-44 |
| Dhrs1 | 0.944 | 4.13E-47 |
| Plin2 | 0.944 | 6.08E-40 |
| Esd | 0.751 | 9.51E-38 |
| Chd1l | 0.673 | 1.16E-45 |
| Ccng1 | 0.555 | 2.25E-58 |
| Nfe2l2 | 0.553 | 6.13E-39 |
| Ftl1-ps1 | 0.522 | 8.85E-47 |
| mt-Cytb | 0.517 | 4.64E-90 |
| mt-Nd1 | 0.499 | 5.17E-56 |
| mt-Co1 | 0.433 | 1.14E-39 |

| DahA 10uM \| top15 downregulated | | |
|---|---|---|
| gene | log2 fold change | adj pvalue |
| Nid2 | -0.554 | 2.34E-59 |
| Kmt2d | -0.519 | 1.23E-37 |
| Ago2 | -0.477 | 1.63E-37 |
| Ptprf | -0.457 | 7.00E-38 |
| Prrc2a | -0.442 | 2.25E-30 |
| Adnp | -0.440 | 6.82E-29 |
| Srcap | -0.416 | 8.61E-34 |
| Arid1a | -0.416 | 6.35E-33 |
| Ankrd11 | -0.413 | 1.95E-29 |
| Dync1h1 | -0.384 | 1.17E-30 |
| Hcfc1 | -0.379 | 4.09E-30 |
| Srrm2 | -0.366 | 3.28E-43 |
| Prrc2c | -0.352 | 4.17E-40 |
| Huwe1 | -0.333 | 1.16E-34 |
| Scd2 | -0.302 | 8.83E-31 |

### Table S3 – Top 15 upregulated and downregulated genes after 24h exposure to 1 nM and 10 µM of EDCs in the class of PCB (PCB118, PCB126, PCB138, PCB153).

**PCB118 1nM | top15 upregulated**

| gene | log2 fold change | adj pvalue |
|---|---|---|
| Car2 | 0.353 | 7.85E-08 |
| Fgf4 | 0.301 | 4.98E-06 |
| L1td1 | 0.298 | 3.54E-11 |
| Lin28a | 0.279 | 7.99E-05 |
| Zfp980 | 0.276 | 3.76E-05 |
| Klhl13 | 0.268 | 3.32E-06 |
| Hells | 0.218 | 2.56E-05 |
| Smarcad1 | 0.217 | 3.01E-05 |
| Gab1 | 0.203 | 3.01E-05 |
| G3bp2 | 0.202 | 1.94E-07 |
| Jade1 | 0.196 | 2.56E-05 |
| Npm1 | 0.185 | 1.39E-05 |
| Hsp90aa1 | 0.185 | 9.82E-06 |
| Msh6 | 0.170 | 3.25E-05 |
| Ncl | 0.165 | 5.95E-05 |

**PCB118 1nM | top15 downregulated**

| gene | log2 fold change | adj pvalue |
|---|---|---|
| Zxda | -30.000 | 1.78E-13 |
| Gm7334 | -30.000 | 1.78E-13 |
| Selenbp1 | -0.943 | 2.56E-05 |
| Trp63 | -0.818 | 2.56E-05 |
| Upk1b | -0.597 | 3.39E-06 |
| Plagl1 | -0.577 | 1.28E-07 |
| Col2a1 | -0.553 | 5.26E-06 |
| Evpl | -0.553 | 1.31E-05 |
| Nnat | -0.533 | 1.69E-06 |
| Fbn2 | -0.445 | 8.26E-05 |
| Erbb2 | -0.403 | 8.50E-05 |
| Grb10 | -0.386 | 1.32E-05 |
| H1f0 | -0.343 | 0.000115317 |
| Rsrp1 | -0.254 | 3.28E-07 |
| Scd2 | -0.200 | 1.94E-07 |

**PCB118 10uM | top15 upregulated**

| gene | log2 fold change | adj pvalue |
|---|---|---|
| Neat1 | 0.325 | 1.46E-10 |
| Fgf4 | 0.253 | 2.01E-03 |
| Tet1 | 0.215 | 4.51E-04 |
| Slc39a14 | 0.205 | 5.88E-03 |

**PCB118 10uM | top15 downregulated**

| gene | log2 fold change | adj pvalue |
|---|---|---|
| Zxda | -30.000 | 5.57E-12 |
| Gm5128 | -1.385 | 6.79E-07 |
| Ppp1ccb | -0.850 | 4.76E-04 |
| Stra8 | -0.578 | 4.70E-08 |
| Dazl | -0.437 | 4.76E-04 |
| Dmrt1 | -0.402 | 1.01E-03 |
| Plagl1 | -0.345 | 5.88E-03 |

**PCB126 1nM | top15 upregulated**

| gene | log2 fold change | adj pvalue |
|---|---|---|
| 4933427D14Rik | 0.656 | 7.30E-03 |
| mt-Rnr2 | 0.490 | 1.59E-05 |
| Tg | 0.276 | 1.07E-06 |
| Car2 | 0.219 | 1.98E-03 |

**PCB126 1nM | top15 downregulated**

| gene | log2 fold change | adj pvalue |
|---|---|---|
| Zxda | -30.000 | 2.04E-13 |
| Gm28037 | -30.000 | 2.04E-13 |
| 2300002M23Rik | -1.155 | 2.18E-03 |
| Gabrp | -0.981 | 2.02E-05 |
| Trp63 | -0.747 | 3.35E-05 |
| Upk1b | -0.709 | 2.93E-06 |
| 4933427D14Rik | -0.601 | 1.70E-03 |
| Selenbp1 | -0.508 | 4.11E-05 |
| Evpl | -0.499 | 2.17E-04 |
| Dazl | -0.497 | 2.02E-05 |
| Tacstd2 | -0.497 | 1.98E-03 |
| Mal | -0.420 | 1.12E-03 |
| Nnat | -0.397 | 5.35E-04 |
| Grb10 | -0.326 | 1.60E-03 |
| Rsrp1 | -0.290 | 6.30E-12 |

**PCB126 10uM | top15 upregulated**

| gene | log2 fold change | adj pvalue |
|---|---|---|
| Cyp1a1 | 4.962 | 1.45E-17 |
| Nqo1 | 0.969 | 1.60E-16 |
| Stmn2 | 0.611 | 7.07E-25 |
| Nr0b1 | 0.597 | 9.82E-14 |
| Klhl13 | 0.431 | 4.71E-25 |
| Psmc4 | 0.382 | 2.23E-14 |
| Ftl1-ps1 | 0.359 | 1.84E-24 |
| Tdh | 0.344 | 1.76E-15 |
| Hsp90aa1 | 0.334 | 1.05E-20 |
| Ftl2-ps | 0.334 | 2.67E-17 |
| Heatr1 | 0.331 | 7.35E-15 |
| Slc25a5 | 0.316 | 1.98E-18 |
| Odc1 | 0.310 | 6.49E-14 |
| Hspd1 | 0.295 | 7.74E-15 |
| Gm15459 | 0.255 | 1.69E-14 |

**PCB126 10uM | top15 downregulated**

| gene | log2 fold change | adj pvalue |
|---|---|---|
| Evpl | -1.018 | 7.50E-20 |
| Loxl2 | -0.902 | 2.38E-28 |
| Egln1 | -0.822 | 7.13E-29 |
| Col2a1 | -0.788 | 1.59E-19 |
| Ephb2 | -0.605 | 5.62E-17 |
| Notch3 | -0.536 | 2.82E-21 |
| Igf2bp2 | -0.529 | 5.91E-30 |
| P4ha1 | -0.528 | 1.56E-18 |
| Lama5 | -0.519 | 5.07E-24 |
| Plod2 | -0.481 | 2.04E-20 |
| Glg1 | -0.459 | 3.27E-24 |
| Clstn1 | -0.413 | 6.73E-20 |
| Nid2 | -0.389 | 6.96E-23 |
| Scd2 | -0.385 | 1.13E-31 |
| Ptprf | -0.367 | 2.82E-21 |

**PCB138 1nM | top15 upregulated**

| gene | log2 fold change | adj pvalue |
|---|---|---|
| mt-Rnr2 | 0.487 | 1.49E-03 |
| Slc5a5 | 0.421 | 1.55E-03 |
| Ccne1 | 0.220 | 3.33E-03 |
| Gab1 | 0.189 | 1.56E-04 |
| Rrn3 | 0.178 | 1.57E-03 |
| Ftl1-ps1 | 0.174 | 1.82E-03 |
| Mdh2 | 0.171 | 3.71E-03 |
| Mcm2 | 0.168 | 3.71E-03 |
| Nolc1 | 0.156 | 4.25E-03 |
| Tuba1c | 0.149 | 2.98E-03 |
| Eif3b | 0.144 | 3.38E-03 |
| Lncenc1 | 0.143 | 8.04E-03 |
| G3bp2 | 0.142 | 2.05E-03 |
| Odc1 | 0.141 | 3.71E-03 |
| Actb | 0.136 | 8.31E-03 |

**PCB138 1nM | top15 downregulated**

| gene | log2 fold change | adj pvalue |
|---|---|---|
| Gm26243 | -7.525 | 2.28E-05 |
| Gm24978 | -7.525 | 2.28E-05 |
| 2610528A11Rik | -1.738 | 3.65E-04 |
| Flg | -2.473 | 9.06E-05 |
| Gm10443 | -1.546 | 1.16E-10 |
| Upk1b | -0.608 | 6.86E-05 |
| Gm6166 | -0.532 | 2.10E-04 |
| Nnat | -0.474 | 9.06E-05 |
| Col2a1 | -0.446 | 6.86E-05 |
| Fabp5 | -0.436 | 4.32E-05 |
| Plagl1 | -0.408 | 1.24E-04 |
| Egln1 | -0.376 | 2.10E-04 |
| P4ha1 | -0.344 | 4.78E-05 |
| Plod2 | -0.284 | 1.56E-04 |
| Rsrp1 | -0.220 | 2.43E-05 |

**PCB138 10uM | top15 upregulated**

| gene | log2 fold change | adj pvalue |
|---|---|---|
| Gm37018 | 1.351 | 9.69E-03 |
| mt-Rnr2 | 0.602 | 1.14E-05 |
| Mmp9 | 0.559 | 8.90E-03 |
| Col6a3 | 0.538 | 1.13E-04 |
| Malat1 | 0.418 | 8.68E-03 |
| Neat1 | 0.407 | 1.93E-13 |
| Tg | 0.354 | 3.29E-09 |
| Tnc | 0.310 | 1.57E-03 |
| Tpo | 0.262 | 8.90E-03 |

**PCB138 10uM | top15 downregulated**

| gene | log2 fold change | adj pvalue |
|---|---|---|
| Stra8 | -0.719 | 9.50E-12 |
| Tnfrsf19 | -0.465 | 9.63E-03 |
| Dmrt1 | -0.398 | 1.21E-03 |
| Kit | -0.298 | 4.04E-05 |
| Gm10222 | -0.293 | 2.16E-03 |
| Alpl | -0.288 | 2.62E-05 |
| Rsrp1 | -0.282 | 3.29E-09 |
| Mdk | -0.274 | 1.23E-03 |
| Slc2a3 | -0.248 | 2.79E-04 |
| Tfrc | -0.178 | 1.59E-03 |

**PCB153 1nM | top15 upregulated**

| gene | log2 fold change | adj pvalue |
|---|---|---|
| mt-Rnr2 | 0.771 | 4.27E-11 |
| Clca3b | 0.698 | 2.60E-03 |
| Atg9b | 0.449 | 1.36E-03 |
| Neat1 | 0.410 | 2.12E-17 |
| Elfn1 | 0.404 | 3.60E-03 |
| Tg | 0.398 | 1.25E-03 |
| Col3a1 | 0.387 | 5.07E-04 |
| Col1a2 | 0.372 | 3.81E-03 |
| Nkx2-1 | 0.327 | 1.78E-05 |
| Tpo | 0.306 | 3.49E-05 |
| Grem1 | 0.287 | 2.91E-03 |
| Cd44 | 0.278 | 1.27E-03 |
| Ltbp1 | 0.274 | 1.25E-03 |
| Ndst1 | 0.232 | 1.25E-03 |
| Slc5a3 | 0.216 | 3.78E-03 |

**PCB153 1nM | top15 downregulated**

| gene | log2 fold change | adj pvalue |
|---|---|---|
| Gm7347 | -19.218 | 3.10E-04 |
| Gm14917 | -19.218 | 3.10E-04 |
| Gm10263 | -19.096 | 4.98E-05 |
| Gm18182 | -19.096 | 4.98E-05 |
| Rsrp1 | -0.367 | 3.21E-19 |
| Ero1a | -0.326 | 4.98E-05 |
| Slc2a3 | -0.303 | 1.62E-05 |
| Tfrc | -0.303 | 1.78E-05 |
| Nid2 | -0.287 | 4.98E-05 |
| Bsg | -0.241 | 4.98E-05 |
| Hmgcr | -0.238 | 4.98E-05 |
| Fads2 | -0.237 | 4.98E-05 |
| Fads1 | -0.233 | 3.10E-04 |
| Ldlr | -0.231 | 1.03E-04 |
| Scd2 | -0.211 | 4.49E-07 |

**PCB153 10uM | top15 upregulated**

| gene | log2 fold change | adj pvalue |
|---|---|---|
| Chac1 | 1.051 | 1.58E-12 |
| mt-Rnr2 | 0.917 | 4.84E-12 |
| Pde7b | 0.717 | 1.21E-09 |
| Txnip | 0.542 | 7.15E-28 |
| Clip4 | 0.456 | 5.50E-11 |
| St3gal1 | 0.431 | 1.23E-09 |
| Tpo | 0.424 | 2.31E-10 |
| Apoe | 0.390 | 4.60E-17 |
| Ctsa | 0.328 | 9.91E-10 |
| Mt2 | 0.316 | 4.65E-11 |
| Asns | 0.314 | 4.65E-11 |
| Mt1 | 0.305 | 6.86E-13 |
| Sars | 0.292 | 1.61E-11 |
| Ftl1-ps1 | 0.274 | 6.86E-13 |
| Ftl2-ps | 0.266 | 3.20E-10 |

**PCB153 10uM | top15 downregulated**

| gene | log2 fold change | adj pvalue |
|---|---|---|
| Ldlr | -0.447 | 6.15E-20 |
| Mat2a | -0.436 | 1.54E-21 |
| Scd2 | -0.425 | 2.53E-24 |
| Hsph1 | -0.414 | 2.10E-20 |
| Rsrp1 | -0.404 | 5.34E-24 |
| Bend3 | -0.392 | 2.30E-10 |
| Hspa5 | -0.378 | 9.19E-14 |
| Fat1 | -0.365 | 7.15E-18 |
| Fads2 | -0.355 | 2.98E-11 |
| Tfrc | -0.312 | 2.80E-11 |
| Hmgcr | -0.311 | 1.60E-10 |
| Dhcr24 | -0.306 | 5.50E-11 |
| Plxna1 | -0.304 | 7.08E-11 |
| Srrt | -0.288 | 1.08E-10 |
| Fasn | -0.272 | 4.15E-13 |

**Table S4 – Identification of DEGs due to 24h EDC exposure (*P* < 0.01) related to synthesis, metabolism, and secretion of thyroid hormones.**

| | | 10 µM dose | | | | 1 nM dose | | | |
|---|---|---|---|---|---|---|---|---|---|
| **DEGs** | **Gene description** | **PAH** | **PCB** | **Phthal** | **OPFR** | **PAH** | **PCB** | **Phthal** | **OPFR** |
| Dio1 | Deiodinase, iodothyronine, type 1 | BkF BaP | | DnOP | BADP | BkF DahA | | DINP | BADP |
| Duoxa2 | dual oxidase maturation factor 2 | BaP | | | | | | | |
| Foxe1 | forkhead box E1 | | | | DMMP | | | | |
| Mct8 | Monocarboxylate Transporter 8 | BkF DahA | | | | BKF | | | |
| Nis | Sodium Iodide Symporter | BkF DahA | | | DMMP | | PCB138 | | |
| Nkx2-1 | NK2 homeobox 1 | | PCB153 | DEHP | | | PCB153 | DEHP | |
| Pax8 | paired box 8 | | | | | | PCB153 | | |
| Tg | thyroglobulin | | PCB138 PCB153 | | | BkF | PCB126 PCB153 | DEHP | |
| Tpo | thyroid peroxidase | BkF DahA | PCB138 | DEHP DIDP | DMMP | | PCB153 | DEHP | DMMP |
| Trh | thyrotropin releasing hormone | BkF BaP | | DEHP DINP | | BaP BaA DahA | | DEHP | |
| Tshr | thyroid stimulating hormone receptor | BaP BkF | PCB153 | | | | | | |

*Upregulated, blue. Downregulated, red.*

**Table S5 – Identification of EDCs that caused significant (*P* < 0.01) dysregulation of expression of key thyroid genes for synthesis, metabolism, and secretion of thyroid hormones after 24h exposure.**

| Class | EDC | 10 μM dosing | 1 μM dosing |
|---|---|---|---|
| PAH | BaA | | Trh |
| | BAP | Dio1, Duoxa2, Trh, Tshr | Trh |
| | BKF | Dio1, Mct8, Nis, Tpo, Trh, Tshr | Dio1, Mct8, Tg |
| | DAHA | Mct8, Nis, Tpo | Dio1, Trh |
| PCBs | PCB153 | Nkx2-1, Tg, Tpo, Tshr | Nkx2-1, Pax8, Tg, Tpo |
| | PCB138 | Tg, Tpo | Nis |
| | PCB126 | | Tg |
| Phthalates | DEHP | Nkx2-1, Tpo, Trh | Nkx2-1, Tg, Tpo, Trh |
| | DIDP | Tpo | |
| OPFR | BADP | Dio1 | Dio1 |
| | DMMP | Foxe1, Nis, Tpo | Tpo |

**Table S6 – Top 15 upregulated and downregulated genes after 24h exposure to 1 nM and 10 µM of EDCs in the class of phthalates (DEHP, DIDP, DINP, DnOP).**

| DEHP 1nM \| top15 upregulated | | |
|---|---|---|
| gene | log2 fold change | adj pvalue |
| Prl2c3 | 0.766 | 1.07E-05 |
| Ctsj | 0.746 | 6.43E-11 |
| Lefty1 | 0.478 | 1.22E-15 |
| Lgals1 | 0.391 | 3.52E-07 |
| Krt18 | 0.310 | 2.95E-09 |
| Krt8 | 0.274 | 2.04E-09 |
| Rbm3 | 0.269 | 1.73E-07 |
| Eif4a-ps4 | 0.215 | 2.20E-09 |
| Tuba1a | 0.180 | 6.17E-08 |
| Rpsa | 0.174 | 6.18E-10 |
| Gm11808 | 0.172 | 1.63E-05 |
| Ftl2-ps | 0.159 | 1.80E-06 |
| Rpl41 | 0.151 | 8.48E-07 |
| Ppia | 0.130 | 1.58E-05 |
| Npm1 | 0.129 | 5.58E-07 |

| DEHP 1nM \| top15 downregulated | | |
|---|---|---|
| gene | log2 fold change | adj pvalue |
| Gm2436 | -26.834 | 1.78E-10 |
| mt-Nd6 | -0.539 | 3.71E-15 |
| Dab2 | -0.451 | 2.25E-14 |
| Timp3 | -0.437 | 3.89E-08 |
| Gm47939 | -0.416 | 9.73E-11 |
| Gm48362 | -0.359 | 9.73E-11 |
| Igfbp5 | -0.357 | 3.82E-17 |
| Gm47761 | -0.350 | 4.77E-10 |
| Gm42047 | -0.316 | 5.65E-09 |
| Vegfa | -0.304 | 2.41E-14 |
| Foxo3 | -0.293 | 6.13E-08 |
| Hspg2 | -0.249 | 9.23E-13 |
| Dlc1 | -0.241 | 1.16E-07 |
| Ago2 | -0.227 | 6.08E-11 |
| Mdm4 | -0.200 | 8.33E-08 |

| DEHP 10uM \| top15 upregulated | | |
|---|---|---|
| gene | log2 fold change | adj pvalue |
| Lefty1 | 0.623 | 1.10E-31 |
| Lefty2 | 0.318 | 4.45E-12 |
| Stra8 | 0.388 | 1.37E-07 |
| Spp1 | 0.282 | 5.85E-08 |
| Vim | 0.219 | 7.68E-06 |
| H3f3b | 0.171 | 1.28E-08 |
| Wdr43 | 0.169 | 2.00E-05 |
| Eif2s2 | 0.158 | 2.68E-06 |
| Cnbp | 0.153 | 2.68E-06 |
| Morf4l2 | 0.142 | 1.75E-05 |
| Rpl7 | 0.137 | 2.68E-06 |
| Rpsa | 0.131 | 1.42E-07 |
| Npm1 | 0.126 | 5.69E-09 |
| Rpl6 | 0.122 | 2.32E-06 |
| Hsp90aa1 | 0.113 | 1.99E-07 |

| DEHP 10uM \| top15 downregulated | | |
|---|---|---|
| gene | log2 fold change | adj pvalue |
| Gm2436 | -23.945 | 3.53E-08 |
| Gm24261 | -23.181 | 1.25E-07 |
| Rptn | -1.245 | 2.73E-09 |
| Lor | -1.168 | 1.56E-06 |
| Gm48362 | -0.413 | 2.39E-13 |
| Gm47939 | -0.412 | 9.79E-11 |
| Gm42047 | -0.401 | 1.66E-14 |
| Gm47761 | -0.390 | 1.48E-15 |
| Map3k1 | -0.262 | 7.68E-06 |
| Vegfa | -0.259 | 7.41E-11 |
| Leng8 | -0.212 | 2.93E-06 |
| Igfbp5 | -0.208 | 2.68E-06 |
| Atp11a | -0.206 | 7.68E-06 |
| Lrp2 | -0.206 | 1.48E-05 |
| | -0.203 | 1.25E-07 |

| DIDP 1nM \| top15 upregulated | | |
|---|---|---|
| gene | log2 fold change | adj pvalue |
| H3c14 | 7.242 | 5.42E-07 |
| Sfrp2 | 0.319 | 7.07E-03 |
| Dab2 | 0.282 | 4.81E-05 |
| Lefty1 | 0.273 | 3.85E-04 |
| Lefty2 | 0.257 | 4.88E-06 |
| Basp1 | 0.234 | 4.10E-04 |
| Amot | 0.228 | 7.39E-04 |
| Lrp2 | 0.222 | 2.41E-05 |
| Tuba1a | 0.164 | 1.91E-05 |

| DIDP 1nM \| top15 downregulated | | |
|---|---|---|
| gene | log2 fold change | adj pvalue |
| Gm2436 | -22.614 | 2.73E-06 |
| Gm42713 | -20.430 | 3.54E-05 |
| Rpl17-ps8 | -1.712 | 5.42E-07 |
| Lypd8l | -1.084 | 1.65E-03 |
| Tmprss11b | -0.883 | 1.57E-04 |
| Lbp | -0.652 | 3.52E-05 |
| Trf | -0.639 | 3.44E-05 |
| Gabrp | -0.624 | 1.67E-03 |
| Muc4 | -0.575 | 1.49E-04 |
| Lgals3bp | -0.503 | 3.54E-05 |
| Ly6a | -0.475 | 9.81E-04 |
| Adm2 | -0.436 | 1.67E-03 |
| Fabp5 | -0.425 | 1.91E-05 |
| A2m | -0.420 | 3.61E-04 |
| Ceacam1 | -0.292 | 2.28E-03 |

| DIDP 10uM \| top15 upregulated | | |
|---|---|---|
| gene | log2 fold change | adj pvalue |
| Vgf | 0.752 | 1.11E-05 |
| Lefty1 | 0.411 | 4.90E-12 |
| Peg10 | 0.365 | 5.70E-31 |
| Klhl13 | 0.346 | 4.57E-24 |
| Nup62cl | 0.339 | 1.38E-05 |
| Cldn6 | 0.332 | 1.29E-06 |
| Lefty2 | 0.323 | 1.19E-10 |
| Cldn4 | 0.314 | 7.44E-07 |
| Basp1 | 0.290 | 4.34E-10 |
| Sfmbt2 | 0.276 | 4.92E-17 |
| Tfap2c | 0.261 | 1.77E-07 |
| H19 | 0.205 | 4.06E-09 |
| Ass1 | 0.200 | 1.26E-06 |
| Tuba1a | 0.172 | 4.86E-06 |
| Glud1 | 0.152 | 3.88E-06 |

| DIDP 10uM \| top15 downregulated | | |
|---|---|---|
| gene | log2 fold change | adj pvalue |
| Gm2436 | -30.000 | 2.88E-14 |
| Gm14440 | -30.000 | 2.82E-14 |
| Sprr3 | -2.157 | 1.06E-10 |
| Ltf | -1.227 | 1.16E-32 |
| Dsg3 | -0.836 | 2.31E-08 |
| Trf | -0.769 | 7.37E-08 |
| Dazl | -0.659 | 1.72E-15 |
| Sohlh2 | -0.446 | 2.29E-10 |
| Otx2 | -0.442 | 2.25E-16 |
| Gm48362 | -0.419 | 1.59E-13 |
| Gm47939 | -0.403 | 2.68E-10 |
| Gm47761 | -0.397 | 1.41E-13 |
| Gm42047 | -0.375 | 1.25E-13 |
| Irf1 | -0.369 | 5.56E-08 |
| Myrf | -0.278 | 1.77E-07 |

| DINP 1nM \| top15 upregulated | | |
|---|---|---|
| gene | log2 fold change | adj pvalue |
| Lefty1 | 0.503 | 3.38E-17 |
| Hypk | 0.598 | 2.12E-11 |
| Scin | 0.527 | 6.68E-08 |
| Grem1 | 0.431 | 4.58E-07 |
| S100a6 | 0.426 | 3.38E-10 |
| Krt18 | 0.346 | 4.24E-12 |
| Thbs1 | 0.342 | 1.61E-10 |
| Cd151 | 0.332 | 6.48E-08 |
| Krt8 | 0.280 | 4.33E-09 |
| Itga3 | 0.272 | 4.47E-12 |
| Ctsl | 0.269 | 2.77E-12 |
| Ftl2-ps | 0.227 | 4.33E-09 |
| Sqstm1 | 0.213 | 1.27E-07 |
| Ftl1-ps1 | 0.212 | 2.78E-07 |
| Rpl41 | 0.171 | 3.03E-09 |

| DINP 1nM \| top15 downregulated | | |
|---|---|---|
| gene | log2 fold change | adj pvalue |
| Gm42713 | -25.784 | 2.25E-09 |
| Krt13 | -1.764 | 1.33E-08 |
| Ltf | -0.960 | 8.01E-19 |
| Muc4 | -0.783 | 7.42E-09 |
| mt-Nd6 | -0.545 | 2.05E-25 |
| Neat1 | -0.500 | 5.98E-36 |
| Otx2 | -0.430 | 2.15E-10 |
| Firre | -0.359 | 4.24E-12 |
| Vegfa | -0.308 | 1.16E-13 |
| Mdm4 | -0.303 | 2.09E-11 |
| Ubn2 | -0.288 | 1.26E-09 |
| Erdr1 | -0.259 | 1.26E-09 |
| Tet1 | -0.226 | 5.12E-09 |
| Suz12 | -0.225 | 1.26E-09 |
| Mid1 | -0.180 | 1.42E-08 |

| DINP 10uM \| top15 upregulated | | |
|---|---|---|
| gene | log2 fold change | adj pvalue |
| Lefty1 | 0.731 | 1.82E-39 |
| Ptgs2 | 0.658 | 2.28E-08 |
| Nqo1 | 0.467 | 1.00E-04 |
| Dhrs1 | 0.348 | 2.08E-05 |
| Plin2 | 0.336 | 4.31E-05 |
| Grem1 | 0.328 | 5.56E-05 |
| Cldn4 | 0.304 | 7.56E-09 |
| Srxn1 | 0.283 | 1.48E-05 |
| Anxa1 | 0.274 | 3.86E-05 |
| Lefty2 | 0.264 | 5.58E-08 |
| Trh | 0.243 | 1.18E-05 |
| Tsc22d1 | 0.198 | 3.71E-06 |
| mt-Nd4 | 0.194 | 3.86E-05 |
| Ftl1-ps1 | 0.157 | 6.99E-05 |
| H3f3b | 0.133 | 7.72E-05 |

| DINP 10uM \| top15 downregulated | | |
|---|---|---|
| gene | log2 fold change | adj pvalue |
| Gm2436 | -29.068 | 1.40E-12 |
| Amy2a3 | -26.438 | 3.21E-10 |
| Gm42713 | -24.162 | 2.28E-08 |
| Klk14 | -1.772 | 5.11E-08 |
| Trf | -0.853 | 1.60E-09 |
| Dazl | -0.485 | 5.52E-08 |
| Gm47939 | -0.460 | 5.99E-14 |
| Pfkfb3 | -0.409 | 2.28E-08 |
| Gm42047 | -0.380 | 7.82E-13 |
| Dab2 | -0.376 | 4.39E-11 |
| Otx2 | -0.375 | 8.78E-07 |
| Gm47761 | -0.372 | 4.39E-11 |
| Gm48362 | -0.355 | 1.92E-09 |
| Lrp2 | -0.263 | 5.58E-08 |
| Rbpj | -0.200 | 1.68E-07 |

| DnOP 1nM \| top15 upregulated | | |
|---|---|---|
| gene | log2 fold change | adj pvalue |
| Col17a1 | 0.532 | 2.18E-05 |
| Gata2 | 0.432 | 6.94E-03 |
| Krt17 | 0.429 | 8.04E-03 |
| Tinagl1 | 0.294 | 1.12E-04 |
| Sfn | 0.272 | 1.88E-03 |
| Jup | 0.195 | 6.94E-03 |
| Krt18 | 0.193 | 1.88E-03 |
| Krt8 | 0.170 | 4.04E-03 |
| Gnas | 0.140 | 7.63E-03 |
| Dag1 | 0.134 | 8.04E-03 |

| DnOP 1nM \| top15 downregulated | | |
|---|---|---|
| gene | log2 fold change | adj pvalue |
| Gm2436 | -25.793 | 9.90E-08 |
| mt-Nd6 | -0.352 | 2.10E-05 |
| Lncenc1 | -0.143 | 5.69E-03 |

| DnOP 10uM \| top15 upregulated | | |
|---|---|---|
| gene | log2 fold change | adj pvalue |
| Lefty1 | 0.501 | 1.01E-17 |
| Lefty2 | 0.260 | 4.44E-07 |
| Dio1 | 0.579 | 5.76E-04 |
| mt-Nd2 | 0.380 | 3.50E-04 |
| mt-Nd1 | 0.359 | 1.40E-03 |
| mt-Cytb | 0.336 | 9.26E-05 |
| mt-Nd4 | 0.309 | 4.55E-05 |
| mt-Nd5 | 0.309 | 4.55E-05 |
| Grem1 | 0.289 | 2.17E-03 |
| Cldn4 | 0.251 | 2.06E-04 |
| mt-Co1 | 0.242 | 1.04E-03 |
| Spp1 | 0.206 | 1.55E-03 |
| Krt18 | 0.188 | 1.60E-03 |
| Ccng1 | 0.171 | 2.15E-04 |
| Ftl1-ps1 | 0.158 | 2.48E-03 |

| DnOP 10uM \| top15 downregulated | | |
|---|---|---|
| gene | log2 fold change | adj pvalue |
| Gm2436 | -23.693 | 2.84E-07 |
| Gm42713 | -22.173 | 1.53E-06 |
| Psenen-ps | -19.472 | 9.26E-05 |
| Fthl17b | -17.939 | 2.00E-08 |
| Gm14552 | -17.331 | 9.23E-06 |
| Gm15785 | -16.759 | 3.53E-04 |
| Hrnr | -2.281 | 7.48E-09 |
| Muc4 | -0.686 | 1.88E-10 |
| Pfkfb3 | -0.403 | 4.76E-06 |
| Dnmt3l | -0.329 | 2.06E-04 |
| Otx2 | -0.312 | 1.26E-07 |
| G2e3 | -0.307 | 4.56E-04 |
| Irf1 | -0.280 | 7.13E-04 |
| Otg | -0.190 | 1.04E-03 |
| Vegfa | -0.185 | 5.79E-04 |

**Table S7 – Top 15 upregulated and downregulated genes after 24h exposure to 1 nM and 10 µM of EDCs in the class of OPFRs (BADP, DMMP, TDCPP, TPP).**

| BADP 1nM \| top15 upregulated | | |
|---|---|---|
| gene | log2 fold change | adj pvalue |
| mt-Rnr2 | 0.910 | 1.11E-20 |
| Cst6 | 0.604 | 2.78E-21 |
| H1f2 | 0.602 | 1.08E-17 |
| Prrg4 | 0.468 | 1.78E-09 |
| Ptgs2 | 0.448 | 2.74E-09 |
| Akr1b3 | 0.407 | 2.32E-11 |
| Areg | 0.402 | 3.24E-11 |
| mt-Nd2 | 0.401 | 2.52E-17 |
| Anxa1 | 0.397 | 1.61E-11 |
| Cnmd | 0.382 | 6.57E-12 |
| Scin | 0.377 | 1.40E-09 |
| mt-Nd1 | 0.334 | 4.30E-18 |
| Akap12 | 0.325 | 1.75E-08 |
| mt-Rnr1 | 0.320 | 1.15E-08 |
| Thbs1 | 0.272 | 6.94E-12 |

| BADP 1nM \| top15 downregulated | | |
|---|---|---|
| gene | log2 fold change | adj pvalue |
| Gm2436 | -26.834 | 1.78E-10 |
| Sftpb | -0.932 | 7.05E-25 |
| Acta2 | -0.644 | 1.66E-13 |
| mt-Nd6 | -0.539 | 3.71E-15 |
| Rsrp1 | -0.477 | 4.83E-28 |
| Dab2 | -0.451 | 2.25E-14 |
| Gm47939 | -0.416 | 9.73E-11 |
| Gm48362 | -0.359 | 9.73E-11 |
| Igfbp5 | -0.357 | 3.82E-17 |
| Gm47761 | -0.350 | 4.77E-10 |
| Clk1 | -0.345 | 5.50E-11 |
| Gm42047 | -0.316 | 5.65E-09 |
| Vegfa | -0.304 | 2.41E-14 |
| Hspg2 | -0.249 | 9.23E-13 |
| Ago2 | -0.227 | 6.08E-11 |

| BADP 10uM \| top15 upregulated | | |
|---|---|---|
| gene | log2 fold change | adj pvalue |
| Gm21742 | 0.973 | 2.71E-08 |
| Ankfn1 | 0.968 | 1.75E-06 |
| Mlph | 0.477 | 2.71E-08 |
| Slc26a7 | 0.391 | 4.18E-07 |
| Nrxn2 | 0.390 | 1.43E-09 |
| Dio1 | 0.387 | 4.64E-09 |
| Raver2 | 0.361 | 6.02E-10 |
| Chchd10 | 0.318 | 5.53E-11 |
| Srxn1 | 0.308 | 8.80E-11 |
| Dap | 0.298 | 6.10E-07 |
| Papss2 | 0.289 | 6.12E-06 |
| Ftl2-ps | 0.286 | 7.37E-08 |
| Ftl1-ps1 | 0.282 | 1.43E-09 |
| Sgpl1 | 0.268 | 1.99E-06 |
| Slc43a2 | 0.234 | 2.31E-06 |

| BADP 10uM \| top15 downregulated | | |
|---|---|---|
| gene | log2 fold change | adj pvalue |
| Gm7336 | -30.000 | 1.02E-12 |
| Rab11b-ps2 | -21.448 | 4.71E-06 |
| Mup15 | -19.567 | 1.09E-06 |
| 6330415B21Rik | -19.361 | 1.31E-05 |
| Gm28516 | -19.016 | 1.25E-04 |
| Gm48866 | -19.016 | 1.25E-04 |
| Gm5424 | -19.004 | 1.25E-04 |
| Gm16479 | -19.004 | 1.25E-04 |
| Oas1e | -16.525 | 2.30E-05 |
| Gm48334 | -16.436 | 8.52E-06 |
| Gm5128 | -4.875 | 4.67E-06 |
| Wfikkn2 | -0.503 | 4.40E-08 |
| Lox | -0.417 | 3.89E-06 |
| Lbh | -0.324 | 1.15E-04 |
| Tgfbi | -0.302 | 4.57E-09 |

| DMMP 1nM \| top15 upregulated | | |
|---|---|---|
| gene | log2 fold change | adj pvalue |
| Krt20 | 0.930 | 9.21E-04 |
| Fmo5 | 0.759 | 3.06E-04 |
| Cyp2s1 | 0.677 | 7.09E-03 |
| Acer2 | 0.616 | 4.09E-03 |
| H1f10 | 0.301 | 5.30E-03 |
| Snx22 | 0.251 | 5.86E-03 |
| Akap12 | 0.250 | 7.72E-06 |
| Cpxm2 | 0.235 | 5.11E-03 |
| Susd2 | 0.223 | 8.07E-03 |
| Itga3 | 0.197 | 8.72E-03 |

| DMMP 1nM \| top15 downregulated | | |
|---|---|---|
| gene | log2 fold change | adj pvalue |
| Gm7334 | -30.000 | 3.31E-13 |
| Gm38317 | -28.191 | 3.12E-11 |
| Rab11b-ps2 | -27.474 | 1.05E-10 |
| A930009A15Rik | -25.984 | 3.31E-13 |
| Gm48943 | -25.794 | 1.80E-11 |
| Mup15 | -25.646 | 3.22E-11 |
| 6330415B21Rik | -25.449 | 1.73E-09 |
| Gm17045 | -25.091 | 1.34E-08 |
| 4930474N09Rik | -25.091 | 1.34E-08 |
| Oas1e | -19.965 | 9.37E-08 |
| Gm20751 | -15.697 | 1.44E-06 |
| Cbr2 | -0.592 | 6.55E-05 |
| Gm5128 | -5.340 | 9.89E-05 |
| Rbp1 | -0.290 | 4.54E-08 |
| Elovl6 | -0.227 | 6.25E-05 |

| DMMP 10uM \| top15 upregulated | | |
|---|---|---|
| gene | log2 fold change | adj pvalue |
| Susd2 | 0.506 | 1.23E-20 |
| Tgfbi | 0.383 | 1.12E-18 |
| Filip1l | 0.381 | 3.09E-06 |
| Cpxm2 | 0.356 | 8.70E-09 |
| Cldn6 | 0.349 | 6.54E-07 |
| Fgf1 | 0.334 | 1.06E-06 |
| Snx22 | 0.330 | 6.24E-08 |
| Col4a3 | 0.319 | 3.09E-06 |
| Cplx2 | 0.315 | 3.43E-10 |
| Thbs1 | 0.293 | 1.40E-08 |
| Gm29216 | 0.272 | 6.40E-06 |
| Cdh16 | 0.263 | 1.04E-05 |
| Msn | 0.235 | 1.38E-07 |
| Foxe1 | 0.216 | 2.64E-06 |
| Krt7 | 0.191 | 3.09E-06 |

| DMMP 10uM \| top15 downregulated | | |
|---|---|---|
| gene | log2 fold change | adj pvalue |
| Mup2 | -27.617 | 3.43E-10 |
| Gm42713 | -26.246 | 5.12E-09 |
| Ak7 | -0.969 | 2.64E-06 |
| Zfp981 | -0.775 | 4.71E-08 |
| Zfp600 | -0.501 | 1.38E-07 |
| Myrf | -0.478 | 5.61E-06 |
| Igfbp2 | -0.419 | 8.71E-07 |
| Rbp1 | -0.404 | 1.45E-18 |
| Tpo | -0.378 | 3.62E-12 |
| Mest | -0.376 | 2.86E-15 |
| Nptx1 | -0.297 | 6.40E-06 |
| Elovl6 | -0.282 | 7.90E-13 |
| Stk40 | -0.277 | 9.71E-06 |
| Slc5a5 | -0.263 | 6.76E-06 |
| Snhg14 | -0.670 | 9.71E-06 |

| TDCPP 1nM \| top15 upregulated | | |
|---|---|---|
| gene | log2 fold change | adj pvalue |
| Gsdmc3 | 2.006 | 8.70E-06 |
| Tgm3 | 1.846 | 8.06E-06 |
| Gm3776 | 0.953 | 8.06E-06 |
| Abca12 | 0.891 | 7.64E-04 |
| Ppbp | 0.743 | 1.34E-11 |
| Krt80 | 0.641 | 3.84E-05 |
| Psca | 0.515 | 4.12E-10 |
| Areg | 0.439 | 1.30E-06 |
| Tnfaip2 | 0.405 | 4.80E-06 |
| Mal | 0.394 | 1.14E-10 |
| Lad1 | 0.369 | 2.93E-05 |
| Sfn | 0.362 | 1.06E-04 |
| Itgb4 | 0.338 | 5.88E-05 |
| Spns2 | 0.271 | 7.17E-04 |
| Ccn2 | 0.230 | 1.15E-04 |

| TDCPP 1nM \| top15 downregulated | | |
|---|---|---|
| gene | log2 fold change | adj pvalue |
| Psenen-ps | -30.000 | 1.91E-13 |
| Gm38317 | -30.000 | 1.63E-13 |
| Gm49359 | -30.000 | 1.25E-13 |
| Gm7334 | -30.000 | 9.07E-14 |
| 6330415B21Rik | -30.000 | 7.20E-14 |
| Gm26630 | -30.000 | 7.20E-14 |
| Gm14808 | -30.000 | 5.94E-14 |
| Mup15 | -30.000 | 7.21E-16 |
| Gm49496 | -30.000 | 7.21E-16 |
| Marco | -30.000 | 1.18E-51 |
| Gm37058 | -29.997 | 4.91E-27 |
| Gm14117 | -29.984 | 1.08E-33 |
| Gm3892 | -26.756 | 2.76E-18 |
| 4632433K11Rik | -26.556 | 7.62E-17 |
| Gm43847 | -24 | 1.66E-14 |

| TDCPP 10uM \| top15 upregulated | | |
|---|---|---|
| gene | log2 fold change | adj pvalue |
| Lce3c | 2.408 | 1.64E-05 |
| Klk6 | 1.282 | 6.64E-04 |
| Lcn2 | 0.943 | 1.95E-03 |
| Krtdap | 0.838 | 9.48E-05 |
| Psca | 0.501 | 9.05E-08 |
| Egr1 | 0.444 | 3.03E-05 |
| Plet1 | 0.374 | 3.03E-05 |
| Ccn2 | 0.301 | 9.72E-05 |
| Susd2 | 0.260 | 7.63E-04 |
| Myo5b | 0.255 | 1.32E-03 |
| Thbs1 | 0.248 | 2.73E-04 |
| Ucp2 | 0.205 | 4.43E-04 |
| Jund | 0.189 | 2.20E-04 |
| Ncor2 | 0.183 | 6.08E-04 |
| Myh9 | 0.154 | 1.27E-03 |

| TDCPP 10uM \| top15 downregulated | | |
|---|---|---|
| gene | log2 fold change | adj pvalue |
| Gm5424 | -30.000 | 2.68E-10 |
| Gm7336 | -30.000 | 7.11E-11 |
| Mup15 | -30.000 | 4.04E-11 |
| Gm49496 | -30.000 | 4.04E-11 |
| 6330415B21Rik | -29.868 | 2.25E-13 |
| 4930474N09Rik | -29.513 | 4.95E-12 |
| Gm42713 | -28.719 | 6.46E-10 |
| Gm38317 | -27.492 | 6.66E-09 |
| Gm13597 | -7.378 | 9.72E-05 |
| Zfp534 | -1.026 | 1.53E-05 |
| Gm13230 | -0.979 | 3.41E-05 |
| Zfp979 | -0.873 | 2.46E-04 |
| Zfp990 | -0.667 | 3.41E-05 |
| Zfp992 | -0.572 | 2.40E-04 |
| Tra2a | -0.240 | 2.40E-04 |

| TPP 1nM \| top15 upregulated | | |
|---|---|---|
| gene | log2 fold change | adj pvalue |
| Zxda | 25.531 | 4.33E-10 |
| Cst6 | 0.525 | 7.61E-09 |
| Col11a2 | 0.523 | 4.89E-04 |
| Hypk | 0.499 | 4.88E-05 |
| Cldn6 | 0.372 | 2.60E-05 |
| Areg | 0.330 | 8.46E-03 |
| Scin | 0.316 | 2.27E-04 |
| Cnmd | 0.301 | 5.93E-04 |
| Ldha | 0.266 | 1.34E-07 |
| Eno1b | 0.262 | 2.22E-04 |
| Akap12 | 0.249 | 3.97E-04 |
| Eno1 | 0.236 | 1.69E-03 |
| Cpxm2 | 0.232 | 8.46E-03 |
| Pkm | 0.229 | 4.43E-04 |
| Tpi1 | 0.203 | 1.28E-03 |

| TPP 1nM \| top15 downregulated | | |
|---|---|---|
| gene | log2 fold change | adj pvalue |
| Gm13486 | -30.000 | 6.13E-14 |
| Gm17045 | -30.000 | 6.13E-14 |
| Gm4938 | -30.000 | 6.13E-14 |
| 4930474N09Rik | -30.000 | 6.13E-14 |
| Gm16479 | -30.000 | 6.13E-14 |
| Rab11b-ps2 | -30.000 | 4.70E-14 |
| Gm1972 | -30.000 | 4.70E-14 |
| Gm8741 | -30.000 | 4.70E-14 |
| Gm42713 | -30.000 | 3.50E-14 |
| 6330415B21Rik | -30.000 | 8.53E-15 |
| Gm26630 | -30.000 | 8.53E-15 |
| Gm11420 | -30.000 | 8.53E-15 |
| Mup15 | -30.000 | 1.26E-16 |
| Gm49496 | -30.000 | 1.26E-16 |
| Gm9347 | -21.400 | 8.56E-16 |

| TPP 10uM \| top15 upregulated | | |
|---|---|---|
| gene | log2 fold change | adj pvalue |
| Gsdmc3 | 2.727 | 1.50E-10 |
| Gsdmc4 | 3.583 | 9.10E-04 |
| Klk6 | 1.550 | 3.54E-04 |
| Nipal4 | 1.155 | 9.10E-04 |
| Sprr1b | 0.920 | 2.65E-03 |
| Abca12 | 0.884 | 2.36E-03 |
| Acer2 | 0.766 | 6.96E-03 |
| Tet1 | 0.304 | 7.16E-03 |
| Akap12 | 0.303 | 1.30E-08 |
| Akr1b3 | 0.280 | 2.18E-03 |
| Osbpl3 | 0.274 | 6.63E-03 |
| Phc1 | 0.238 | 9.10E-04 |
| Itga3 | 0.232 | 8.21E-03 |
| Cldn4 | 0.222 | 1.39E-03 |
| Notch1 | 0.207 | 7.24E-03 |

| TPP 10uM \| top15 downregulated | | |
|---|---|---|
| gene | log2 fold change | adj pvalue |
| Gm37752 | -28.642 | 1.50E-10 |
| Gm38317 | -28.635 | 1.50E-10 |
| Gm8741 | -23.781 | 1.13E-06 |
| Rab11b-ps2 | -22.695 | 4.68E-06 |
| Fthl17e | -20.600 | 7.36E-05 |
| Mup15 | -20.099 | 1.53E-05 |
| Idi1-ps1 | -5.537 | 5.07E-05 |
| Dcn | -0.510 | 1.29E-04 |
| Sftpb | -0.462 | 3.47E-04 |
| Col1a1 | -0.371 | 2.11E-06 |
| Lox | -0.356 | 9.83E-05 |
| Col1a2 | -0.323 | 4.57E-04 |
| Clk1 | -0.287 | 1.08E-05 |
| Cdh11 | -0.271 | 1.10E-05 |
| Rsrp1 | -0.238 | 4.53E-06 |

**Table S8 – Top 15 upregulated and downregulated proteins after 24h exposure to EDCs at 10 µM concentration.**

| 24h BaA 10uM \| top15 upregulated | | |
|---|---|---|
| Protein | log2 fold change | adj pvalue |
| Hspg2 | 2,28 | 9,63E-04 |
| Tcof1 | 0,87 | 9,90E-04 |
| Hmbs | 0,42 | 4,42E-03 |

| 24h BaA 10uM \| top15 downregulated | | |
|---|---|---|
| Protein | log2 fold change | adj pvalue |
| Tubgcp3 | -2,23 | 9,27E-03 |
| Emc3 | -1,00 | 4,46E-03 |
| Trrap | -0,81 | 9,20E-03 |
| Ubfd1 | -0,46 | 1,84E-03 |
| Pdia5 | -0,43 | 3,22E-03 |
| Atp6v1b2 | -0,33 | 7,06E-03 |
| Vamp8 | -0,31 | 2,82E-03 |

| 24h BaP 10uM \| top15 upregulated | | |
|---|---|---|
| Protein | log2 fold change | adj pvalue |
| Slc5a5 | 2,00 | 1,12E-03 |
| Sap30 | 1,97 | 5,39E-03 |
| Ehd2 | 1,85 | 1,01E-03 |
| Slc25a35 | 1,62 | 4,65E-03 |
| Atp6v0a4 | 1,46 | 4,28E-03 |
| Poglut3 | 1,32 | 2,15E-03 |
| Tor3a | 1,15 | 4,18E-03 |
| Bphl | 1,09 | 1,08E-03 |
| Ppm1h | 1,09 | 8,04E-03 |
| Mrpl15 | 1,02 | 9,50E-03 |
| Rras | 1,01 | 7,85E-03 |
| Mrps25 | 0,98 | 9,11E-03 |
| Rps6ka5 | 0,97 | 5,05E-03 |
| Fbxo7 | 0,96 | 6,11E-03 |
| Atp6v0c | 0,94 | 9,36E-03 |

| 24h BaP 10uM \| top15 downregulated | | |
|---|---|---|
| Protein | log2 fold change | adj pvalue |
| Aqp3 | -3,39 | 6,53E-03 |
| Agr2 | -3,18 | 9,24E-03 |
| Dcun1d1 | -1,50 | 7,10E-03 |
| Zwint | -1,30 | 3,93E-03 |
| Ung | -1,08 | 9,22E-03 |
| Nop56 | -0,67 | 9,98E-03 |
| Emc3 | -0,64 | 4,49E-03 |
| Pes1 | -0,60 | 6,02E-03 |
| Rps3a | -0,41 | 9,28E-03 |

| 24h BKF 10uM \| top15 upregulated | | |
|---|---|---|
| Protein | log2 fold change | adj pvalue |
| Ttc38 | 1,03 | 5,55E-03 |
| Smpd1 | 0,96 | 4,13E-03 |
| Cryl1 | 0,94 | 9,77E-03 |
| Nqo1 | 0,93 | 1,90E-03 |
| Lsm1 | 0,87 | 1,83E-03 |
| Rad21 | 0,56 | 9,97E-03 |
| Gpx1 | 0,41 | 2,20E-03 |
| Psme1 | 0,35 | 6,12E-03 |
| Pdcd6 | 0,31 | 6,04E-03 |
| Slc25a4 | 0,30 | 2,92E-03 |

| 24h BKF 10 uM \| top15 downregulated | | |
|---|---|---|
| Protein | log2 fold change | adj pvalue |
| Cc2d1b | -1,71 | 6,18E-03 |
| Cutc | -1,26 | 2,82E-03 |
| Mbd3 | -1,17 | 3,78E-03 |
| Pdia5 | -0,49 | 2,40E-03 |
| Pum1 | -0,49 | 8,92E-03 |
| Capzb | -0,23 | 5,23E-03 |

| 24h DahA 10uM \| top15 upregulated | | |
|---|---|---|
| Protein | log2 fold change | adj pvalue |
| Ehd2 | 1,48 | 2,81E-03 |
| Hypk | 1,25 | 7,56E-03 |
| Polr1a | 0,96 | 9,43E-03 |
| Lamb1 | 0,78 | 7,07E-03 |
| Nme1 | 0,18 | 6,12E-03 |

| 24h DahA 10uM \| top15 downregulated | | |
|---|---|---|
| Protein | log2 fold change | adj pvalue |
| Fuca2 | -2,16 | 7,74E-03 |
| Gtse1 | -1,75 | 2,02E-03 |
| Fhl1 | -1,06 | 8,28E-03 |

| 24h PCB118 10uM \| top15 upregulated | | |
|---|---|---|
| Protein | log2 fold change | adj pvalue |
| Col4a2 | 3,43 | 7,79E-03 |
| Rnf20 | 3,21 | 3,09E-03 |
| Rif1 | 2,88 | 1,51E-03 |
| Lamb2 | 2,84 | 5,65E-03 |
| Cav1 | 2,60 | 6,65E-06 |
| Top2a | 2,29 | 4,59E-03 |
| Lama1 | 1,97 | 1,14E-03 |
| Nid1 | 1,89 | 9,65E-04 |
| Lamb1 | 1,81 | 1,49E-03 |
| Lamc1 | 1,60 | 4,26E-03 |
| Dnmt1 | 1,52 | 5,29E-03 |
| Nedd4l | 1,42 | 8,42E-03 |
| Chd4 | 1,41 | 4,91E-05 |
| Rrp7a | 1,31 | 8,44E-03 |
| | 0,96 | 7,67E-03 |

| 24h PCB118 10uM \| top15 downregulated | | |
|---|---|---|
| Protein | log2 fold change | adj pvalue |
| Scaf8 | -2,38 | 3,82E-03 |
| Qki | -1,70 | 1,22E-03 |
| Amt | -1,61 | 1,31E-03 |
| Bphl | -1,59 | 3,52E-03 |
| Pdcl3 | -1,46 | 2,99E-03 |
| Atad3 | -1,40 | 7,70E-03 |
| Sec23ip | -1,35 | 6,90E-03 |
| Mtco2 | -1,31 | 5,71E-03 |
| Rbpms | -1,31 | 1,70E-03 |
| Kif15 | -1,25 | 2,29E-04 |
| Tmem205 | -1,24 | 5,13E-03 |
| Igbp1 | -1,23 | 5,46E-04 |
| Naxe | -1,15 | 8,61E-03 |
| Tomm70 | -1,15 | 7,37E-03 |
| Pbdc1 | -1,13 | 2,99E-03 |

| 24h PCB126 10uM \| top15 upregulated | | |
|---|---|---|
| Protein | log2 fold change | adj pvalue |
| Hspg2 | 6,05 | 7,86E-03 |
| Col4a1 | 3,36 | 9,14E-06 |
| Lamb2 | 3,34 | 1,92E-03 |
| Col4a2 | 3,04 | 8,03E-03 |
| Rif1 | 3,04 | 5,11E-07 |
| Nid2 | 2,37 | 5,32E-04 |
| Lama1 | 2,19 | 6,81E-05 |
| Nid1 | 2,12 | 1,55E-05 |
| Lamb1 | 2,10 | 1,68E-04 |
| Lamc1 | 2,05 | 1,37E-04 |
| Tcof1 | 1,94 | 6,82E-06 |
| Pum2 | 1,83 | 1,17E-06 |
| Gpd1 | 1,76 | 7,77E-03 |
| Sall4 | 1,61 | 5,79E-03 |
| Dnmt1 | 1,59 | 4,44E-04 |

| 24h PCB126 10uM \| top15 downregulated | | |
|---|---|---|
| Protein | log2 fold change | adj pvalue |
| Plac8 | -2,23 | 4,16E-03 |
| Cavin1 | -2,01 | 2,08E-03 |
| Rbpms2 | -1,81 | 3,43E-03 |
| Klhl22 | -1,57 | 3,26E-03 |
| Pbxip1 | -1,49 | 2,32E-03 |
| Zdbf2 | -1,44 | 2,14E-03 |
| Dusp3 | -1,34 | 2,52E-03 |
| Rbpms | -1,21 | 2,67E-03 |
| Tor1aip1 | -1,11 | 6,51E-03 |
| Rida | -1,06 | 1,23E-03 |
| Cnot9 | -0,98 | 6,33E-04 |
| Pycr2 | -0,97 | 3,83E-03 |
| Uchl5 | -0,96 | 6,98E-03 |
| Me2 | -0,94 | 4,27E-03 |
| Cstf1 | -0,90 | 1,65E-03 |

| 24h PCB138 10uM \| top15 upregulated | | |
|---|---|---|
| Protein | log2 fold change | adj pvalue |
| Rif1 | 2,06 | 9,97E-04 |
| Pum2 Kiaa023 | 1,52 | 7,07E-04 |
| Lamb1 Lamb-1 | 1,43 | 5,95E-04 |
| Lama1 Lama L | 1,26 | 2,70E-03 |
| Nid1 Ent | 1,21 | 4,03E-03 |
| Lamc1 Lamb-2 | 1,17 | 2,75E-03 |
| Srxn1 Npn3 Sr | 1,15 | 1,07E-03 |
| Chd4 | 1,01 | 3,38E-03 |
| Uqcrh | 0,95 | 6,49E-03 |
| Plcg1 Plcg-1 | 0,79 | 7,18E-04 |
| FLJ45252 | 0,75 | 5,95E-03 |
| Col18a1 | 0,64 | 1,05E-03 |
| Ddx18 | 0,62 | 8,97E-03 |
| Mybbp1a P160 | 0,57 | 2,37E-04 |
| Nop56 Nol5a | 0,41 | 6,92E-03 |

| 24h PCB138 10uM \| top15 downregulated | | |
|---|---|---|
| Protein | log2 fold change | adj pvalue |
| Vps25 | -1,76 | 2,28E-03 |
| Mrpl4 | -0,87 | 2,69E-03 |
| Nup35 | -0,82 | 4,18E-03 |
| Ces1d | -0,80 | 5,02E-03 |
| Cnot9 | -0,78 | 5,77E-04 |
| Srsf10 | -0,75 | 5,77E-03 |
| Cstf1 | -0,67 | 3,47E-03 |
| Srrm2 | -0,65 | 8,93E-04 |
| Pmm2 | -0,62 | 9,75E-04 |
| Rpl6 | -0,50 | 3,58E-04 |
| Atl3 | -0,49 | 3,69E-03 |
| Cox6c | -0,48 | 5,22E-03 |
| Tg | -0,39 | 5,65E-03 |
| Camk1d | -0,38 | 2,24E-03 |
| Rpl7a | -0,37 | 6,14E-03 |

| 24h PCB153 10uM \| top15 upregulated | | |
|---|---|---|
| Protein | log2 fold change | adj pvalue |
| Lamb2 | 3,08 | Q61292 |
| Col4a1 | 2,52 | P02463 |
| Nid2 | 2,15 | O88322 |
| Lamb1 | 1,97 | P02469 |
| Lama1 | 1,97 | P19137 |
| Lamc1 | 1,87 | P02468 |
| Hypk | 1,82 | Q9CR41 |
| Nid1 | 1,77 | P10493 |
| Pum2 | 1,70 | Q80U58 |
| Git1 | 1,51 | Q68FF6 |
| Dsc2 | 1,48 | P55292 |
| Fhl1 | 1,47 | P97447 |
| Cog7 | 1,17 | Q3UM29 |
| Uhrf1 | 1,14 | Q8VDF2 |
| H1-2 | 1,08 | P15864 |

| 24h PCB153 10uM \| top15 downregulated | | |
|---|---|---|
| Protein | log2 fold change | adj pvalue |
| Flot1 | -2,15 | 1,79E-03 |
| Exoc2 | -2,04 | 5,80E-03 |
| Nme3 | -1,56 | 4,76E-03 |
| Mtco2 | -1,47 | 1,28E-03 |
| Ndufb9 | -1,12 | 3,00E-04 |
| Eif4e2 | -1,08 | 1,54E-03 |
| Rap1gds1 | -1,07 | 3,46E-03 |
| Tfip11 | -1,07 | 6,71E-03 |
| Csk | -0,96 | 4,85E-03 |
| Hdhd5 | -0,93 | 2,46E-03 |
| Mgst1 | -0,92 | 2,63E-04 |
| Samhd1 | -0,90 | 8,82E-03 |
| Uap1 | -0,90 | 1,49E-03 |
| Abhd11 | -0,89 | 3,74E-03 |
| Sco1 | -0,89 | 2,26E-03 |

| 24h DEHP 10uM \| top15 upregulated | | |
|---|---|---|
| Protein | log2 fold change | adj pvalue |
| Ppih | 2,70 | 7,12E-03 |
| Mpdu1 | 2,27 | 9,76E-03 |
| Plac8 | 2,09 | 1,91E-04 |
| Dhcr7 | 1,87 | 9,09E-03 |
| Gkn1 | 1,72 | 8,52E-03 |
| Tmem33 | 1,65 | 2,13E-03 |
| Dnmt3b | 1,43 | 5,86E-03 |
| Riok1 | 1,36 | 7,98E-03 |
| Rhot1 | 1,34 | 6,04E-03 |
| Psrc1 | 1,25 | 5,28E-03 |
| Fahd1 | 1,14 | 3,19E-03 |
| Smpd4 | 1,11 | 6,21E-03 |
| Utp3 | 1,02 | 5,27E-03 |
| Tomm40 | 0,99 | 1,70E-03 |
| Sox2 | 0,77 | 9,83E-03 |

| 24h DEHP 10uM \| top15 downregulated | | |
|---|---|---|
| Protein | log2 fold change | adj pvalue |
| Rdh13 | -2,24 | 3,77E-03 |
| Pfdn6 H2-Ke2 I | -1,23 | 2,74E-03 |
| Fkbp2 Fkbp13 | -0,84 | 5,96E-03 |
| Clta | -0,74 | 5,60E-03 |
| Tbca | -0,68 | 2,93E-03 |
| Itgb4 | -0,47 | 8,81E-04 |
| Tpm1 Tpm-1 T | -0,40 | 6,10E-03 |
| Casp7 Lice2 M | -0,39 | 9,94E-03 |
| Nqo1 Dia4 Nm | -0,36 | 4,25E-03 |
| Sod1 | -0,32 | 5,03E-03 |

| 24h DIDP 10uM \| top15 upregulated | | |
|---|---|---|
| Protein | log2 fold change | adj pvalue |
| Slc1a5 | 1,61 | 5,82E-03 |
| Ireb2 | 1,53 | 1,84E-03 |
| Fer | 1,53 | 1,46E-03 |
| Dnmt3b | 1,32 | 8,38E-03 |
| Rrp7a | 1,11 | 9,52E-03 |
| Sec61g | 1,02 | 6,25E-03 |
| Cldn3 | 1,02 | 3,36E-03 |
| Lama5 | 0,94 | 7,52E-03 |
| Iyd | 0,90 | 1,20E-03 |
| Spin1 | 0,84 | 6,06E-03 |
| Cryzl1 | 0,83 | 9,37E-03 |
| Ccnt1 | 0,76 | 2,07E-03 |
| Ssr3 | 0,68 | 5,61E-03 |
| Trmt5 | 0,58 | 6,08E-03 |
| Fxr2 | 0,57 | 8,22E-03 |

| 24h DIDP 10uM \| top15 downregulated | | |
|---|---|---|
| Protein | log2 fold change | adj pvalue |
| Phf6 Kiaa1823 | -1,90 | 1,06E-03 |
| Mtarc2 Marc2 I | -1,12 | 3,63E-03 |
| Fkbp2 Fkbp13 | -1,06 | 2,18E-03 |
| Vps35l | -0,94 | 3,94E-03 |
| Chmp7 | -0,94 | 6,32E-03 |
| Uqcrh | -0,86 | 8,50E-03 |
| Nit1 | -0,81 | 2,31E-03 |
| Ccdc91 Ggabp | -0,77 | 8,18E-03 |
| Tbca | -0,69 | 1,06E-03 |
| Clta | -0,68 | 4,46E-03 |
| Ranbp1 Htf9-a | -0,46 | 9,53E-03 |
| Heatr5a Kiaa1 | -0,24 | 6,86E-04 |
| Anxa2 Anx2 Ca | -0,23 | 2,47E-03 |
| Psma7 | -0,15 | 3,17E-03 |
| Flna Fln Fln1 | -0,13 | 2,55E-03 |

| 24h DINP 10uM \| top15 upregulated | | |
|---|---|---|
| Protein | log2 fold change | adj pvalue |
| Sec61g | 1,13 | 2,35E-03 |
| Ptpn12 | 0,91 | 1,93E-03 |
| Cth | 0,88 | 3,03E-03 |
| Tom1 | 0,81 | 6,34E-03 |
| Ndufa13 | 0,81 | 6,96E-03 |
| Eps15l1 | 0,75 | 5,10E-03 |
| Gnaq | 0,70 | 2,23E-03 |
| Cdc16 | 0,67 | 4,00E-03 |
| Bcap29 | 0,60 | 1,80E-04 |
| Exoc4 | 0,46 | 1,36E-03 |
| Rps16 | 0,41 | 6,75E-03 |
| Nup54 | 0,38 | 2,35E-03 |
| Ncs2 | 0,37 | 7,11E-03 |
| Mmut | 0,36 | 7,73E-03 |
| Yif1b | 0,33 | 1,59E-03 |

| 24h DINP 10uM \| top15 downregulated | | |
|---|---|---|
| Protein | log2 fold change | adj pvalue |
| Tbca | -1,40 | 5,86E-03 |
| Fkbp2 | -1,13 | 6,75E-03 |
| Rpl32-ps | -0,80 | 6,25E-04 |
| Ubp1 | -0,70 | 3,36E-03 |
| Prdx1 | -0,48 | 9,23E-03 |
| Ero1b | -0,45 | 6,93E-03 |
| Sod1 | -0,39 | 5,23E-03 |
| Pa2g4 | -0,30 | 3,83E-03 |
| Hspa8 | -0,20 | 5,11E-03 |

| 24h DnOP 10uM \| top15 upregulated | | |
|---|---|---|
| Protein | log2 fold change | adj pvalue |
| Ckap2 | 1,91 | 8,06E-03 |
| Gfer | 1,19 | 1,33E-04 |
| Armc10 | 1,19 | 1,13E-03 |
| Scamp2 | 0,95 | 4,86E-03 |
| Fntb | 0,92 | 6,18E-03 |
| C20orf27 | 0,84 | 5,87E-03 |
| Prim1 | 0,82 | 3,46E-03 |
| Gapvd1 | 0,79 | 5,07E-03 |
| Vps26b | 0,79 | 8,18E-03 |
| Hmg20a | 0,75 | 3,80E-04 |
| Polr2l | 0,68 | 9,86E-03 |
| Imp4 | 0,59 | 2,14E-03 |
| Dnaja1 | 0,55 | 5,28E-03 |
| Ndufb9 | 0,53 | 6,31E-03 |
| Eif1ad | 0,52 | 2,77E-03 |

| 24h DnOP 10uM \| top15 downregulated | | |
|---|---|---|
| Protein | log2 fold change | adj pvalue |
| Emc3 | -1,10 | 8,97E-03 |
| Sumf2 | -0,99 | 4,80E-03 |
| Sypl1 | -0,83 | 5,54E-03 |
| Flot1 | -0,81 | 7,35E-03 |
| Rab5b | -0,67 | 9,10E-03 |
| Abr | -0,65 | 2,66E-03 |
| P4ha1 | -0,55 | 7,72E-03 |
| Bcap29 | -0,52 | 1,10E-03 |
| Galnt2 | -0,29 | 2,51E-03 |
| Anxa2 | -0,29 | 1,99E-03 |

| 24h BADP 10uM \| top15 upregulated | | |
|---|---|---|
| Protein | log2 fold change | adj pvalue |
| Tpx2 | 2,75 | 4,53E-04 |
| Mrps11 | 2,51 | 6,89E-04 |
| Col18a1 | 2,04 | 1,48E-03 |
| Tubgcp3 | 1,95 | 6,83E-03 |
| Cetn2 | 1,68 | 9,39E-03 |
| Rela | 1,35 | 3,17E-03 |
| Chmp7 | 1,25 | 5,02E-04 |
| Scrn3 | 1,10 | 8,11E-03 |
| Prpf4b | 1,03 | 6,52E-04 |
| Mtfr1l | 0,95 | 9,87E-04 |
| Psmb4 | 0,73 | 1,04E-03 |
| Denr | 0,42 | 7,80E-03 |
| Sh2b1 | 0,33 | 7,24E-03 |
| Stx12 | 0,27 | 7,46E-03 |

| 24h BADP 10uM \| top15 downregulated | | |
|---|---|---|
| Protein | log2 fold change | adj pvalue |
| Ap3b1 | -1,44 | 2,63E-03 |
| Tmx2 | -1,31 | 8,33E-03 |
| Fads2 | -0,90 | 2,99E-03 |
| Cr1l | -0,72 | 5,69E-03 |
| Tecr | -0,60 | 1,32E-03 |
| Rpl10a | -0,43 | 7,21E-03 |
| Rpl4 | -0,42 | 4,78E-03 |
| Gcn1 | -0,36 | 3,21E-03 |
| Arg1 | -0,29 | 4,65E-03 |
| Iqgap2 | -0,28 | 2,01E-03 |
| Psmd12 | -0,23 | 3,78E-03 |
| Ap2a1 | -0,15 | 8,47E-03 |

| 24h DMMP 10uM \| top15 upregulated | | |
|---|---|---|
| Protein | log2 fold change | adj pvalue |
| Pla2g4a | 1,62 | 2,66E-04 |
| Stxbp3 | 1,62 | 1,71E-03 |
| Htt | 1,57 | 2,32E-03 |
| Rbbp6 | 1,44 | 9,34E-03 |
| H1a | 1,11 | 7,97E-03 |
| Aatf | 1,11 | 8,39E-03 |
| Abitram | 1,09 | 1,44E-05 |
| Lama5 | 1,00 | 3,61E-03 |
| Plscr3 | 0,97 | 2,31E-03 |
| Slmap | 0,83 | 4,33E-03 |
| Rtf1 | 0,62 | 1,29E-03 |
| Cav1 | 0,59 | 9,65E-03 |
| Myo1b | 0,57 | 8,05E-04 |
| Pkn1 | 0,49 | 9,68E-03 |
| Cadm4 | 0,43 | 4,26E-03 |

| 24h DMMP 10uM \| top15 downregulated | | |
|---|---|---|
| Protein | log2 fold change | adj pvalue |
| Pdk3 | -1,19 | 7,67E-03 |
| Tmem176a | -0,63 | 3,72E-03 |

| 24h TDCPP 10uM \| top15 upregulated | | |
|---|---|---|
| Protein | log2 fold change | adj pvalue |
| Rif1 | 1,39 | 4,22E-03 |
| Nes | 1,29 | 7,00E-03 |
| Commd9 | 1,09 | 6,45E-03 |
| Pum3 | 1,04 | 6,68E-03 |
| Vmp1 | 0,76 | 5,35E-03 |
| Akap9 | 0,69 | 8,66E-03 |
| Cpne1 | 0,55 | 3,07E-03 |
| Nipsnap3b | 0,47 | 1,76E-03 |
| Tubb6 | 0,47 | 3,96E-03 |
| Sord | 0,45 | 3,68E-04 |
| Stx12 | 0,31 | 9,96E-03 |
| Agps | 0,28 | 2,75E-03 |

| 24h TDCPP 10uM \| top15 downregulated | | |
|---|---|---|
| Protein | log2 fold change | adj pvalue |
| Meak7 | -2,22 | 9,82E-04 |
| Trir | -0,79 | 2,64E-03 |
| Alg2 | -0,76 | 4,92E-03 |
| Tecr | -0,41 | 9,65E-03 |
| Dvl2 | -0,30 | 2,73E-03 |
| Rab25 | -0,27 | 7,77E-03 |

| 24h TPP 10uM \| top15 upregulated | | |
|---|---|---|
| Protein | log2 fold change | adj pvalue |
| Rnf7 | 2,07 | 4,40E-03 |
| Pum3 | 1,41 | 3,26E-03 |
| Optn | 1,34 | 5,63E-04 |
| Cltb | 1,32 | 4,28E-03 |
| Arpc3 | 1,32 | 2,45E-03 |
| Paip1 | 1,30 | 6,45E-04 |
| Cwc15 | 1,08 | 9,00E-03 |
| Mtpn | 1,03 | 5,35E-03 |
| Coq5 | 0,89 | 6,07E-03 |
| Atg4b | 0,87 | 9,87E-03 |
| Psmb4 | 0,63 | 4,24E-03 |
| Cpne1 | 0,63 | 5,32E-04 |
| Erh | 0,61 | 3,44E-03 |
| Ywhaq | 0,53 | 8,66E-03 |
| Ipo4 | 0,53 | 4,52E-03 |

| 24h TPP 10uM \| top15 downregulated | | |
|---|---|---|
| Protein | log2 fold change | adj pvalue |
| Ephx2 | -2,21 | 7,96E-03 |
| Camk1 | -1,82 | 5,43E-03 |
| Cadm4 | -1,81 | 8,37E-03 |
| Znf622 | -1,31 | 7,26E-03 |
| Rmdn3 | -1,17 | 9,63E-03 |
| Golph3 | -1,14 | 6,10E-04 |
| Abcd3 | -1,14 | 5,34E-03 |
| Rpia | -1,10 | 7,75E-03 |
| Alg2 | -1,07 | 4,58E-03 |
| Bzw1 | -0,99 | 7,45E-03 |
| Adss2 | -0,95 | 8,40E-03 |
| Stxbp3 | -0,71 | 4,74E-03 |
| Tmem176a | -0,62 | 5,32E-03 |
| Aifm1 | -0,60 | 9,99E-03 |
| Nup214 | -0,58 | 9,26E-03 |

**Table S9 – Top 15 upregulated and downregulated genes after 10 days exposure to 10 µM BaP, PCB153, and DEHP.**

| BAP 10uM \| top15 upregulated | | |
|---|---|---|
| gene | log2 fold change | adj pvalue |
| C2cd4b | 0.779 | 4.60E-08 |
| Slc2a6 | 0.675 | 8.81E-07 |
| Smoc2 | 0.664 | 1.84E-08 |
| Serpinf1 | 0.645 | 1.12E-08 |
| Ppp1r1b | 0.575 | 5.64E-09 |
| Eva1b | 0.567 | 2.25E-06 |
| Mt1 | 0.553 | 1.20E-07 |
| Cebpd | 0.543 | 2.88E-08 |
| Tg | 0.543 | 1.42E-07 |
| Foxe1 | 0.517 | 4.60E-08 |
| Iyd | 0.495 | 1.43E-13 |
| Tns1 | 0.445 | 3.55E-08 |
| Adm2 | 0.442 | 3.46E-07 |
| Stk40 | 0.282 | 3.63E-07 |
| Srebf1 | 0.257 | 3.46E-07 |

| BAP 10uM \| top15 downregulated | | |
|---|---|---|
| gene | log2 fold change | adj pvalue |
| Rxfp1 | -2.231 | 3.38E-04 |
| Defb4 | -1.356 | 5.74E-04 |
| Crnn | -1.162 | 2.38E-05 |
| Slco1a5 | -1.136 | 2.43E-05 |
| Dapl1 | -0.839 | 3.07E-04 |
| Pglyrp1 | -0.813 | 5.59E-05 |
| Gm48362 | -0.811 | 4.03E-04 |
| Il36g | -0.776 | 8.46E-04 |
| Tmem45a | -0.775 | 2.48E-06 |
| Alox12 | -0.751 | 2.37E-04 |
| Sptssb | -0.642 | 6.91E-04 |
| Ehf | -0.618 | 5.74E-04 |
| Tfap2a | -0.566 | 5.95E-04 |
| Lgalsl | -0.530 | 1.54E-05 |
| Ly6d | -0.528 | 1.70E-04 |

| DEHP 10uM \| top15 upregulated | | |
|---|---|---|
| gene | log2 fold change | adj pvalue |
| Gkn1 | 2.594 | 6.30E-03 |
| Lypd8l | 2.028 | 2.51E-03 |
| Gm10925 | 0.895 | 3.95E-03 |
| Hormad1 | 0.394 | 5.40E-03 |
| Slc7a3 | 0.374 | 6.30E-03 |
| Spp1 | 0.344 | 5.56E-03 |
| Mt2 | 0.315 | 4.18E-05 |
| Cox7b | 0.202 | 5.56E-03 |
| Lactb2 | 0.189 | 9.10E-03 |
| Acsl4 | 0.161 | 3.39E-03 |
| Kif5b | 0.110 | 6.94E-03 |

| DEHP 10uM \| top15 downregulated | | |
|---|---|---|
| gene | log2 fold change | adj pvalue |
| mt-Atp6 | -0.816 | 1.19E-06 |
| Fam163a | -0.536 | 2.82E-04 |
| Pdzd2 | -0.411 | 1.91E-07 |
| Rgma | -0.295 | 3.88E-06 |
| Zfhx3 | -0.287 | 2.04E-03 |
| Plce1 | -0.245 | 2.75E-03 |
| Atn1 | -0.210 | 2.00E-03 |
| Kctd17 | -0.209 | 2.75E-03 |
| Zmiz1 | -0.202 | 5.19E-04 |
| Prr12 | -0.195 | 3.88E-06 |
| Phf2 | -0.184 | 2.04E-03 |
| Adnp | -0.175 | 2.51E-03 |
| Dvl3 | -0.171 | 2.66E-03 |
| Wdfy3 | -0.153 | 2.51E-03 |
| Khsrp | -0.147 | 2.03E-03 |

| PCB153 10uM \| top15 upregulated | | |
|---|---|---|
| gene | log2 fold change | adj pvalue |
| Rps6-ps4 | 11.994 | 1.78E-03 |
| Actc1 | 0.902 | 9.98E-03 |

| DMMP 1nM \| top15 downregulated | | |
|---|---|---|
| gene | log2 fold change | adj pvalue |
| Bfsp1 | -1.086 | 8.36E-04 |
| Rbp2 | -1.049 | 5.80E-04 |
| Krt16 | -0.951 | 8.42E-06 |
| Il36g | -0.950 | 1.54E-04 |
| Il36a | -0.933 | 1.34E-04 |
| Fam25c | -0.855 | 1.25E-04 |
| Gm5478 | -0.851 | 1.60E-05 |
| Krt6b | -0.843 | 8.15E-06 |
| Dsc3 | -0.830 | 2.24E-03 |
| Urah | -0.640 | 1.12E-03 |
| Tmprss11d | -0.562 | 5.80E-04 |
| Fabp5 | -0.535 | 5.80E-04 |
| Tlcd3a | -0.510 | 5.80E-04 |
| Tgm1 | -0.421 | 2.13E-03 |
| Gpx1 | -0.257 | 1.78E-03 |

**Table S10 – Top 15 upregulated and downregulated proteins after 10 days exposure to 10 µM BaP, PCB153, and DEHP.**

| 10 days BaP 10uM \| top15 upregulated | | |
|---|---|---|
| Protein | log2 fold change | adj pvalue |
| Nol8 | 1,32 | 7,88E-03 |

| 10 days BaP 10uM \| top15 downregulated | | |
|---|---|---|
| Protein | log2 fold change | adj pvalue |
| Mylk | -3,85 | 2,83E-05 |
| Tmem97 | -3,20 | 2,03E-05 |
| Nolc1 | -2,17 | 9,26E-03 |
| Pbx1 | -2,17 | 9,62E-03 |
| Vkorc1 | -1,90 | 7,96E-03 |
| Mical1 | -1,57 | 1,73E-03 |
| Eed | -1,10 | 5,21E-03 |

| 10 days BaP 1nM \| top15 upregulated | | |
|---|---|---|
| Protein | log2 fold change | adj pvalue |

| 10 days BaP 1nM \| top15 downregulated | | |
|---|---|---|
| Protein | log2 fold change | adj pvalue |
| Tmem97 | -2,91 | 1,04E-03 |
| Tspan9 | -2,00 | 1,41E-03 |
| Cdc73 | -1,80 | 4,18E-03 |
| Ppp1r14b | -1,46 | 5,73E-03 |
| Synj2bp | -1,17 | 7,37E-03 |

| 10 days PCB153 10uM \| top15 upregulated | | |
|---|---|---|
| Protein | log2 fold change | adj pvalue |
| Nol8 | 1,93 | 6,44E-03 |
| Bag3 | 1,89 | 1,47E-03 |
| Rpap3 | 1,46 | 3,66E-03 |

| 10 days PCB153 10uM \| top15 downregulated | | |
|---|---|---|
| Protein | log2 fold change | adj pvalue |
| Cdc73 | -3,38 | 1,80E-04 |
| Eif2b5 | -2,08 | 2,60E-03 |
| Pum1 | -1,87 | 3,42E-03 |
| Mettl16 | -1,72 | 4,88E-03 |
| Tmem97 | -1,70 | 2,71E-03 |
| Eed | -1,31 | 5,97E-03 |

| 10 days PCB153 1nM \| top15 upregulated | | |
|---|---|---|
| Protein | log2 fold change | adj pvalue |
| Aqp3 | 3,99 | 8,63E-03 |
| Ptgs2 | 1,56 | 8,01E-03 |

| 10 days PCB153 1nM \| top15 downregulated | | |
|---|---|---|
| Protein | log2 fold change | adj pvalue |
| Cdc73 | -2,27 | 2,15E-03 |
| Nolc1 | -2,23 | 7,42E-04 |
| Nol6 | -2,21 | 1,63E-05 |
| Agfg1 | -2,07 | 8,17E-03 |
| Gnai1 | -1,70 | 7,79E-03 |
| Mettl16 | -1,61 | 2,26E-03 |
| Ddx41 | -1,32 | 7,20E-03 |
| Fdft1 | -0,76 | 1,90E-03 |

| 10 days DEHP 10uM \| top15 upregulated | | |
|---|---|---|
| Protein | log2 fold change | adj pvalue |
| Cdc40 | 1,31 | 8,91E-03 |

| 10 days DEHP 10uM \| top15 downregulated | | |
|---|---|---|
| Protein | log2 fold change | adj pvalue |
| Fbxo7 | -1,01 | 5,06E-03 |

| 10 days DEHP 1nM \| top15 upregulated | | |
|---|---|---|
| Protein | log2 fold change | adj pvalue |
| Afg3l2 | 2,17 | 5,92E-04 |
| Pglyrp1 | 1,77 | 8,76E-03 |
| Mtch1 | 1,59 | 5,67E-03 |
| Ado | 1,49 | 4,78E-03 |
| Abcb10 | 1,46 | 2,32E-03 |
| Tmem97 | 1,43 | 9,20E-03 |
| Abcc4 | 1,21 | 8,88E-03 |
| Ly6g6c | 1,16 | 5,25E-03 |
| Wbp11 | 0,99 | 7,22E-03 |
| Clcc1 | 0,91 | 4,57E-04 |
| Enpp5 | 0,76 | 7,94E-03 |
| Ceacam1 | 0,73 | 6,89E-03 |
| Fnta | 0,67 | 3,64E-03 |
| Nipsnap3b | 0,58 | 6,09E-03 |
| Slc7a6 | 0,53 | 1,55E-03 |

| 10 days DEHP 1nM \| top15 downregulated | | |
|---|---|---|
| Protein | log2 fold change | adj pvalue |
| Sdf2l1 | -3,14 | 5,33E-04 |
| Snap23 | -2,45 | 9,30E-03 |
| Hars1 | -2,30 | 1,06E-03 |
| Rps27l | -2,09 | 3,17E-03 |
| Acadvl | -1,78 | 8,11E-03 |
| Glrx5 | -1,60 | 4,53E-03 |
| Ndufv1 | -1,54 | 1,17E-03 |
| H1-5 | -1,38 | 2,27E-03 |
| Rbm15 | -1,32 | 2,11E-04 |
| Atp11a | -1,29 | 3,57E-03 |
| Lrrc40 | -1,15 | 7,67E-03 |
| Enkur | -1,07 | 5,02E-03 |
| Camk1d | -0,98 | 7,16E-03 |
| Khdrbs1 | -0,88 | 7,35E-03 |

**Table S11. Thyroid hormone concentrations**. Targeted quantification of the thyroid hormones 3,3'-T2, T3 and T4 (pmol/mL) in medium was performed upon 10 days-exposure to BaP, PCB153, and DEHP (1 nM and 10 µM). Exposure to the reference compound Methimazol (MMI, 10 µM) and the unexposed control (DMSO) were included in both batches. The average ± standard deviation and the number of replicates included in the average (n) is shown.

| | **T2** (pmol/mL) | **T3** (pmol/mL) | **T4** (pmol/mL) |
|---|---|---|---|
| DMSO – Batch 1 | 14.4 ± 1.1 *(n=4)* | 30.1 ± 4.7 *(n=4)* | 85.6 ± 33.5 *(n=6)* |
| DMSO – Batch 2 | 13.7 ± 1.8 *(n=7)* | 22.8 ± 3.4 *(n=9)* | 84.8 ± 46.0 *(n=7)* |
| MMI – Batch 1 | 12.7 ± 1.0 *(n=6)* | 8.2 ± 1.1 *(n=8)* | 62.6 ± 27.3 *(n=7)* |
| MMI – Batch 2 | 12.2 ± 0.5 *(n=9)* | 9.4 ± 2.7 *(n=10)* | 39.5 ± 18.5 *(n=10)* |
| BaP- 1 nM | 16.4 ± 2.0 *(n=9)* | 32.3 ± 8.1 *(n=8)* | 65.1 ± 29.7 *(n=5)* |
| BaP – 10 µM | 16.8 ± 1.5 *(n=9)* | 37.1 ± 2.9 *(n=9)* | 62.8 ± 14.3 *(n=8)* |
| DEHP- 1 nM | 15.3 ± 3.1 *(n=8)* | 17.3 ± 4.2 *(n=9)* | 138.8 ± 24.6 *(n=4)* |
| DEHP – 10 µM | 18.6 ± 3.4 *(n=7)* | 21.3 ± 5.5 *(n=6)* | 48.8 ± 8.4 *(n=3)* |
| PCB153- 1 nM | 20.3 ± 8.3 *(n=7)* | 38.6 ± 12.1 *(n=7)* | 68.0 ± 39.6 *(n=3)* |
| PCB153 – 10 µM | 19.2 ± 4.6 *(n=8)* | 43.1 ± 13.0 *(n=8)* | 77.2 ± 50.5 *(n=6)* |
